# Metabolism of glucose and trehalose by cyclic pentose phosphate pathway is essential for effective immune response in *Drosophila*

**DOI:** 10.1101/2023.08.17.553657

**Authors:** Michalina Kazek, Lenka Chodáková, Katharina Lehr, Lukáš Strych, Pavla Nedbalová, Ellen McMullen, Adam Bajgar, Petr Šimek, Martin Moos, Tomáš Doležal

## Abstract

Activation of immune cells requires the remodeling of cell metabolism in order to support immune function. We study these metabolic changes through the infection of *Drosophila* larvae by parasitoid wasp. Neutralization of the parasitoid egg involves the differentiation of lamellocytes, which encapsulate the egg. A melanization cascade is initiated, producing toxic molecules to destroy the egg; meanwhile the capsule created also protects the host from the toxic reaction. We combine transcriptomics and metabolomics, including 13C-labeled glucose and trehalose tracing, as well as genetic manipulation of sugar metabolism to study changes in metabolism, specifically in *Drosophila* hemocytes. We found that hemocytes increase the expression of several carbohydrate transporters, and accordingly uptake of sugar during infection. These carbohydrates are metabolized by increased glycolysis, associated with lactate production, and cyclic pentose phosphate pathway (PPP), in which glucose-6-phosphate is re-oxidized to maximize NADPH production. Oxidative PPP is required for lamellocyte differentiation and resistance, as is systemic trehalose metabolism. In addition, fully differentiated lamellocytes use a cytoplasmic form of trehalase to cleave trehalose to glucose, within the cell, to fuel cyclic PPP. Intracellular trehalose metabolism is not required for resistance, but may be associated with host protection from its own toxic response. Thus, our results suggest that sugar metabolism within immune cells, and specifically in cyclic PPP, may be important for not only fighting infection, but also for protecting the host from its own immune response and ensuring sufficient fitness of the survivor.

## Introduction

Quiescent immune cells require nutrients to efficiently carry out their basic functions. When activated, however, they must rapidly undertake many more functions, facilitated by the rapid generation of energy and biosynthetic intermediates [1]. Most types of mammalian immune cells and, as we have recently shown, insect immune cells become more dependent on glucose and have an increased rate of glycolysis during infection [1,2]. This allows for the rapid generation of ATP and branch into other metabolic pathways, for example the pentose phosphate pathway (PPP) [3]. The high rate of energy production and large amounts of intermediates make immune cells highly nutrient demanding during their activation. We have previously shown that *Drosophila melanogaster* larval immune cells (hemocytes) increase their consumption of total systemic glucose from 10% to 27% during the immune response to a parasitoid wasp [4]. Hemocytes release adenosine to suppress carbohydrate consumption by non-immune tissues to ensure their own supply, which is critical for an effective immune response [4].

Trehalose is the primary carbohydrate of insects; when cleaved by trehalase (Treh) it provides a rapid source of glucose [5]. Unlike glucose, trehalose is a non-reducing sugar and thus the hemolymph of *Drosophila* larvae contains a tenfold higher concentration of trehalose than glucose [4,6]. This serves as a buffer for glucose homeostasis to ensure robust and stable development [7]. We previously found that hemocytes strongly upregulate expression of the trehalose transporter Tret1-1 and Treh during immune response [4]. This was later confirmed by single cell transcriptomics studies [8,9], which showed lamellocyte-specific expression of Tret1-1 and Treh. These results suggest an important role for trehalose during immune response, which has also been shown in house flies [10].

We use a model of infection of *Drosophila* larvae by the parasitoid wasp *Leptopilina boulardi* where the wasp injects its egg during larval development [11]. Within a few hours, the egg is recognized by circulating hemocytes (plasmatocytes), which induce an immune response. Sessile hemocytes enter the circulation, some attach to the egg while others differentiate into specialized large flat cells called lamellocytes, which later encapsulate the egg. The melanization cascade is initiated in the forming capsule [12] and within approximately 48 hours the parasitoid egg is destroyed. If the immune response is not fast or efficient, the parasitoid larva emerges from the egg and eventually consumes the developing fly in the pupa [4].

Melanization is associated with both production and scavenging of toxic substances [12]. Encapsulation/melanization thus serves the dual purpose of producing and concentrating toxic substances inside the capsule to kill the parasitoid, while protecting the host from escaping toxic radicals. Therefore, there are metabolic requirements associated with lamellocyte differentiation (global changes in gene expression, cytoskeletal and membrane rearrangements, etc.), production of toxic molecules (e.g., reactive oxygen species, ROS), and host protection mechanisms (e.g., antioxidant production). Metabolic reprogramming for the individual tasks involved in killing pathogens and protecting the host is essential for the survival of the infected larva and the future fitness of the surviving organism. However, changes in the actual metabolism of larval hemocytes during immune response to wasp infection have not yet been investigated.

One of the most important branches from increased glycolysis during immune response is the PPP [3], generating NADPH and pentoses as precursors for nucleotides and coenzymes [13]. NADPH is essential in activated immune cells for lipid biosynthesis, for the ROS production needed to fight pathogens [14], as well as for the production of antioxidants, such as glutathione, to protect the host from excessive ROS exposure [15]. Cells can dramatically increase NADPH production through cyclic PPP, which repeatedly oxidizes glucose-6-phosphate (G6P), as recently demonstrated in neutrophils [3].

Immune cells have privileged access to nutrients during immune response [16]. We hypothesize that hemocytes can secure prioritized allocation of carbohydrates using trehalose, i.e. by expressing Tret1-1 along with Treh. To test this hypothesis, we employed 13C stable isotope tracing [17] to analyze metabolic changes in hemocytes as well as genetically manipulating trehalose metabolism to investigate its role during immune response to parasitoid wasps. We found that hemocytes express several carbohydrate transporters, some of which are dramatically up-regulated during infection. 13C tracing experiments show that activated hemocytes increase the uptake and metabolism of glucose and trehalose via glycolysis and in particular the cyclic PPP, which is essential for lamellocyte production and resistance. Systemic trehalose metabolism is important for an effective immune response, but trehalose itself is only metabolized in fully differentiated lamellocytes, which is not necessary for resistance but instead appears to be important for lamellocyte-mediated host protection.

## Results

### Activated hemocytes increase the expression of carbohydrate transporters and trehalase

*Drosophila* hemocytes increase their uptake of carbohydrates during parasitoid wasp infection [4], we have therefore analyzed the expression of carbohydrate transporters in immune cells. The SLC2 family of hexose sugar transporters in *Drosophila* comprises 31 genes (S1 File; FlyBase ID: FBgg0000691), most of which have not been functionally characterized. Our bulk transcriptomic analysis showed that the expression of ten of these genes is greater than three transcripts per million (TPM) in hemocytes (S1 file and S1 Table) and four of these genes are expressed in hemocytes during infection – *Tret1-1, CG4607, sut1* and *CG1208* (Fig 1A). Tret1-1 has been functionally characterized as glucose and trehalose transporter [18] and CG4607 appears to be involved in lysosomal glucose metabolism [19]. Both *Tret1-1* and *CG4607* are weakly expressed in hemocytes in the uninfected state, but their expression strongly increases during infection (Fig 1A, S1 file). Based on scRNAseq [9], both are expressed exclusively in lamellocytes (S1 file). *sut1* is highly expressed in most hemocyte types in both uninfected and infected state (Fig 1A and S1 file). MFS3, which belongs to the SLC17 family of organic anion transporters, has also been shown to transport glucose and trehalose [20], it’s strongest expression in hemocytes is in the uninfected state, and decreases during infection (Fig 1A and S1 file). The functionally uncharacterized CG1208 shows the strongest increase in expression upon infection, mainly in lamellocytes (Fig 1A and S1 file). Thus, MFS3 and sut1 appear to provide basal carbohydrate transport in most hemocyte types in the uninfected state, whereas Tret1-1 and presumably CG4607 and CG1208 provide carbohydrate transport in lamellocytes during infection.

**Fig 1.**
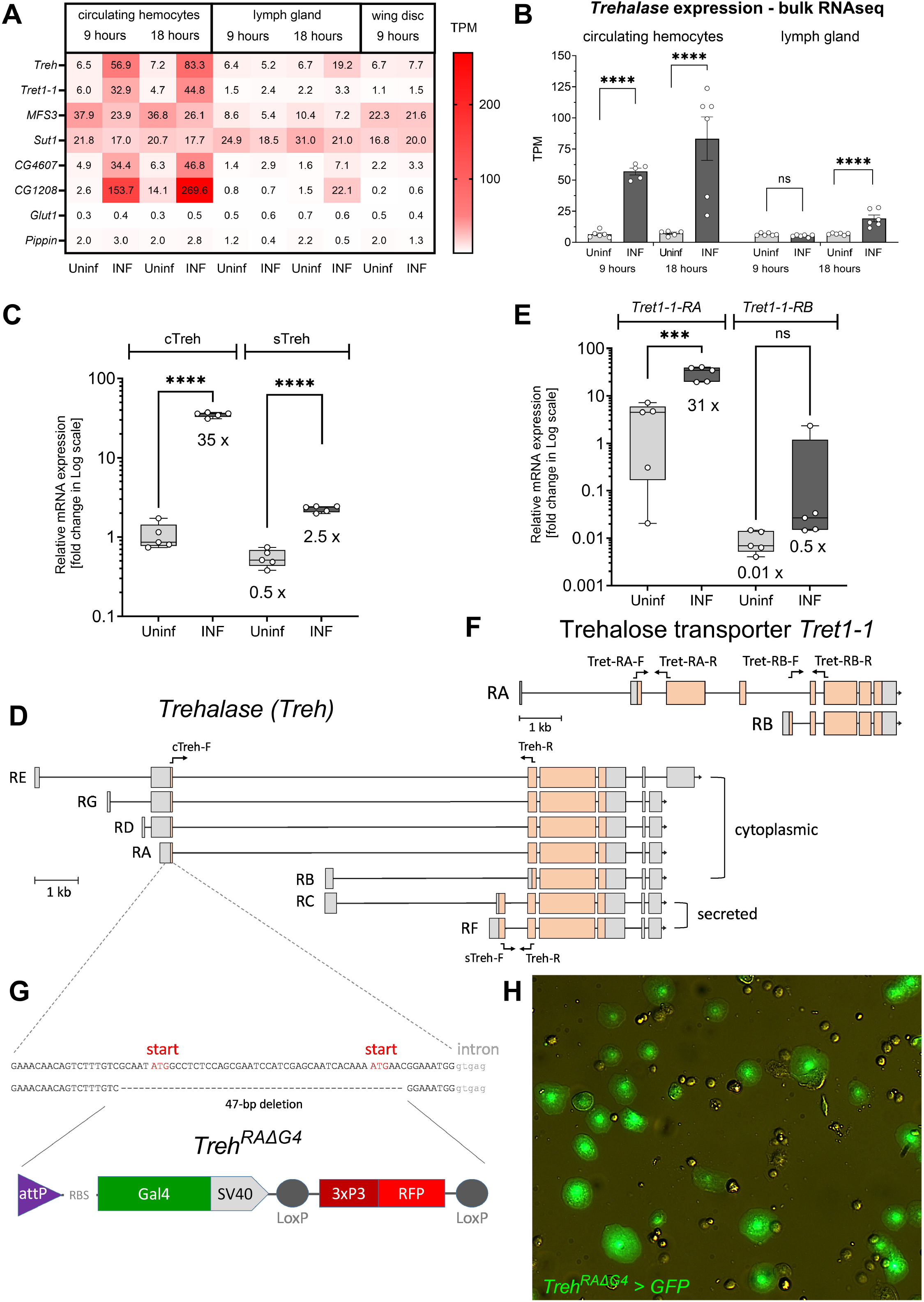
Analysis of carbohydrate metabolism gene expression. (A) Heat map of expression (bulk RNAseq) of selected transporters and trehalase in circulating hemocytes, lymph gland and wing disc from uninfected (Uninf) and infected (INF) third-instar larvae collected 9 and 18 hours after the start of infection = 0 hour = 72 hours after egg laying. Values given in each cell are transcripts per million (TPM). (B) Expression of the *trehalase* (*Treh*) gene (bulk RNAseq) in circulating hemocytes and lymph gland. Each dot represents a biological replicate in TPM, bars represent mean ± SEM. Samples were analyzed using DESeq2 in Geneious Prime (S2 Table), **** shows adjusted P < 0.0001, ns = not significant. (C) Transcript-specific analysis of *Treh* expression by RT-qPCR 18 hours after the start of infection. cTreh, represents cytoplasmic trehalase (primers cTreh-F and Treh-R shown in (D)), increases 35-fold after infection. sTreh represents secreted trehalase (primers sTreh-F and Treh-R shown in (D)). Box and whiskers plots (median, 75th and 25th percentiles, and maximum/minimum) show fold change compared to uninfected cTreh samples (expression levels were normalized by *RpL32* expression in each sample), each dot represents a biological replicate. An unpaired two-tailed t test was used to compare uninfected and infected samples; ****P < 0.0001. (D) Map of the *Treh* gene with individual transcripts (RA-RG). Lines show introns, boxes show exons with coding sequence in orange. Labeled arrows show primers used for RT-qPCR expression analysis. (E) Transcript specific analysis of trehalose transporter *Tret1-1* by RT-qPCR 18 hours after the start of infection. *Tret1-1-RA* (primers Tret-RA-F and Tret-RA-R shown in (F)) increases 31-fold after infection. *Tret1-1-RB* (primers Tret-RB-F and Tret-RB-R shown in (F)). Box and whiskers plots (median, 75th and 25th percentile, and maximum/minimum) show fold change compared to uninfected *Tret1-1-RA* samples (expression levels normalized by *RpL32* expression in each sample), each dot represents a biological replicate. Unpaired two-tailed t test was used to compare uninfected and infected samples; \*\*\**P* < 0.001, ns = not significant. (F) Map of the *Tret1-1* gene with RA and RB transcripts. Lines show introns, boxes show exons with coding sequence in orange. Labeled arrows show primers used for RT-qPCR expression analysis. (G) Schematic representation of a Gal4 knock-in into the first exon of *Treh-RA*, creating a 47-base deletion that removes both cTreh start codons, replaced by a cassette containing the Gal4 coding sequence and an RFP marker, expressed downstream of the P3 regulatory sequence in the fly eye. The resulting fly strain is *Treh[RAΔG4]*. (H) *Treh[RAΔG4*], expressing Gal4 in the cTreh expression pattern, drives UAS-GFP expression in differentiated lamellocytes (green) but not in plasmatocytes. Differential interference contrast (DIC) combined with fluorescence microscopy using 20x objective.

While *sut1* is moderately expressed in the lymph gland, we did not detect increased expression of transporters other than *CG1208* at 18 hours post infection (Fig 1A). MFS3 and Sut1 also appear to be major carbohydrate transporters in the wing disc, but there are no changes in carbohydrate transporters expression during infection (Fig 1A).

Interestingly, along with the trehalose transporter Tret1-1, expression of the enzyme Treh, which converts trehalose into two glucose molecules, is also strongly increased in hemocytes upon infection (Fig 1A and 1B), this increase is primarily due to expression in lamellocytes (S1 File). Treh exists in two forms, cytoplasmic (cTreh) and secreted (sTreh; Fig 1D) [21], we therefore used transcript-specific qPCR to determine which form is expressed in hemocytes. cTreh transcripts (*Treh-RA, -RD, -RG, -RE*) increase 35-fold in hemocytes upon infection (Fig 1C). The transcripts for the sTreh form (*Treh-RC* and *-RF*) also increase during infection, but only slightly more than the basal level of cTreh. Therefore, substantially more cTreh than sTreh is expressed in hemocytes during infection. There are also two transcriptional variants of Tret1-1 (Fig 1F), with *Tret1-1-RA* increasing 31-fold upon infection, whereas there is no increase in the *Tret1-1-RB* variant (Fig 1E).

We generated a cTreh-specific mutant, *Treh[RAΔGal4]*, by replacing 47 bases including the first two start codons with the Gal4 coding sequence, which can also serve as an expression reporter (Fig 1G). By crossing this line with flies carrying *UAS-GFP* (Fig 1H and S1 Fig), we verified lamellocyte-specific expression of cTreh among hemocytes. cTreh is also expressed in other larval tissues, such as imaginal discs (S1 Fig) and brain (S2 Fig), with and without infection.

Our expression analysis showed that hemocytes express multiple carbohydrate transporters, some of which also dramatically increase expression during infection. This is consistent with our previous results that showed increased sugar consumption under these conditions. The combination of dramatically increased expression of trehalose transporters together with a cytoplasmic form of Treh exclusively in lamellocytes suggests the importance of trehalose metabolism in these cells.

### Activated hemocytes increase uptake and metabolism of glucose and trehalose via glycolysis and pentose phosphate pathway

Our bulk transcriptomic analysis shows that all enzymes associated with glycolysis and PPP are strongly expressed in hemocytes in both the uninfected and infected state (Fig 2, S2 file and S2 Table). The expression of specific genes in each pathway corresponds well with scRNAseq analyses (S2 file; [8,9]). Same glycolytic and PPP genes are also similarly expressed in the lymph gland and wing disc, both in uninfected and infected state (S2 file and S2 Table). Combining our bulk transcriptomics results with scRNAseq reveals a tendency for a slight decrease in the expression of glycolytic genes in most prohemocytes and plasmatocyte-like cells during infection, while expression shifts towards lamellocytes and crystal cells (S2 file). Expression of PPP genes generally does not change during infection. Phosphofructokinase (Pfk) shows the lowest expression of all glycolytic genes and is further reduced in all hemocyte types during infection, suggesting a shift away from glycolysis to PPP and back to glycolysis at the glyceraldehyde-3-phosphate level. Overall, the expression analysis does not indicate specific changes in these metabolic pathways in either hemocyte types, or during infection.

**Fig 2.**
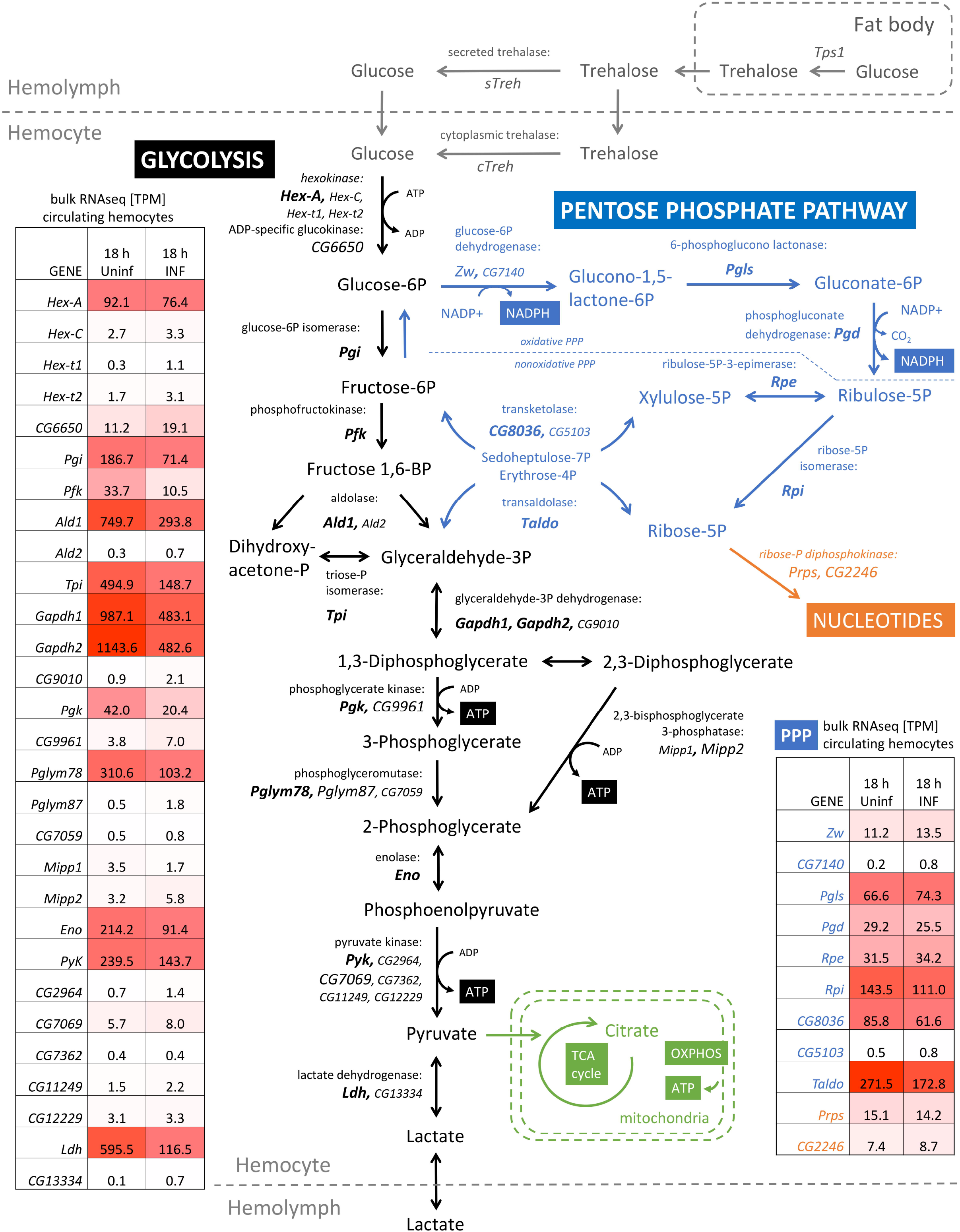
Expression of glycolytic and pentose phosphate pathway enzymes in hemocytes. Diagram showing metabolic pathways; metabolites and enzymes are abbreviated with *Drosophila* gene names; strongly expressed genes in hemocytes are in large font in bold, moderately expressed genes in large font, and insignificantly expressed genes in small font. Trehalose metabolism is shown in grey, glycolysis in black, pentose phosphate pathway in blue, purine metabolism in orange and mitochondrial metabolism in green. Tables show glycolytic and PPP genes expression based on bulk RNAseq in transcripts per million (TPM) in circulating hemocytes at 18 hours after start of infection from the uninfected (18 h Uninf) and infected (18 h INF) larvae; the red shading of the table highlights the strength of expression -white the weakest, red the strongest.

In order to monitor changes in the metabolism of hemocytes during infection, we used two approaches involving the tracing of metabolites labeled with stable 13C_6_ isotopes. In the first, we fed 13C-labeled glucose (D-glucose-^13^C_6_) to larvae at 16 hours post infection for 6 hours before collecting hemocytes at 22 hours post infection. After 6 hours of feeding, 22% of the glucose in the hemolymph was labeled (Fig 3A). Some of the labeled glucose was also converted to trehalose in the fat body, resulting in 16% of the circulating trehalose being labeled (at one of the two glucose units; Fig 3B). Both glucose and trehalose in the hemolymph increase during infection, and the increase is mainly due to the unlabeled sugars (Fig 3A and 3B), suggesting they come from stores, which is consistent with our previous results [4]. Hemocytes can directly take up labeled dietary glucose or convert labeled trehalose to glucose. There is five times more glucose in the infection-activated hemocytes than in the control (Fig 3C). The increased 13C-labeled glucose shows that hemocytes increase uptake of sugars upon infection (Fig 3C), again in agreement with our previous results [4].

**Fig 3.**
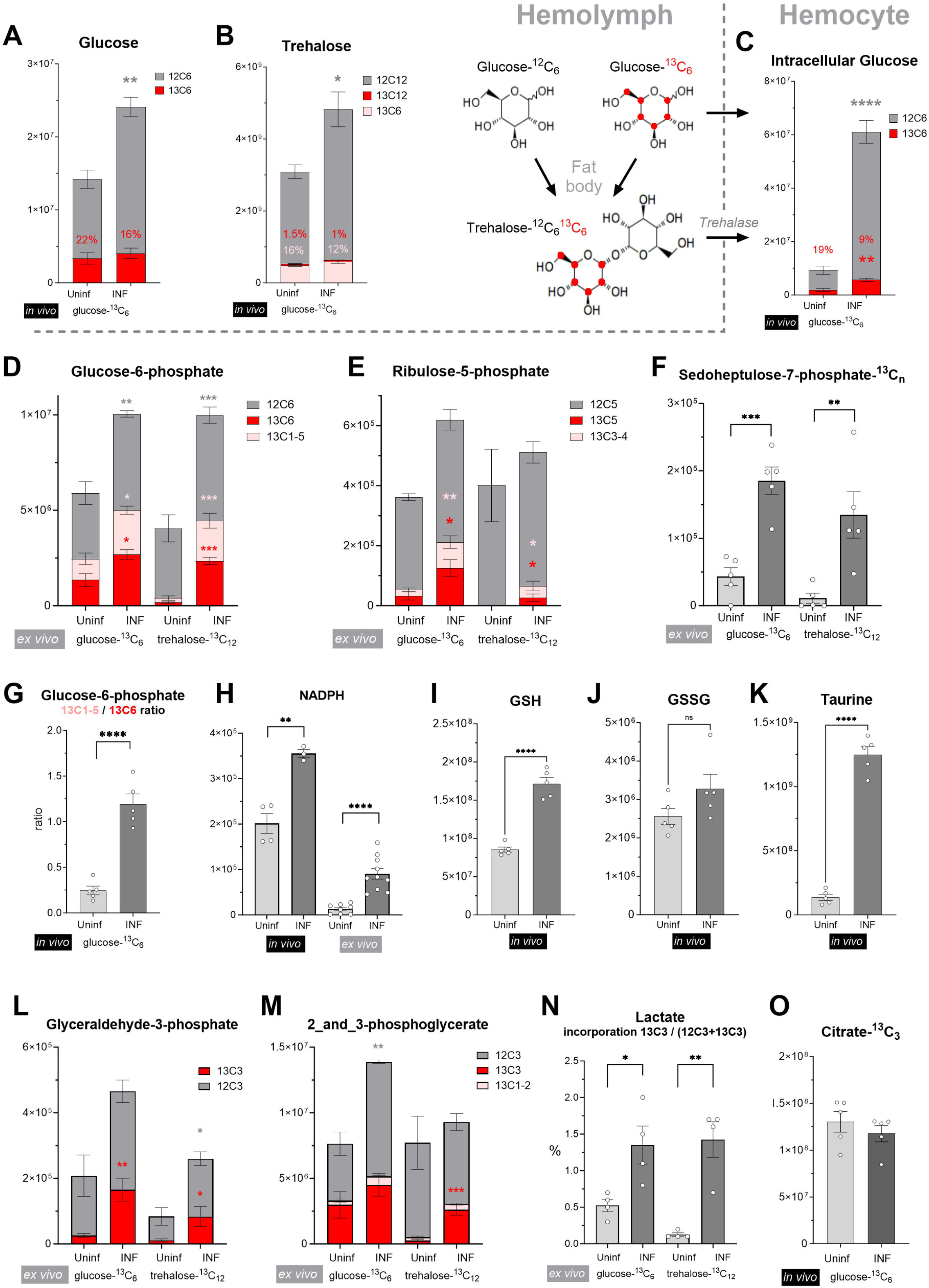
Analysis of hemocyte metabolism by stable 13C isotope tracing. The bars show the mean metabolite amount - unlabeled form, or labeled with 13C stable isotope, or both, stacked in one bar - expressed by the normalized peak area, unless otherwise indicated in the graph title and at the Y-axis. Graphs labeled “*in vivo*” in black box - larvae were fed labeled D-glucose-^13^C_6_. Graphs labeled “*ex vivo*” in gray box - hemocytes were incubated *ex vivo* with either labeled D-glucose-^13^C_6_ or α,α−trehalose- ^13^C_12_. Samples were obtained from hemocytes from uninfected (Uninf) or infected (INF) larvae. (A,B) Circulating glucose and trehalose levels in hemolymph and (C) intracellular glucose levels in hemocytes after *in vivo* feeding – (A,C) unlabeled (grey), fully labeled (^13^C_6_ - red), (B) unlabeled (grey), fully labeled (^13^C_12_ - red), partially labeled (^13^C_6_ – pink). The percentages above the columns express the fraction of the labelled from the total amount. (D,E) Intracellular glucose-6-phosphate and ribulose-5- phosphate levels in hemocytes upon *ex vivo* incubation with labeled glucose (left bars) or trehalose (right bars). The stacked bars combine unlabeled (gray), fully labeled (^13^C_6_ or ^13^C_5_ in red), and partially labeled (one to five out of the six carbons labeled in the molecule, in pink). (F) Levels of 13C-labeled sedoheptulose-7-phosphate (all forms with any labeled carbon) in hemocytes upon *ex vivo* incubation. (G) Ratio of normalized peak areas of partially labeled relative to fully labeled glucose-6-phosphate after *in vivo* feeding. (H-K) Levels of unlabeled NADPH, GSH, GSSG, and taurine in hemocytes *ex vivo* or *in vivo* (indicated below the bars). (L,M) Levels of glyceraldehyde-3-phosphate and combined 2-phosphoglycerate with 3-phosphoglycerate in hemocytes upon *ex vivo* incubation with labeled glucose (left bars) or trehalose (right bars). Stacked bars show unlabeled (gray), fully labeled (^13^C_3_ in red), and partially labeled (one and two out of the three carbons labeled in the molecule, in pink). (N) Fraction (%) of 13C-labeled intracellular lactate in total lactate upon *ex vivo* incubation with labeled glucose (left bars) and trehalose (right bars). (O) Levels of 13C-labeled citrate in hemocytes after *in vivo* feeding. Levels from infected larvae were corrected 1.6-fold based on labeling of the preceding pyruvate. (A-O) The sample from infected larvae was compared with that from uninfected larvae using unpaired t test or ordinary two-way ANOVA with multiple comparisons. Asterisks indicate p value (* P < 0.05, ** P < 0.01, *** P < 0.001, **** P < 0.0001) and are either above the bar in the corresponding color of the bar they compare within the stacked bars, or in black for a simple comparison. A bar without asterisks indicates a non-significant difference. Error bars represent ± SEM.

A direct comparison of the labeling of downstream metabolites between uninfected and infected larvae is complicated by the different percentage of labeled glucose within the hemocytes - 19% versus only 9% (Fig 3C) - which affects our ability to compare entry of labeled glucose into downstream pathways. To obtain a comparable amount of labeled glucose entering hemocytes, we used a second approach in which we incubated hemocytes in 100% D-glucose-^13^C_6_ *ex vivo*. This approach also allowed us to use labeled trehalose (α,α−trehalose-^13^C_12_) as a source and distinguish its metabolism from glucose. We first collected hemocytes by bleeding larvae 22 hours post infection and immediately incubated them *ex vivo* for 40 minutes in a solution containing 100% 13C-labeled glucose or trehalose.

By examining only one time point and working with a heterogeneous population of hemocytes *in vivo* for our 13C-labeled carbohydrates experiments we are unable to determine metabolic fluxes, however, we can still observe metabolic changes. Several interesting aspects of metabolism in activated hemocytes are revealed by glucose-6-phosphate (G6P) labeling. Firstly, G6P increases in activated hemocytes, including newly formed, i.e. labeled, G6P (Fig 3D), demonstrating that glucose metabolism is indeed enhanced during infection. Secondly, hemocytes of uninfected larvae metabolize glucose but almost no trehalose, which they metabolize substantially only during infection (e.g., Fig 3D, 3E and 3F). This is consistent with the fact that the cytoplasmic form of *Treh* is expressed only in lamellocytes. Therefore, using labeled trehalose as a source, most of the 13C incorporation can be attributed to lamellocyte metabolism. Additionally, fully labeled G6P-^13^C_6_ (red in Fig 3D) shows increased levels upon infection, but we also detected an increase in partially labeled G6P-^13^C_1-5_ (pink in Fig 3D). Partially labeled G6P can be generated from fully labeled glucose by cyclic PPP, in which pentoses formed by oxidative PPP are converted back to G6P by the action of transketolase/transaldolase in the non-oxidative PPP and the reverse action of glucose-6-phosphate isomerase (Fig 2 and S3; [22,23]). Since labeled pentoses represent a minor fraction (Fig 3E), they almost always combine with unlabeled pentoses to form partially labeled G6P (S3 Fig). This is also the case when labeled trehalose is used as the source (Fig 3D), demonstrating that cyclic PPP is active in lamellocytes. Increased labeling of ribulose-5-phosphate (Fig 3E), a product of PPP, and sedoheptulose-7-phosphate (Fig 3F), an intermediate of transketolase/transaldolase activity, further supports increased PPP in hemocytes during infection. The intensification of cyclic PPP is also evidenced by the *in vivo* experiment mentioned above in which 13C-labeled glucose is fed to larvae. Although we cannot directly compare uninfected and infected larvae for the reasons mentioned above, we can compare the ratio of partially labeled G6P to fully labeled G6P. That is, how many molecules were converted back to G6P relative to the number of glucose molecules that entered the PPP. Whereas, in uninfected larvae there is only one partially labeled G6P molecule for every four fully labeled G6P molecules, in infected larvae this ratio increases to more than 1:1 (Fig 3G), indicating an intensified cyclic PPP. Oxidative PPP produces NADPH, which increases in hemocytes upon infection both *in vivo* and *ex vivo* (Fig 3H). One of the roles of NADPH is to reduce antioxidants such as glutathione, and we observed increased levels of GSH (Fig 3I), the reduced form of glutathione, in hemocytes from infected larvae, while the oxidized form GSSG is at comparable levels (Fig 3J). Flies use the thioredoxin system instead of glutathione reductase to reduce GSSG [24], and hemocytes highly express thioredoxin reductase Trxr1 and thioredoxin Trx-2 with greater expression in lamellocytes (S4 Fig). Trx2 is also a substrate for peroxiredoxins [25], which are also highly expressed in hemocytes (S4 Fig). Besides GSH, infection elevates another antioxidant, taurine, in hemocytes (Fig 3K).

Comparison of fractions with different numbers of labeled carbons in G6P and ribulose-5-phosphate reveals further insights (S5 Fig and S6 Fig). Before increasing amounts of labeled xylulose-5-phosphate and ribose-5-phosphate enter the transketolase/transaldolase conversion of pentoses to hexoses, the most common product is G6Pm+3 (pink in S3 Fig), which is formed by coupling labeled glyceraldehyde-3-phosphate with unlabeled sedoheptulose-7-phosphate. This is evident after a short *ex vivo* incubation, including incubation with trehalose, which is metabolized in lamellocytes (S5 Fig). Once the labeled pentoses enter the transketolase/transaldolase conversion to hexoses, G6Pm+2 begins to dominate (yellow in S3 Fig). We observe this after prolonged exposure to 13C in vivo (S6 Fig). Ribulose-5-phosphate and ribose-5-phosphate can be formed both in oxidative PPP (thus generating NADPH) and in non-oxidative PPP, in which the opposite direction of transketolase/transaldolase activity generates pentoses without NADPH (scheme in Fig 2 and S7). If pentoses were formed predominantly by oxidative PPP, their partial/complete labeling pattern should match that of G6P (in some form minus one labeled carbon due to decarboxylation). This is the case in the *ex vivo* experiment, especially during infection (S5 Fig). However, our *in vivo* experiment showed a different pattern for both ribulose-5-phosphate and ribose-5-phosphate, where we detected much more m+1, m+2, and m+3 than m+5 (S6 Fig). This pattern is more consistent with a combination of fructose-6-phosphate and glyceraldehyde-3-phosphate with the opposite action of transketolase/transaldolase in non-oxidative PPP (S7 Fig). This suggests that at least some pentose phosphates are generated by non-oxidative PPP. This metabolism is most likely present in hemocytes from uninfected larvae as well as in some types of hemocytes present in the heterogeneous population during infection. *In vivo* experiments begin several hours before full lamellocyte differentiation; therefore, m+1 and m+3 likely combine both pentoses generated by non-oxidative PPP in some hemocytes and pentoses generated by oxidative PPP in other hemocytes.

To summarize this part, hemocytes from both uninfected and infected larvae use non-oxidative PPP to generate pentoses as well as cyclic PPP to generate NADPH. Cyclic PPP increases in hemocytes during infection and particularly in lamellocytes as evidenced by trehalose metabolism. As we present bulk metabolomics data, it is likely that different hemocyte types at different stages of infection use these pathways differently.

The total amount of glyceraldehyde-3-phosphate, which is a product of both glycolysis and PPP, as well as the labeled form, increased during infection, from both labeled glucose and trehalose (Fig 3L). Increased 13C-labeling was much less pronounced for the other glycolytic metabolites, 2/3-phosphoglycerate (Fig 3M) and phosphoenolpyruvate (S8 Fig), but was again strongly detected in lamellocytes from labeled trehalose. Pyruvate was more difficult to detect in our metabolomics analysis (S8 Fig), and in the *ex vivo* experiment, the labeled pyruvate reached detectable levels only after infection (S2 Table). We have previously shown that hemocytes release more lactate during infection [26], which is confirmed here by lower lactate levels in hemocytes and higher levels in hemolymph (S8 Fig). The increased incorporation of 13C into lactate, from both glucose and trehalose (Fig 3N), demonstrates its increased production during infection.

As incorporation of 13C into tricarboxylic acid cycle metabolites is very low (less than 1%) after 40 minutes of *ex vivo* incubation, we compared incorporation into citrate from the experiment with feeding 13C-labeled glucose for 6 hours. Incorporation is the same between uninfected and infected samples (Fig 3O) when values in infected samples are corrected for incorporation into pyruvate, the closest upstream metabolite (S8 Fig); there is a 1.6-fold higher probability that the 13C-labeled pyruvate is used in citrate formation in hemocytes from uninfected larvae than in those from infected larvae. Although the levels of 13C-labeled citrate and malate in the *ex vivo* experiment are very low, and thus difficult to compare, they also do not appear to be dramatically different (S8 Fig). This supports the notion that glucose-derived pyruvate is similarly translocated to mitochondria during infection. Nevertheless, mitochondrial metabolism needs to be further investigated.

Ribose-5-phosphate can be used for *de novo* purine/pyrimidine synthesis. This is indicated by the 13C labeling of AMP/ADP/ATP *in vivo* (S9 Fig), which increases upon infection, even without correction for the lower 13C fraction in ribose-5P. The partial labeling pattern of AMP/ADP/ATP with the m+1 fraction dominating (S9 Fig) indicates that nucleotides are primarily produced from ribose-5P generated by oxidative PPP. Increased labeling of UDP-glucose upon infection suggests that some of the glucose may be used for glucuronate interconversion and glycosylation (S8 Fig), but we did not further analyze this part of metabolism.

In summary, metabolic changes in hemocytes during infection include increased glucose uptake and, in lamellocytes, glucose production from trehalose. The major glucose utilization during infection appears to be in cyclic PPP, where G6P is oxidized in multiple rounds, as evidenced by increased partial G6P labeling. Downstream glycolysis is slightly increased, the increase is more robustly documented by lamellocyte-specific trehalose metabolism and is associated with increased lactate production released from hemocytes, while mitochondrial utilization of glycolysis products appears to be similar.

### Oxidative PPP is required for proper immune response

Our metabolomics analysis shows that carbohydrate uptake and the oxidative PPP increase in hemocytes upon infection. To test the importance of the oxidative PPP, we used the double mutant *Pgd[n39] pn[1] Zw[lo2a]* in glucose 6-phosphate dehydrogenase (*Zwischenferment* or *Zw*), the first enzyme in the oxidative PPP, and in 6-phosphogluconate dehydrogenase (*Pgd*), the third enzyme in the oxidative PPP (Fig 4A); the double mutant is fully viable [27]. We also used a hemocyte-specific RNAi of *Zw* (Fig 4A and 4B) using *Srp-Gal4* and the *Zw* RNAi construct under the UAS promoter (*P{TRiP*.*HMC03068}attP2*). The number of plasmatocytes is not altered in either the double mutant, or hemocyte-specific knockdown of *Zw*, suggesting that regular hematopoiesis is not affected by these manipulations (Fig 4C). However, both in the double mutant and knockdown there is a decrease in lamellocyte number (Fig 4C) and survival (Fig 4D), demonstrating that oxidative PPP is important for effective lamellocyte differentiation and parasitoid killing.

**Fig 4.**
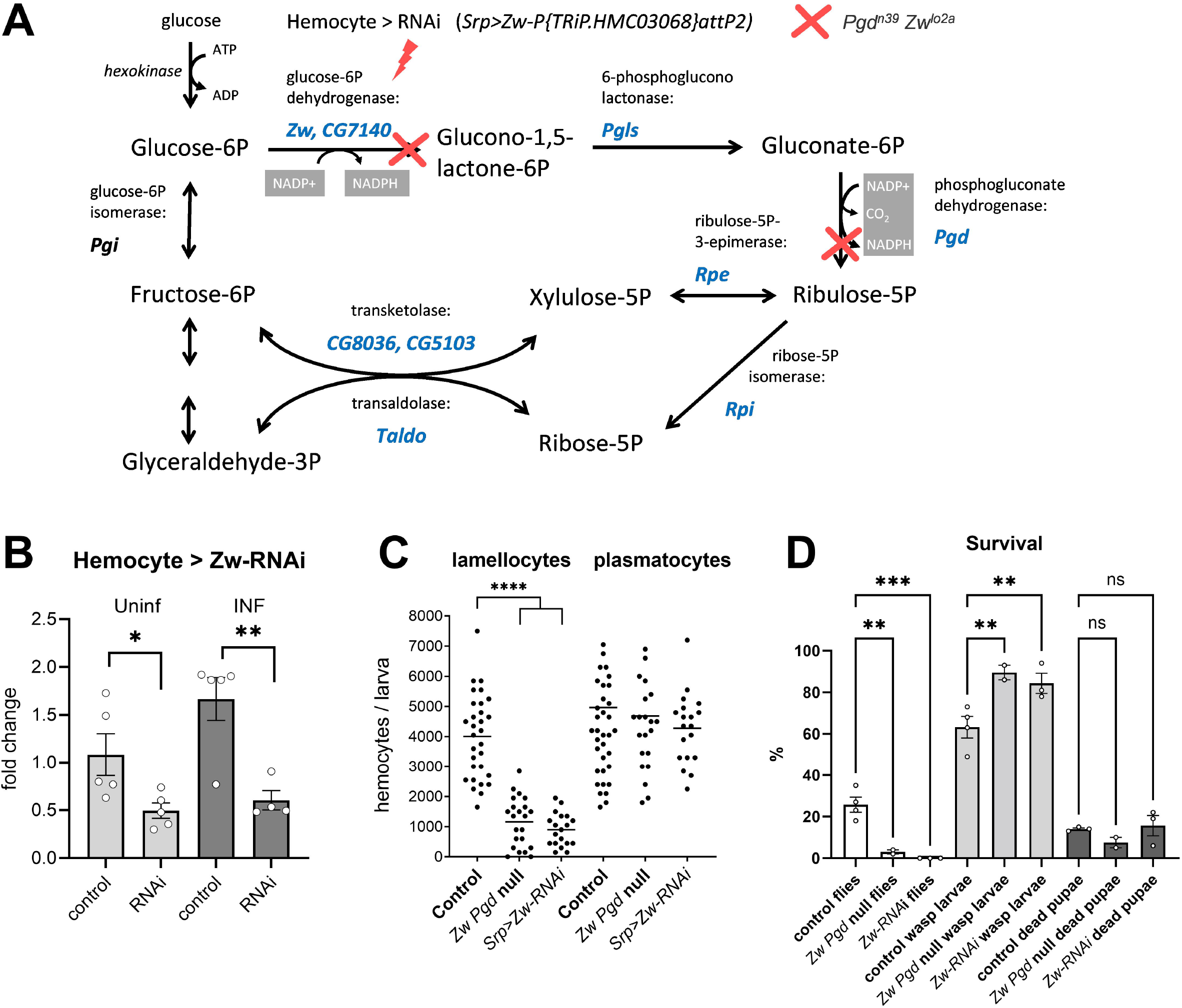
Oxidative pentose phosphate pathway is required for lamellocyte differentiation and resistance. (A) Schematic representation of pentose phosphate pathway with metabolites in black and *Drosophila* genes encoding enzymes in blue. Genetic manipulations include hemocyte-specific RNAi of *Zw* induced by Srp-Gal4 (*Srp>P{TRiP*.*HMC03068}attP2*; red lightning) and double null mutant in *Zw* and *Pgd* (*Pgd[n39] pn[1] Zw[lo2a];* red crosses). (B) Efficiency of hemocyte-specific RNAi of *Zw* tested by RT-qPCR in hemocytes from uninfected (Uninf) and infected (INF) larvae 18 hours after beginning of infection. Bars represent mean value of fold change compared to uninfected control samples (*Zw* expression levels normalized by *RpL32* expression in each sample); dots represent individual samples, error bars represent ± SEM, the control sample (*Srp> P{y[+t7*.*7]=CaryP}attP2*) was compared with RNAi (*Srp>P{TRiP*.*HMC03068}attP2*) using unpaired t test, asterisks indicate p value (* P < 0.05, ** P < 0.01). (C) Number of hemocytes 22 hours after beginning of infection in control (*Srp> P{y[+t7*.*7]=CaryP}attP2*), *Zw Pgd* null mutant (*Pgd[n39] pn[1] Zw[lo2a])* and hemocyte-specific RNAi of *Zw (Srp>P{TRiP*.*HMC03068}attP2)*. Each dot represents number of hemocytes in one larva, line represents the mean, samples were compared by unpaired t test, asterisks indicate p value (* P < 0.05, **** P < 0.0001). (D) Survival of parasitoid wasp infection in control, null *Zw Pgd* mutant and hemocyte-specific RNAi of *Zw*. White bars show the percentage of surviving flies, grey bars the developing parasitoids and dark bars the dead pupae when neither fly nor parasitoid survived. Bars represent mean values; dots represent biological replicates, error bars represent ± SEM; survival rates were compared using ordinary one-way ANOVA with multiple comparisons; asterisks indicate p value (** P < 0.01, *** P < 0.001).

### Systemic trehalose metabolism and carbohydrates supply to hemocytes are required for efficient immune response

Metabolomics with the 13C-labeled trehalose shows that trehalose is metabolized by infection-activated hemocytes. Therefore, we tested the importance of trehalose for the immune response. We first tested the hypomorphic mutation in trehalose-synthesizing gene *Tps1* (*Mi{y[+mDint2]=MIC}Tps1[MI03087] / Tps1[d2]*). *Tps1* hypomorph have normal larval development with normal levels of glucose, glycogen and triglycerides but results in a reduction of trehalose levels to 20% of control [7]. The infected *Tps1* mutant differentiated significantly fewer lamellocytes than control larvae (Fig 5A). Similarly, a null mutant in the *Treh* gene (*Treh[cs1]*) with elevated trehalose, which cannot be converted to glucose either in the circulation or in the cells [21], also differentiated significantly fewer lamellocytes (Fig 5B). We verified that hemocytes from the *Treh[cs1]* mutant were unable to metabolize the 13C-labeled trehalose supplied *ex vivo* (Fig 5C and S2 Table). These results demonstrate that trehalose is important for efficient lamellocyte differentiation.

**Fig 5.**
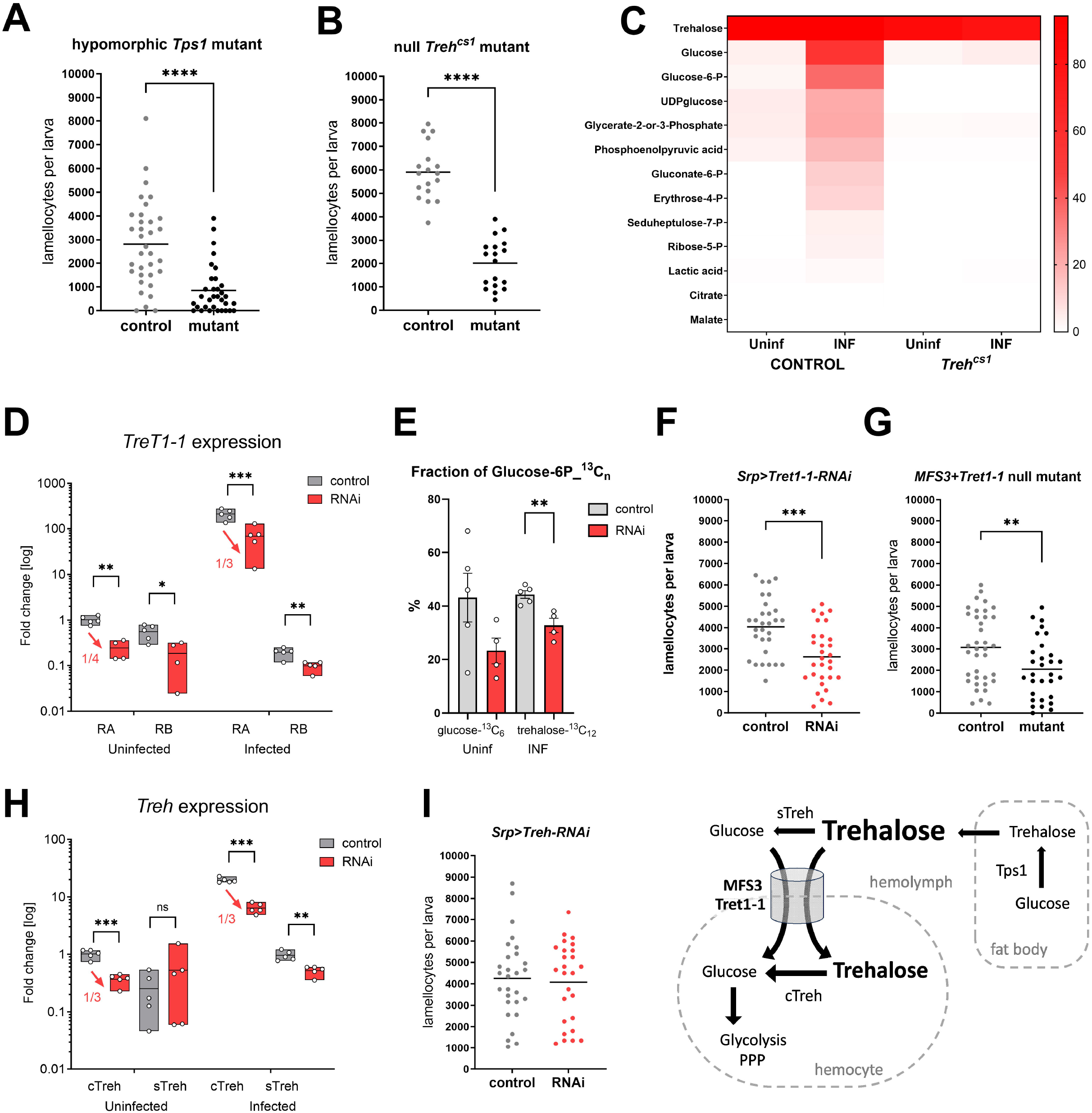
Effects of systemic carbohydrate metabolism and carbohydrate supply to hemocytes on lamellocyte differentiation. (A) Number of lamellocytes 22 hours after beginning of infection in control (heterozygous *Mi{y[+mDint2]=MIC}Tps1[MI03087] / +)* and hypomorphic *Tps1* mutant (*Mi{y[+mDint2]=MIC}Tps1[MI03087] / Tps1[d2]*). Each dot represents number of hemocyte in one larva, line represents mean, samples were compared by unpaired t test, asterisks indicate p value (**** P < 0.0001). (B) Number of lamellocytes 22 hours after beginning of infection in control (heterozygous *Treh[cs1] / CyO Ubi-GFP*) and null *Treh* mutant (homozygous *Treh[cs1] / Treh[cs1]*). (C) Heat map of 13C-labeled fraction of metabolites from control and null *Treh[cs1]* hemocytes in uninfected (Uninf) and infected (INF) conditions incubated *ex vivo* with labeled α,α−trehalose-^13^C_12_. (D) Efficiency of hemocyte-specific RNAi of *Tret1-1* tested by RT-qPCR in hemocytes from uninfected and infected larvae 18 hours after beginning of infection. Box plots (median, 75th and 25th percentiles) show fold change compared to uninfected *Tret1-1-RA* control samples (expression levels were normalized by *RpL32* expression in each sample), each dot represents a biological replicate. Grey boxes represent control (*Srp> P{y[+t7*.*7]=CaryP}attP2*) and red boxes hemocyte-specific *Tret1-1* RNAi (*Srp>P{TRiP*.*HMS02573}attP2*). Unpaired two-tailed t test was used to compare control with RNAi; asterisks indicate p value (* P < 0.05, ** P < 0.01, ***P < 0.001). (E) Fraction (%) of glucose-6-phosphate-^13^C_n_ in hemocytes from uninfected larvae incubated *ex vivo* with labeled glucose (left bars) and from infected larvae incubated with labeled trehalose (right bars). Grey bars represent mean in control (*Srp> P{y[+t7*.*7]=CaryP}attP2*) and red bars mean in hemocyte-specific *Tret1-1* RNAi (*Srp>P{TRiP*.*HMS02573}attP2*). Unpaired two-tailed t test was used to compare control with RNAi; asterisks indicate p value (** P < 0.01). (F) Number of lamellocytes 22 hours after beginning of infection in control (*Srp> P{y[+t7*.*7]=CaryP}attP2)* and in hemocyte-specific *Tret1-1* RNAi (*Srp>P{TRiP*.*HMS02573}attP2*). Each dot represents number of lamellocytes in one larva, line represents mean, samples were compared by unpaired t test, asterisks indicate p value (*** P < 0.001). (G) Number of lamellocytes 22 hours after beginning of infection in control (heterozygous *MFS3[CRISPR] Tret1-1[XCVI]* / Cyo Ubi-GFP) and in *MFS3 Tret1-1* double null mutant (*MFS3[CRISPR] Tret1-1[XCVI]*). Each dot represents number of lamellocytes in one larva, line represents mean, samples were compared by unpaired t test, asterisks indicate p value (** P < 0.01). (H) Efficiency of hemocyte-specific RNAi of *Treh* tested by RT-qPCR in hemocytes from uninfected and infected larvae 18 hours after beginning of infection. Box plots (median, 75th and 25th percentiles) show fold change compared to uninfected *c-Treh* control samples (expression levels were normalized by *RpL32* expression in each sample), each dot represents a biological replicate. Grey boxes represent control (*Srp>P{y[+t7*.*7]=CaryP}attP40*) and red boxes hemocyte-specific *Treh* RNAi (*Srp>P{TRiP*.*HMC03381}attP40*). Unpaired two-tailed t test was used to compare control with RNAi; asterisks indicate p value (** P < 0.01, ***P < 0.001). (I) Number of lamellocytes 22 hours after beginning of infection in control (*Srp>P{y[+t7*.*7]=CaryP}attP40)* and in hemocyte-specific *Treh* RNAi (*Srp>P{TRiP*.*HMC03381}attP40*). Each dot represents number of lamellocytes in one larva, line represents mean, samples were compared by unpaired t test.

Next, we knocked down the *Tret1-1* transporter specifically in hemocytes (*Srp>Tret1-1-RNAi* -*P{TRiP*.*HMS02573}attP2*), which resulted in a reduction of *Tret1-1* expression upon infection to one-third compared to control (Fig 5D). Because knockdown did not prevent the increase in *Tret1-1* expression during infection (it only partially suppressed the increase), *ex vivo* metabolism of 13C-labeled trehalose was only slightly reduced (as indicated by lower incorporation into G6P – Fig 5E and S2 Table), as was the number of lamellocytes (Fig 5F). Since Tret1-1 knockdown did not substantially suppress trehalose uptake, we used a null mutant of *Tret1-1* together with a null mutation of *MFS3*. The double mutant *MFS3[CRISPR] Tret1-1[XCVI]* also significantly reduced the number of lamellocytes, but again only slightly (Fig 5G), suggesting that other highly expressed transporters are providing the carbohydrate supply.

We could not discriminate between glucose and trehalose metabolism in hemocytes by these manipulations. To test the importance of trehalose metabolism in hemocytes, we specifically knocked down *Treh* in hemocytes by Srp-driven RNAi, since the null mutant in *Treh* impairs systemic trehalose metabolism. As with *Tret1-1* knockdown, hemocyte specific knockdown of *Treh* reduced expression to one-third of the control, but did not prevent infection-induced increase in *Treh* expression (Fig 5H). Knockdown of *Treh* also did not affect the number of lamellocytes (Fig 5I); either because reduction of the increase by RNAi was not sufficient, or because trehalase activity is not required for lamellocytes differentiation, as suggested by the c-Treh expression in fully differentiated lamellocytes. Rather than attempt to resolve this issue, we decided to use a mitotic recombination strategy (see the last section of Results).

Since lamellocytes specifically increase the expression of cTreh, we tested the effect of specific mutations in the cytoplasmic version of Treh. The *Treh[c1]* mutation (S10 Fig) should block the utilization of trehalose inside the cells, but not disrupt trehalose-glucose metabolism in the circulation [21]. However, when we incubated *Treh[c1]* mutant hemocytes with 13C-labeled trehalose *ex vivo*, we did not observe any effect on trehalose metabolism in hemocytes (S10 Fig and S2 Table). The failure of the *Treh[c1]* mutation to block intracellular trehalose metabolism could be due to the second start codon in the cytoplasmic *Treh-RA/RD/RG/RE* transcripts (S10 Fig). Therefore, we decided to remove both start codons by replacing 47 bases with a Gal4 coding sequence (Fig 1 and S10). The resulting homozygous *Treh[RAΔGal4]* mutant shows a similar phenotype to the *Treh[c1]* mutant, with 20% lethality during pupal development and two-thirds of emerging adult flies dying within three days of eclosion in our *w*^1118^ genetic background (data not shown). The *Treh[RAΔGal4]* mutant hemocytes still metabolized 13C-labeled trehalose almost normally (S10 Fig and S2 Table), and the mutation did not affect the number of lamellocytes (S10 Fig). Although the overall expression of *Treh* is reduced to one third in the uninfected *Treh[RAΔGal4]* mutant (S11 Fig), it is increased nine fold in the infected mutant (compared to a 25-fold increase in the wild type), indicating compensatory expression (S11 Fig). Using transcript-specific qPCR, we found that this compensatory expression occurs at the transcription start site common to both *Treh-RB* (cTreh) and *Treh-RC* (sTreh) transcripts (see S11 Fig for details). The strong increase in expression of cytoplasmic Treh as well as this compensatory expression in the *Treh[RAΔGal4]* mutant suggests that trehalase activity is very important in lamellocytes.

### Cell-autonomous role of trehalose in hemocytes

Since we could not test the importance of trehalose metabolism in hemocytes with cTreh mutations, we decided to generate mitotic recombination clones [28] in the hematopoietic lineage with the *Treh[cs1]* mutation, which we verified as blocking trehalose metabolism (Fig 5C). We recombined flippase (Flp) target site *FRT42D, Treh[cs1]* mutation and RFP marker on the second chromosome and crossed this line to flies with *FRT42D* and the GFP marker to generate heterozygous *FRT Treh[cs1] RFP / FRT GFP* flies (Fig 6A). The parental flies also carried *Srp-Gal4* and *UAS-Flp* on the third chromosomes, to induce mitotic recombination in the hematopoietic lineage of the progeny. Mitotic recombination resulted in RFP-labeled hemocytes with a *Treh[cs1]* null mutation and their GFP-labeled wild-type siblings (Fig 6A and 6B). When mitotic clones were first induced with only RFP and GFP markers without any mutation, the expected equal number of RFP and GFP sibling hemocytes was detected (Fig 6C), approximately 40% each (the remaining 20% were non-recombined GFP/RFP heterozygous hemocytes), demonstrating the efficiency of the method. A similar result was obtained in uninfected larvae with the *Treh[cs1]* mutation, although there was a minor increase in RFP-labeled hemocytes with the *Treh[cs1]* null mutation compared to GFP-labeled wild-type hemocytes (49% vs. 43%; Fig 6D). The observed difference could be due to the different genetic background of the chromosomal arms that become homozygous after recombination, not necessarily due to the lack of trehalase activity. A comparable result was obtained with lamellocytes from infected animals, where we found similar proportions of red and green cells - the small difference observed is consistent with the difference in uninfected larvae (Fig 6E). Generating clones with the *Treh[cs1]* null mutation did not change the total number of lamellocytes compared with larvae without induced recombination (Fig 6F). These results suggest that trehalase does not play an important, cell-autonomous role, in lamellocyte differentiation, which is consistent with the expression of cytoplasmic trehalase only in fully differentiated lamellocytes (Fig 1H).

**Fig 6.**
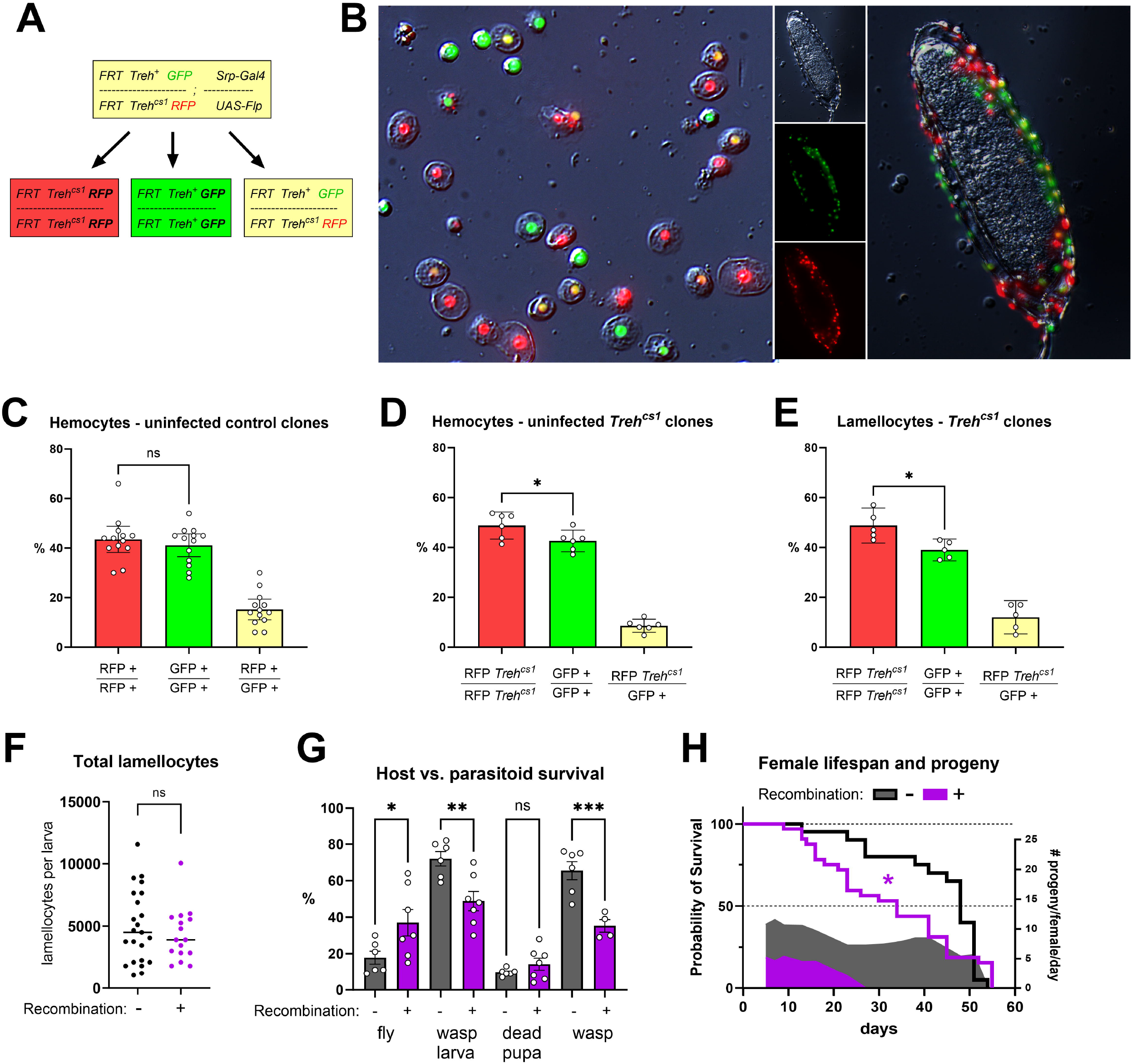
Cell-autonomous role of trehalose metabolism in hemocytes. (A) Generation of hemocyte clones with null *Treh[cs1]* mutation by *Srp-Gal4 UAS-Flp* induced mitotic recombination - genotypes and color of parental and daughter cells after mitotic recombination (red: RFP-marked *Treh[cs1]* mutant clone, green: GFP-marked wild-type sister clone, yellow: RFP/GFP nonrecombinant heterozygous cell). (B) Red, green and yellow-marked hemocytes (both plasmatocytes and lamellocytes) upon mitotic recombination from infected larvae in circulation (left) and attached to parasitoid egg (right). Differential interference contrast (DIC) combined with fluorescence microscopy using 20x objective. (C) Percentage of red- and green-marked wild-type recombinant sister clone hemocytes and yellow-marked non-recombinant heterozygous hemocytes in uninfected control larvae without any mutation. (D) Percentage of red-marked *Treh[cs1]* mutant, green-marked wild-type sister clone hemocytes and yellow-marked non-recombinant heterozygous hemocytes in uninfected larvae. (E) Percentage of red-marked *Treh[cs1]* mutant lamellocytes, green-marked wild-type sister clone lamellocytes and yellow-marked non-recombinant heterozygous lamellocytes in infected larvae. (C-E) Bar represents mean percentage, dot represents counting from one larva, error bars represent ± SEM; red and green samples were compared using unpaired two-tailed t test; asterisks indicate p value (* P < 0.05; ns = not significant). (F-H) Control individuals without recombination due to missing UAS-Flp (*FRT42D GFP / FRT42D Treh[cs1] RFP; Srp-Gal4 / +;* white/grey/black) compared to individuals with *Treh[cs1]* hemocyte mutant clones (*FRT42D GFP / FRT42D Treh[cs1] RFP; Srp-Gal4 / UAS-Flp;* purple). (F) Number of lamellocytes 22 hours after beginning of infection. Each dot represents number of lamellocytes in one larva, line represents mean, samples were compared by unpaired t test, no significant difference. (G) Percentage of surviving adult flies (marked fly), parasitoid wasp larvae, dead pupae (neither fly, nor parasitoid survived) and adult parasitoid wasp (marked wasp). Bars represent mean percentage, dots represent biological replicates, error bars represent ± SEM, samples were compared by ordinary one-way ANOVA with Šídák’s test for multiple comparisons, asterisks indicate p value (* P < 0.05; ** P < 0.01, ***P < 0.001; ns = not significant). (H) Lifespan of females surviving infection (lines) and their daily average production of progeny (shaded area). Lifespan was tested by Gehan-Breslow-Wilcoxon test, median survival 48 days for control and 34 days for females with *Treh[cs1]* mutant clones (P =0.0237). Cumulative average progeny per female was 414 for control and 94 for females with *Treh[cs1]* mutant clones.

To test the effect on resistance, we used these larvae with almost half of the lamellocytes mutant for *Treh*. If trehalose metabolism in the lamellocytes was necessary for parasitoid killing, we would expect lower resistance. However, induction of *Treh[cs1]* mutant clones resulted in the opposite effect, with increased resistance compared to animals with the same genetic background in which clones were not induced due to the absence of Flp (S12 Fig). Although a relatively large variability was observed, the average percentage of surviving adult flies in controls was 18% and never exceeded 30%, whereas in flies with induced clones the average was 37% and reached up to 60% (Fig 6G). Subsequently, the percentage of surviving parasitoid wasps decreased from 65% in controls to 35% when clones were induced (Fig 6G). A possible explanation for these surprising results is that trehalose metabolism in fully differentiated lamellocytes is important for protecting the host from toxic reactions in the encapsulated egg.

Removing this ability in half of the lamellocytes could increase toxicity and thus resistance. To explore further, we looked at the lifespan of adult flies that survived the infection. Male lifespan was comparable (data not shown), however a greater number of females with induced clones died earlier than controls, with the median survival significantly reduced from 48 to 34 days, although the maximum lifespan was comparable (Fig 6H). A more pronounced effect was observed in the production of viable offspring: females with induced clones produced an average of only five viable offspring per female per day and ceased production after 26 days. Whereas, control females produced an average of 8-10 viable offspring per female per day throughout their lifespan (Fig 6H) - cumulatively, control females produced 4.4 times more offspring. It is important to note that verification of the link between the observed reduced fitness and possible increased toxicity during the larval immune response will require further detailed research.

In conclusion, we used mitotic clones to test the cell-autonomous role of trehalose metabolism in hemocytes and found that it is not required for lamellocyte differentiation or resistance. Removing the ability to metabolize trehalose in 40% of lamellocytes improved resistance but reduced the fitness of survivors.

## Discussion

We previously demonstrated a systemic metabolic switch during infection of *Drosophila* larvae by parasitoid wasps, when sugar consumption in non-immune tissues is reduced to provide nutrients for the immune system [4]. We, and others, have also found a strong increase in the expression of the trehalose transporter *Tret1-1* and *Treh* in hemocytes during infection [4,8,9]. This suggests that trehalose is likely an important carbohydrate source for privileged immune cells.

Analysis of transcriptional changes at single cell resolution suggested that larval hemocytes use lipids as the primary source to fuel the TCA cycle in the uninfected state [8]. However, half of the plasmatocytes in all cluster express *MFS3*, a functionally characterized glucose/trehalose transporter [20]; as well as the putative monosaccharide transporter *sut1* [9], suggesting that plasmatocytes also use glucose as a source to be metabolized in the PPP. While plasmatocytes may continue to utilize lipids during infection, as deduced from gene expression, lamellocytes strongly upregulate expression of several carbohydrate transporters and appear to rely much more on saccharides as a source [8]. Our bulk transcriptomic data are consistent with these findings, and 13C-labeled carbohydrate tracing clearly supports these gene expression-based conclusions, showing that plasmatocytes from uninfected larvae do indeed metabolize glucose with a significant fraction metabolized in PPP.

The infection-induced increase in carbohydrate consumption by hemocytes is mediated by a marked increase in the expression of three other carbohydrate transporters besides MFS3 and sut1. Tret1-1, like MFS3, is functionally characterized as a glucose and trehalose transporter [18]. The mild reduction in lamellocyte production even in the *Tret1-1 MFS3* double null mutant indicates that at least one other putative carbohydrate transporters: sut1, CG4607, or CG1208, which have yet to be functionally characterized, contributes to the glucose transport in hemocytes and is sufficient in the double mutant. Notably, increased carbohydrate supply during immune response is so critical that it is ensured by the expression of multiple redundantly functioning transporters (Fig 7). This redundancy in carbohydrate transporters means that knockout of a single, or even two transporters, has no serious impact on immune response. However, the importance of carbohydrate supply to hemocytes is demonstrated as hemocyte-specific silencing of the oxidative PPP by Zw RNAi leads to almost no resistance. We have previously described the significance of a switch in systemic carbohydrate metabolism during the response. Here, we show that hemocytes indeed require carbohydrates for an effective response in order to fuel PPP.

**Fig 7.**
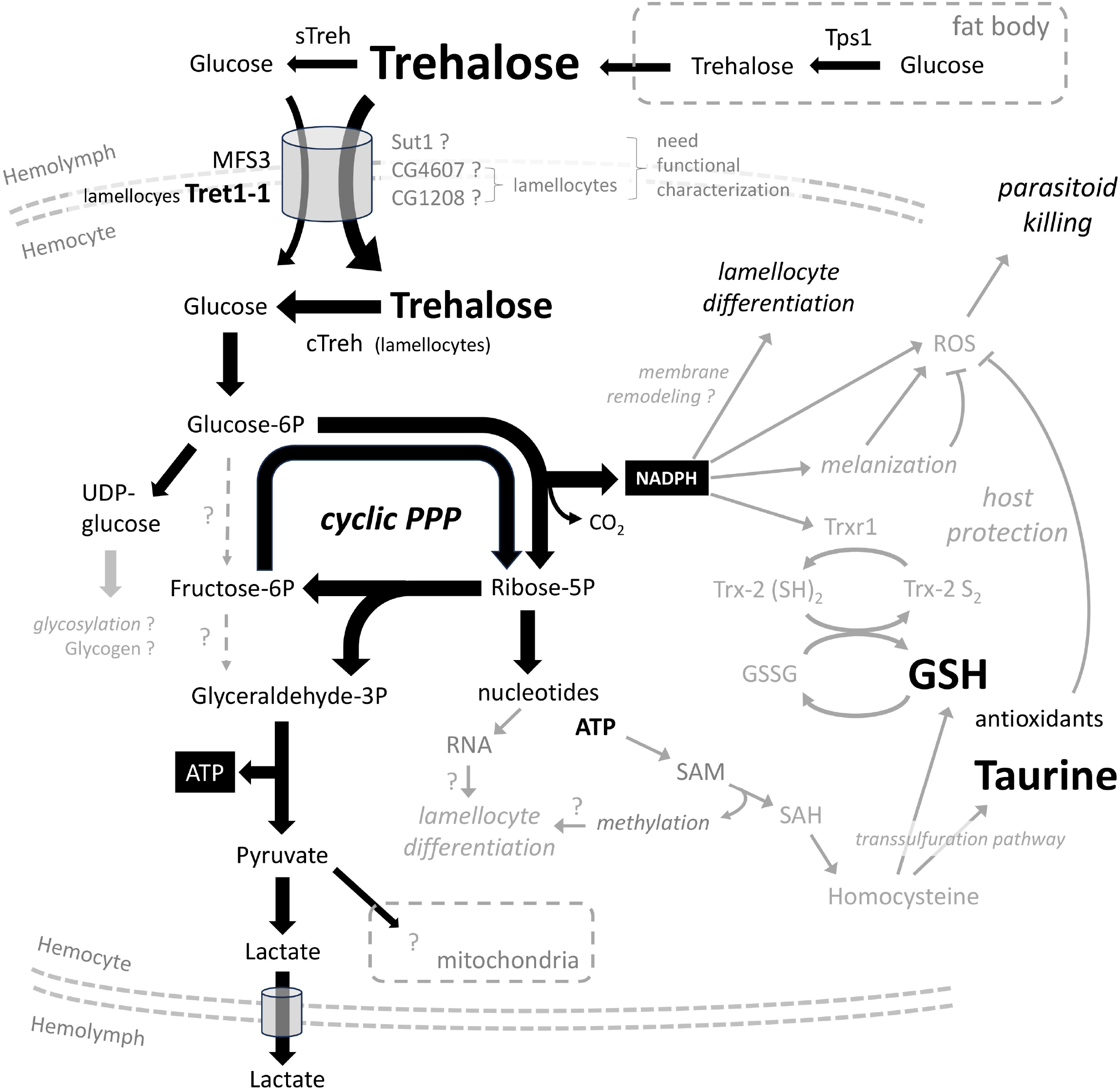
Scheme of hemocyte metabolism during the response to parasitoids and possible links to immune processes. Scheme showing hemocyte metabolism during parasitoid wasp infection. Metabolites, metabolic reactions/pathways and processes marked in black have been studied here, while possible links to other processes and pathways (discussed in the main text) are marked in grey. cTreh, cytoplasmic trehalase; GSH, reduced glutathione; GSSG, oxidized glutathione; PPP, pentose phosphate pathway; SAH, S-Adenosylhomocysteine; SAM, S-Adenosylmethionine; sTreh, secreted trehalase.

The different ways in which glucose is metabolized are determined by the actual needs of the cell (Fig 7 and [13]). If the cell primarily requires ATP, glucose is metabolized by glycolysis producing pyruvate, which can be further metabolized in the mitochondria, and ATP. If the main requirement of the cell is nucleotides (typically a proliferating cell), glucose is metabolized by non-oxidative PPP to form ribose-5-phosphate (bypassing NADPH production by oxidative PPP), which is further metabolized in the de novo synthesis of purines/pyrimidines. If the cell’s primary need is NADPH, glucose is metabolized in cyclic PPP, where the ribose-5-phosphate produced by oxidative PPP is recycled back to G6P, which can enter further rounds of oxidation in PPP. The cell can combine these pathways to obtain the optimal ratio of ATP, NADPH, pyruvate, and pentoses.

Our 13C tracing shows that hemocytes from uninfected larvae metabolize predominantly glucose and very little trehalose. When infected, hemocytes increase glucose metabolism and additionally metabolize trehalose. Expression of cTreh indicates that only differentiated lamellocytes metabolize trehalose; and most likely also metabolize glucose - both MFS3 and Tret1-1 transport trehalose as well as glucose [18,20]. Thus, by tracing 13C-labeled trehalose, we can specifically monitor lamellocyte metabolism. It is important to note that we are observing the metabolism of a heterogeneous pool of hemocyte types, and some hemocytes may use only some of the pathways in which we see 13C labeling.

As summarized in the overall scheme (Fig 7), hemocytes, including lamellocytes, increase their rate of glycolysis and lactate production during infection. Our experiments showed no change in the amount of labeled glucose entering mitochondria, however mitochondrial metabolism can nevertheless be significantly altered during infection.. A significant amount of glucose is shunted into the PPP, both in the absence of infection and during infection. Our 13C tracing experiments show the partial labeling of G6P; this indicates that a significant fraction of hemocytes use cyclic PPP, including lamellocytes, as indicated by trehalose metabolism. Cyclic PPP reoxidizes G6P to generate the maximum amount of NADPH per glucose molecule [3]. A portion is returned to glycolysis at the glyceraldehyde-3P level from PPP, ultimately producing lactate. Some of the ribose-5P formed by oxidative PPP is diverted to de novo nucleotide synthesis. Thus, hemocytes, especially lamellocytes, couple NADPH production via cyclic PPP with *de novo* nucleotide synthesis and ATP production via downstream glycolysis. In addition, the specific pattern of partial labeling of pentoses suggests that some hemocytes generate pentoses via non-oxidative PPP, both in the absence of infection and during infection.

Our 13C tracing experiments and impaired resistance in *Zw*-deficient larvae indicate the importance of cyclic PPP in immune cells. Although the significance of NADPH production in activated immune cells, particularly in conjunction with oxidative burst, is well known [29], we are aware of only one study showing the importance of cyclic PPP in immunity, specifically in mammalian neutrophils [3]. Our findings providing evidence of the importance of cyclic PPP in invertebrate immune cells highlight its evolutionary significance. NADPH generated by cyclic PPP may serve a variety of purposes in immunity [29]. *Zw* null and RNAi animals show that oxidative/cyclic PPP is important for lamellocyte differentiation and for resistance (Fig 7). It is likely that NADPH is used during lamellocytes differentiation for reductive biosynthesis, associated with differentiation processes. For example, lamellocytes undergo significant changes in size, morphology, and adhesion capacity, to distinguish them from their precursors [30]. Therefore, the use of NADPH in reductive biosynthesis of fatty acids and cholesterol can be hypothesized to play a key role in the remodeling of cell membranes during lamellocyte differentiation. In addition, there is a global change in gene expression [8,9] and thus a requirement for RNA synthesis. Consequently, ribose-5P generated by PPP is used for *de novo* nucleotide synthesis, as we observed by 13C labeling in AMP/ADP/ATP. *De novo* synthesized ATP is also required for S-adenosylmethionine production and increased methylation pathway in activated immune cells [31]. Methylation of new molecules (e.g., newly synthesized proteins) may be important during lamellocyte differentiation, but homocysteine, a product of the methylation pathway, is also used in the transsulfuration pathway [32] to produce the antioxidants glutathione and taurine (see below).

Suppression of oxidative/cyclic PPP can also affect resistance. NADPH is needed to produce ROS and these are needed to kill the parasitoid. NADPH oxidase reduces oxygen to superoxide anions, which are subsequently dismutated to hydrogen peroxide [33]. Interestingly, Nappi and Vass detected hydrogen peroxide in plasmatocytes, which eventually attached to the egg, but not in lamellocytes [34]. This suggests that plasmatocytes use cyclic PPP to generate NADPH and hydrogen peroxide; although, our work does not exclude other sources of NADPH, such as isocitrate dehydrogenase and malic enzyme. Hydrogen peroxide can serve as a signal for further immune stimulation [35,36] or react with nitric oxide to form the hydroxyl radical, a very potent ROS [33]. Lamellocytes expressing the prophenoloxidase PPO3 and crystal cells expressing PPO2 [37] produce additional toxic molecules associated with the melanization cascade, which again requires NADPH [12]. Lamellocyte-mediated encapsulation and melanization concentrate the toxic reaction within the capsule, which is crucial for resistance. Thus, NADPH produced by oxidative PPP may be required for resistance due to reductive biosynthesis during lamellocyte differentiation and encapsulation and due to the production of toxic molecules inside the capsule. To kill the parasitoid, the metabolic activities of different hemocytes need to be coordinated. Resolving these roles requires time-controlled genetic manipulation of cyclic PPP, ROS production and melanization cascade specific to the cell type in combination with, for example, genetically encoded metabolic sensors. Such manipulations would make a model of the *Drosophila* response to a parasitoid wasp infection an invaluable tool for investigating the role of cyclic PPP in immunity.

Trehalose is specifically metabolized by lamellocytes because they express cytoplasmic trehalase. The compensatory expression of an alternative transcript in a mutant of cTreh raises the question of why the capacity to metabolize trehalose is so important in lamellocytes. While systemic metabolism of trehalose is necessary for efficient lamellocyte differentiation, metabolism of trehalose within differentiating hemocytes is not. Thus, trehalose in the circulation appears to be important for maintaining adequate glucose levels [7] for differentiating hemocytes. Based on the expression of cTreh, trehalose is only metabolized by fully differentiated lamellocytes in a cyclic PPP, i.e. generating NADPH. Survival rate of individuals with almost half of their hemocytes mutant for *Treh* is rather increased compared to controls. Thus, trehalose metabolism in hemocytes does not appear to be important for resistance mechanisms. Encapsulation and melanization are thought to play a role not only in killing the pathogen, but also in protecting the host from its own toxic immune reaction [12]. Based on our results, we propose that trehalose metabolism in lamellocytes may play a specific role in protecting the host: (1) Trehalose is metabolized by cyclic PPP, which generates NADPH. NADPH is required for the reduction of GSSG to GSH, which we observed to increase in hemocytes after infection. Most hemocytes highly express genes of the thioredoxin system, but according to scRNAseq studies [8,9], lamellocytes show an even stronger expression. The thioredoxin system produces antioxidants, including GSH. However, it is important to add that we do not provide direct evidence that lamellocytes specifically are responsible for the observed increase in antioxidants. (2) Larvae with half of their hemocytes deficient in *Treh* are in fact more resistant than controls, however surviving adults show reduced fitness. This suggests that trehalose metabolism in cyclic PPP is important specifically for antioxidant production and thus host protection from the toxic reaction occurring within the melanizing capsule. Reducing this protection could increase the toxicity of the reaction, leading to the observed increased resistance, but also harming the host. This could manifest itself in a number of ways, such as the observed reduced production of viable offspring by surviving females. Molina-Cruz et al. showed that mosquito strains with higher ROS levels survived bacterial and Plasmodium infections at a higher rate, while dietary antioxidant supplementation reduced resistance[38]. Interestingly, the same antioxidants also significantly improved age-related loss of fecundity in mosquitoes [39]. It is important to emphasize that we do not know whether the observed reduced fitness is directly caused by increased toxicity during the larval immune response. Further studies are needed to verify the connection between trehalose metabolism in cyclic PPP in lamellocytes and the production of antioxidants to protect the host while responding to the parasitoid.

In summary, an effective immune response to parasitoid wasp infection requires rapid and coordinated hemocyte activity, which includes lamellocyte differentiation. Additionally, capsule formation around the parasitoid egg is required, associated with a melanization cascade, and thus production of toxic molecules within the capsule. Lastly, protection of the host cavity from this toxic reaction is required for host survival. All these actions require changes in carbohydrate metabolism in hemocytes (Fig 7). Here we show that systemic trehalose metabolism, including synthesis by Tps1 and conversion to glucose by Treh, is essential for adequate carbohydrate supply to hemocytes during infection, for lamellocyte differentiation and resistance. Hemocyte supply is ensured by the expression of several carbohydrate transporters. While glucose is generally metabolized by hemocytes, trehalose is specifically metabolized only within lamellocytes by cytoplasmic trehalase. Here, we demonstrate that both glucose and trehalose are metabolized by PPP, and in particular by cyclic PPP, which oxidizes G6P in multiple rounds to maximize NADPH production. PPP also connects to downstream glycolysis, which produces ATP and ends with the release of lactate. PPP and its connections to several metabolic pathways support various activities required in the response to parasitoids. (1) We have shown that oxidative PPP is required for lamellocyte differentiation, implicating a role for NADPH in reductive biosynthesis, for example in membrane remodeling. Differentiation could also be promoted by ribose-5P associated with nucleotide synthesis, which is required for broad changes in gene expression (new RNA and methylation). (2) We have shown that oxidative PPP is required for resistance, which likely involves a role of NADPH in the melanization cascade and in the ROS production, both required for pathogen killing. (3) We observed increased production of the antioxidants glutathione and taurine, which requires NADPH/Trxr1-mediated reduction of GSSG to GSH and could also be promoted by coupling PPP-produced ribose-5P to the ATP-SAM-homocysteine-transsulfuration pathway. Antioxidants could explain the observed effects on the reduced fitness of trehalase knockout in hemocyte clones. Nevertheless, the potential link between sugars metabolized in PPP and host protective mechanisms requires further work, such as manipulating the thioredoxin system specifically in fully differentiated lamellocytes and studying the effects on larvae as well as development and physiology of surviving animals. It is difficult to separate resistance mechanisms from host protection when, for example, NADPH produced by cyclic PPP appears to be required for both. At first, we were surprised that although trehalose metabolism seems to be important in hemocytes, we did not observe any effect on resistance. However, from the evolutionary perspective, protecting the host from its own immune response is no less important and probably no less energetically demanding. In the long term, the trade-off between higher survival and lower reproductive fitness may be of great evolutionary importance.

## Materials and methods

### Fly strains and cultivation

*Drosophila melanogaster* strain *w*^*1118*^ (FBal0018186) in Canton S genetic background (FBst0064349) was used as a control line unless otherwise stated. Strains *Pgd*^*n39*^ *pn*^*1*^ *Zw*^*lo2a*^ (FBst0006033), UAS-Zw-RNAi *Zw*^*HMC03068*^ (FBal0292280), UAS-Tret1-1-RNAi *Tret1-1*^*HMS02573*^ (FBal0281575), UAS-Treh-RNAi *Treh*^*HMC03381*^ (FBal0292531) and control lines for RNAi *y*^*1*^ *v*^*1*^; *P{CaryP}attP2* (FBst0036303) and *y*^*1*^ *v*^*1*^; *P{CaryP}Msp300attP40* (FBst0036304) were obtained from the Bloomington Drosophila Stock Center. Strains *Treh*^*c1*^ (FBal0321693), *Treh*^*cs1*^ (FBal0321690), *Tps1*^*MI03087*^ (FBal0260512) and *Tps1*^*d2*^ (FBal0302039) were obtained from T. Nishimura, *MFS3*^*CRISPR*^ (FBal0366542) and *Tret1-1*^*XCVI*^ (FBal0319692) were obtained from S. Schirmeier. The *SrpD-Gal4* strain (FBtp0020112) was obtained from M. Crozatier, backcrossed into the *w*^*1118*^ background, and recombined with P{tubP-GAL80ts}2 (FBti0027797), which was also backcrossed into *w*^*1118*^ background, to generate the *w*^*1118*^; +/+; *SrpD-Gal4* P{tubP-GAL80ts}2 line with Gal4 expression in all hemocytes but very low expression in the fat body at 25°C (expression in the fat body is only present at 29°C in this line). Line *w*^*1118*^; *P{ry[+t7*.*2]=neoFRT}42D, Treh*^*cs1*^, *P{w[+mC]=Ubi-mRFP*.*nls}2R / CyO; SrpD-Gal4 / TM6B* was generated by recombination of *P{ry[+t7*.*2]=neoFRT}42D P{w[+mC]=Ubi-mRFP*.*nls}2R* (FBst0035496) with *Treh*^*cs1*^ and by crossing to *SrpD-Gal4*. Line *w*^*1118*^; *P{ry[+t7*.*2]=neoFRT}42D, P{w[+mC]=Ubi-GFP*.*nls}2R1 P{Ubi-GFP*.*nls}2R2 / CyO; P{y[+t7*.*7] w[+mC]=20XUAS-FLPD5*.*PEST}attP2 / TM6B* was generated by recombination of *P{FRT(whs)}G13 P{Ubi-GFP*.*nls}2R1 P{Ubi-GFP*.*nls}2R2* (FBst0005826) with *P{ry[+t7*.*2]=neoFRT}42D* (FBti0141188) and by crossing to *P{y[+t7*.*7] w[+mC]=20XUAS-FLPD5*.*PEST}attP2* (FBti0161054). All flies were grown on cornmeal medium (8% cornmeal, 5% glucose, 4% yeast, 1% agar, 0.16% methylparaben) at 25°C.

### Generation of Treh[RAΔG4] mutant

CRISPR-mediated mutagenesis was performed by WellGenetics Inc. using modified methods of Kondo and Ueda [40]. In brief, the gRNA sequence TGATTGCTCGATGGATTCGC[TGG] was cloned into U6 promoter plasmid. Cassette *attP-Gal4-3xP3-RFP*, which contains attP, Gal4, RBS and a floxed 3xP3-RFP, and two homology arms were cloned into pUC57-Kan as donor template for repair. *Treh*-targeting gRNAs and hs-Cas9 were supplied in DNA plasmids, together with donor plasmid for microinjection into embryos of control strain *w*^*1118*^. F1 flies carrying selection marker of 3xP3-RFP were further validated by genomic PCR and sequencing. CRISPR generates a 47-bp deletion allele of *Treh* and is replaced by cassette attP-Gal4-3xP3-RFP (Fig 1). The line is depicted here as *Treh*^*RAΔG4*^. *Treh*^*RAΔG4*^ was 10 times backcrossed to our control *w*^*1118*^ genetic background.

### Parasitoid wasp infection

Parasitoid wasps *Leptopilina boulardi* were reared on sugar agar medium (6% sucrose, 1.5% agar, 0.75% methylparaben) and grown by infection of wild-type *Drosophila* larvae. Early third instar larvae (72 hours after egg laying) were infected with parasitoid wasps (= time point 0 hours). Weak infection (1-2 eggs per larva) was used for resistance and survival analysis. Strong infection (4-8 eggs per larva) was used for the rest of the experiments to obtain a strong and more uniform immune response. Infections were performed on 60-mm Petri dishes with standard cornmeal medium for 15 minutes with periodic interruption of infecting wasps for weak infection and 45 minutes for strong infection.

### Hemocyte counting

Hemocytes were obtained from larvae by cuticle tearing of one larva in 15 µl PBS and counted based on morphology in Neubauer hemocytometer (Brand GMBH) using differential interference contrast microscopy.

### Resistance, survival and fitness analysis

To determine survival and parasitoid resistance rates, infected/control larvae were placed in fresh vials (typical 1 experiment = 30 larvae/vial and 3 vials/genotype in 3 independent biological replicates of infection). To determine resistance, pupae were dissected 4 days after infection to count melanized wasp eggs (winning host kills pathogen) or surviving wasp larvae (winning parasitoid). For the survival experiment, emerged adult flies were counted as survivors of infection excluding flies without melanized capsule (if no melanized capsule was visible in the abdomen, the fly was dissected). Adult wasps that emerged from pupae were counted as adult winning parasitoids. The lifespan of surviving infected flies was determined by transferring flies to a fresh vial (20 flies per vial) every 2 to 3 days and counting the days until death. Fitness was determined for flies that survived infection by leaving at least 5 males with at least 5 females (maximum 10 females per vial) in a vial, transferring them to a fresh vial every 2 to 3 days, and counting the number of offspring pupae throughout their lifetime.

### Gene expression analysis

Hemolymph was collected by rupturing 50 larvae in 2 µl of PBS on a microscope slide on ice, the hemolymph was transferred into 200 µl of Trizol reagent (Ambion), homogenized using a plastic motorized pestle, incubated for 15 min at room temperature and either frozen at -80°C or directly followed by RNA isolation using a Direct-zol RNA microprep kit (Zymo Research) according to the manufacturer’s protocol. Reverse transcription was performed using PrimeScript reverse transcriptase (Takara) and gene expression was analyzed using the TP SYBR 2x mastermix (TopBio) on a CFX 1000 Touch Real time cycler (BioRad). Expression of a specific gene in each sample was normalized to expression of *RpL32* (FBgn0002626).

Primers:

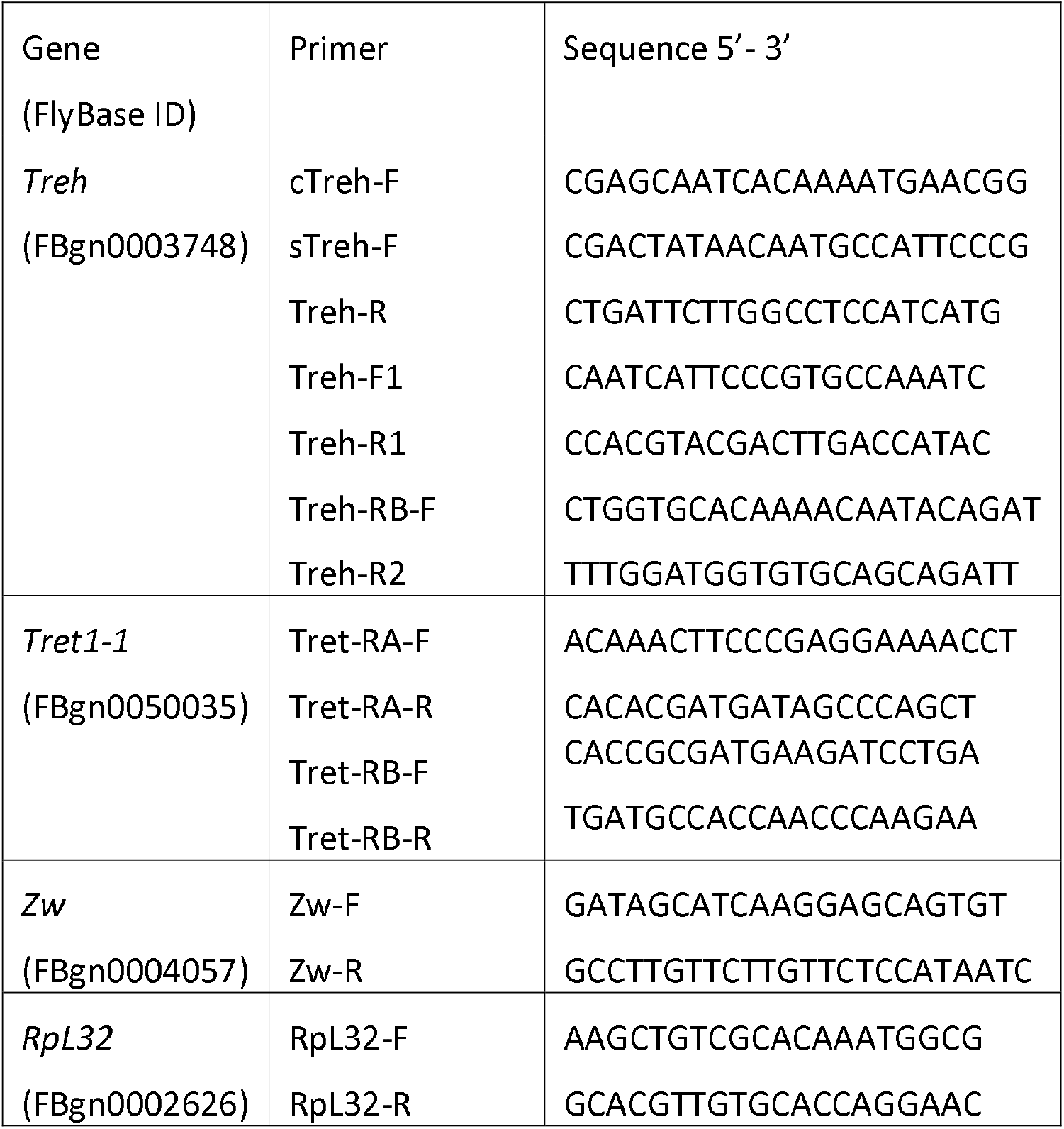

### Bulk RNAseq analysis

RNA was extracted from circulating hemocytes (72 hours after egg laying = time of infection = 0 hours, 81 hours after egg laying = 9 hours post infection/hpi and 90 hours after egg laying = 18 hpi), from lymph glands (9 and 18 hpi) and from wing discs (9 hpi) of uninfected and infected third instar *w*^*1118*^ larvae. Circulating hemocytes were obtained by ripping 100 larvae in ice-cold PBS directly into 1.5 mL centrifuge Eppendorf tubes, centrifuging 5 min at 360xg, removing the supernatant, and isolating RNA using Trizol reagent (200 µL) (Ambion) according to the manufacturer’s protocol. Lymph glands and wing discs were dissected from larvae in ice-cold PBS, transferred to 1.5 mL centrifuge Eppendorf tubes with Trizol (200 µL) reagent (Ambion), homogenized using a plastic motorized pestle, followed by Direct-zol RNA microprep kit (Zymo Research) according to protocol. Frozen total RNA samples were sent to the Genomics Core Facility (EMBL Heidelberg) for preparation of barcoded 3’-end seq forward libraries, followed by deep uni-directional sequencing of 75-base long reads using Illumina NextSeq. Trimmed reads in Fastq files were mapped to the BDGP *Drosophila melanogaster* Release 6.29 genomic sequence using the Mapper for RNA Seq in Geneious prime software (Biomatters). Normalized counts of reads mapped to each gene annotation were calculated as transcripts per million (TPM), expression levels were compared using the DESeq2 method in Geneious prime software, and data were exported to an Excel file (S1 Table).

### Metabolomics and stable 13C isotope tracing

13C-labeled glucose feeding. 13C-labeled glucose (D-Glucose-^13^C_6_ isotope purity ≥99 atom % 13C - Sigma-Aldrich) was added to the semi-defined diet (per 100 ml: 0.62 g of agar, 8 g of brewer’s yeast, 2 g of yeast extract, 2 g of peptone, 3 g of sucrose, 3 g of unlabeled glucose, 0.05 g of MgSO_4_ x 6H_2_O, 0.05 g of CaCl_2_ x 2H_2_O, 600 µl of propionic acid, 1 ml of 10% p-hydroxy-benzoic acid methyl ester in 95% ethanol). The diet mixture was brought to a boil, then cooled to 50-60°C with stirring and p-hydroxy-benzoic acid and propionic acid were added. 2 ml of medium was taken into a Falcon tube and 100 µl of 13C-labeled glucose (600 mg/ml) was added (50% of the glucose in the diet was labeled). 1 ml of diet was poured into a glass vial. 100 uninfected or infected larvae (16 hours after the start of infection) were placed in the vial for 6 hours. The larvae were then removed from the vial, washed twice in water and once in PBS and placed on a microscope slide covered with parafilm, and the PBS residue was removed with filter paper. 4 µl of PBS was added to the larvae and each larva was ruptured, 20 µl of hemolymph was collected and transferred to sterile 1.5 ml Eppendorf polypropylene centrifuge tubes with 60 µl of PBS. Larvae were washed with an additional 25 µl of fresh PBS and 20 µl was recovered in the same tubes. Samples were centrifuged for 5 minutes at 360xg. 95 µl of supernatant (with extracellular metabolites) was removed, 380 µl of cold acetonitrile-methanol (1:1) extraction buffer was added and placed in liquid nitrogen. To collect metabolites from pelleted hemocytes, 50 µl of water was added, frozen in liquid nitrogen/thawed at 37°C three times (to disrupt the rigid cells of hemocytes), then 200 µl of cold acetonitrile-methanol (1:1) was added and stored at -80°C until LC-HRMS analysis.

Ex vivo hemocyte incubation with 13C-labeled glucose and trehalose. Larvae were washed first with distilled water and then with PBS to reduce contamination. Larval hemolymph was collected by carefully tearing the larvae on a glass microscope slide covered with parafilm. Hemolymph from 50 larvae was immediately collected into sterile 1.5 ml Eppendorf polypropylene centrifuge tubes prefilled with 100 µl PBS and centrifuged for 5 min at 25°C, 360xg. The supernatant was then removed and the cells were mixed with labeled medium (5mM ^13^C_12_ labeled or unlabeled trehalose, 0.5mM ^13^C_6_ labeled or unlabeled glucose, 5mM proline, 0.3mM methionine and 5mM glutamine, all reagents from Sigma/Merck,) supplemented with gentamicin (10 mg/ml; Gibco), amphotericin B (250 µg/ml; Gibco) and 0.1 mM phenylthiourea (PTU; Sigma/Merck) to prevent melanization. The hemocytes were then incubated for 40 min at 25°C and 80-90% humidity. The cells were then centrifuged for 5 min at 25°C, 360xg, the supernatant was removed, the cells were mixed with 50 µl of cold PBS and frozen in liquid nitrogen/thawed at 37°C three times (to disrupt the rigid hemocyte cells). Finally, 200 µl of cold acetonitrile-methanol (1:1) was added and samples were stored at -80°C until LC-HRMS analysis.

Frozen samples were melted on ice, then internal standards, p-fluoro-DL-phenylalanine (Sigma-Aldrich, Saint Luis, MI, USA) was added to the extraction buffer, both at a final concentration of 200 nmol/mL. Samples were homogenized using a TissueLyser LT (Qiagen, Hilden, Germany) set to 50 Hz for 5 min (with a rotor pre-chilled to −20°C). Homogenization and centrifugation (at 20 000×g for 5 min at 4°C) was repeated twice and the two supernatants were combined. Samples were analyzed by a high-resolution mass spectrometer (Orbitrap-Q Exactive Plus) coupled to a Dionex Ultimate 3000 liquid chromatograph and a Dionex open autosampler (all from ThermoFisher Scientific, Waltham, MA, USA) as previously described [41]. Data were acquired and metabolites identified using an in-house Metabolite Mapper platform equipped with an internal metabolite database in conjunction with Xcalibur™ software (v4.0, ThermoFisher Scientific, Waltham, MA, USA). All metabolites were quantified relatively using the areas under respective chromatographic peaks. The data were normalized to the total content of all screened unlabeled metabolites - the peak area of the metabolite in a particular sample was divided by the peak area of the same metabolite of the selected reference sample and this procedure was repeated for each individual unlabeled metabolite. These ratios of all metabolites in one particular sample were averaged to determine a normalization factor. We then divided the measured peak area by the normalization factor for that sample to obtain the normalized peak area values (S2 Table).

### Generation of hemocyte mutant clones by mitotic recombination

The *FRT RFP; SrpD-Gal4* control line (*w*^*1118*^; *P{ry[+t7*.*2]=neoFRT}42D, P{w[+mC]=Ubi-mRFP*.*nls}2R / CyO; SrpD-Gal4 / TM6B*) or the *FRT Treh*^cs1^ *RFP; SrpD-Gal4* mutant line (*w*^*1118*^; *P{ry[+t7*.*2]=neoFRT}42D, Treh*^cs1^, *P{w[+mC]=Ubi-mRFP*.*nls}2R / CyO; SrpD-Gal4 / TM6B*) were crossed with either flippase-free *FRT GFP* line (*w*^*1118*^; *P{ry[+t7*.*2]=neoFRT}42D, P{w[+mC]=Ubi-GFP*.*nls}2R1 P{Ubi-GFP*.*nls}2R2 / CyO)* as control without clone induction or with *FRT GFP; UAS-Flp* line (*w*^*1118*^; *P{ry[+t7*.*2]=neoFRT}42D, P{w[+mC]=Ubi-GFP*.*nls}2R1 P{Ubi-GFP*.*nls}2R2 / CyO; P{y[+t7*.*7] w[+mC]=20XUAS-FLPD5*.*PEST}attP2 / TM6B*) to induce mitotic clonal recombination in hemocytes by expressing flippase using *SrpD-Gal4* driver. Larvae with ubiquitous red and green fluorescence, i.e. without balancers, were selected and dissected in PBS on a microscope slide to obtain hemocytes. Images of hemocytes were taken using red and green fluorescence and differential interference contrast microscopy. Merged images were used to count green, red and heterozygous yellow hemocytes.

### Immunohistochemistry

The central nervous system of infected and non-infected third instar larvae were dissected and stained according to standard protocols. The following primary antibodies were used: GFP anti-chicken (Abcam, 1:1000), Rabbit anti-Rumpel (1:500; [42]), Elav anti-rat and Repo anti-mouse (Developmental Studies Hybridoma Bank, 1:5), Tret1-1 anti-guinea pig, 1:50; [43]). All secondary antibodies conjugated to Alexa Fluor 488, Alexa Fluor 568 or Alexa Fluor 647 were used at a ratio of 1:1000 (Thermo Fisher Scientific). Confocal images were obtained using a Zeiss LSM 880 (Zeiss, Oberkochen, Germany) and analyzed using Fiji [44].

## Data analysis

Data were analyzed and graphed using GraphPad Prism (GraphPad Software), with specific statistical tests shown in the legend of each figure.

## Acknowledgments

The authors acknowledge funding from the Grant Agency of the Czech Republic to TD (Project 20-09103S; www.gacr.cz) and from the European Union’s Horizon 2020 research and innovation programme under the Marie Skłodowska-Curie grant agreement No 867430 to MK (IMMUNETREH). We thank Dr. Takashi Nishimura, Dr. Stefanie Schirmeier, Michele Crozatier and Bloomington Drosophila Stock Center for fly and wasp stocks. We thank to Lucie Hrádková for laboratory management and Marcela Jungwirthová for project managemenet, and all members of Doležal and Šimek laboratories for their help with work. We thank Dr. Vladimír Beneš and Genomics Core Facility (EMBL Heidelberg, Germany) for RNAseq services and WellGenetics Inc. (New Taipei City, Taiwan) for CRISPR-mediated mutagenesis. We thank Dr. Jason Tennessen for advice on metabolomics.

## Supporting information

**S1 Fig. Cytoplasmic trehalase expression in hemocytes and wing imaginal disc**.

*Treh[RAΔG4]* with a knocked-in Gal4 in the *Treh-RA* transcriptional variant drives UAS-GFP expression in the cytoplasmic trehalase expression pattern (cTreh>GFP). (A,B) Differential interference contrast (DIC) combined with fluorescence microscopy using 20x objective. (A) Hemocytes from uninfected 3^rd^ instar larvae with no expression of cTreh>GFP. (B) Hemocytes from larvae 22 hours after wasp infection - while large flat lamellocytes express cTreh>GFP, no expression was detected in both round and spread plasmatocytes. (C) Parasitoid egg encapsulated by lamellocytes expressing cTreh>GFP 24 hours after infection - DIC (left), green fluorescence (middle) and merged (right) image from a Leica Thunder Imaging Systems microscope using 20x objective. (D,E) Wing imaginal disc expressing cTreh>GFP similarly in uninfected (D) and infected (E) larvae. (D) DIC (left), green fluorescence (middle) and merged (right) image using 20x objective. (E) Merged image only.

**S2 Fig. Cytoplasmic trehalase is expressed in glial cells of the central nervous system**.

*Treh[RAΔG4]* with a knocked-in Gal4 in the *Treh-RA* transcriptional variant drives UAS-GFP expression in the cytoplasmic trehalase expression pattern (cTreh>GFP). (A-C, E-G) are single focal plane images that show cTreh>GFP (green) expression, Rumpel (magenta) predominantly expressed in ensheathing glia cells and Elav (blue) a neuronal specific marker. (D and H) are maximum projections highlighting the expression of cTreh>GFP (green). cTreh>GFP (green) shows overlap with Rumpel (magenta) but not Elav (blue), suggesting that the cytoplasmic trehalase is expressed in ensheathing glia, but not neurons. (I-K, M-O) are single confocal sections, cTreh>GFP (green), Repo (magenta) expressed in all glial nuclei and Tret1-1 (blue) expressed in perineurial glia, the outermost glial cell layer of the blood-brain barrier. (L and P) show a Z projection of larval brains with cTreh>GFP (green) and Repo (magenta) staining. There is overlap in expression of cTreh>GFP (green) and Repo (magenta). There is no evidence of cTreh>GFP (green) expression in perineurial glia (blue). (A-D and I-L) are brains of uninfected 3^rd^ instar larvae. (E-H and M-P) are 3^rd^ instar larval brains of infected animals. (A, C-D, E, G-H, I, K-L, M, O-P) show an overview of the central nervous system using 20x objective. (B, F, J, N) show a close up of the ventral nerve cord using 63x objective.

**S3 Fig. Scheme of 13C isotope labeling of metabolites in the cyclic pentose phosphate pathway**. The cyclic pentose phosphate pathway (PPP) can recycle six pentoses – 5C in black boxes (ribose-5-phosphate or xylulose-5-phosphate), which are formed from glucose-6-phosphate by oxidative PPP (OxPPP), into four hexoses - 6C in blue boxes (glucose-6-phosphates) and two trioses - 3C in blue boxes (glyceraldehyde-3-phosphates) by using transketolase and transaldolase. The recycled glucose-6-phosphates can enter further rounds of cyclic PPP to maximize NADPH production. Glyceraldehyde-3-phosphates can re-enter glycolysis. Metabolizing fully labeled glucose-^13^C_6_ in cyclic PPP produces partially labeled glucose-6-phosphate. Initially, when labeled metabolites begin to enter cellular metabolism and represent a minority fraction, the most common intermediate in cyclic PPP (and also in glycolysis) is fully labeled glyceraldehyde-3-phosphate-^13^C_3_, which combines with unlabeled metabolites to form glucose-6-phosphate-^13^C_3_ partially labeled at 3 carbons (pink highlight). Later, when more labeled metabolites enter cellular metabolism, the most common product of cyclic PPP is glucose-6-phosphate-^13^C_2_ (three of seven possible combinations after the first round, as highlighted in yellow). Red circles represent ^13^C carbons, gray circles ^12^C carbons.

**S4 Fig. Thioredoxin system in hemocytes**. (A) Scheme of the thioredoxin system in *Drosophila*.

The reduction of the disulfide thioredoxin Trx S_2_ to the reduced dithiol form Trx (SH)_2_ is catalyzed by NADPH-dependent thioredoxin reductase (Trxr). Thioredoxin reduces glutathione disulfide (GSSG) to glutathione (GSH), an antioxidant that scavenges radicals via glutathione peroxidase (Gpx). *Drosophila* thioredoxin Trx-2 may also be a substrate for thioredoxin peroxidases (peroxiredoxins, Prx) that detoxify peroxides. (B) Bulk RNAseq of genes of the thioredoxin system expressed in circulating hemocytes from uninfected (Uninf, light gray bars) and infected (INF, dark gray bars) larvae 18 hours after the start of infection. Hemocytes express thioredoxin reductase *Trxr1*, thioredoxin *Trx-2*, and various putative peroxiredoxins and glutathione peroxidases. Expressions are shown in transcripts per million (TPM), bars represent mean values; dots represent biological replicates, error bars represent ± SEM. (C) Single-cell RNAseq plot of *Trxr1* expression in hemocytes from wasp-infected larvae for 48 hours, obtained from the single-cell RNA-seq data portal of DRSC/Perrimon lab (https://www.flyrnai.org/scRNA/), showing stronger expression in lamellocytes. *Atilla* (lamellocyte marker) and *Hml* (plasmatocyte marker) expression is shown for comparison. (D) Graph of *Trx-2* expression in hemocytes based on single-cell RNA-seq data portal (https://www.flyrnai.org/tools/single_cell/web/) showing that a higher percentage of lamellocytes express *Trx-2* more strongly than other hemocytes.

**S5 Fig. Partial labelling of 13C glucose-6-phosphate and ribulose-5-phosphate ex vivo**.

Bars show the mean normalized peak area of glucose-6-phosphate and ribulose-5-phosphate with different numbers of ^13^C in the molecules (m+1 with one ^13^C … m+6 with six ^13^C) measured in hemocytes incubated *ex vivo* with either D-glucose-^13^C_6_ (gray) or α,α-trehalose-^13^C_12_ (blue). Samples were obtained from hemocytes of uninfected (depicted by Uninf) or infected (depicted by INF) larvae. Each dot represents a biological replicate.

**S6 Fig. Partial labelling of 13C glucose-6-phosphate, ribulose-5-phosphate and ribose-5-phosphate in vivo**.

Bars show the mean normalized peak area of glucose-6-phosphate, ribulose-5-phosphate and ribose-5-phosphate with different numbers of ^13^C in the molecules (m+1 with one ^13^C… m+6 with six ^13^C) measured in hemocytes from larvae fed D-glucose-^13^C_6_ for six hours. Samples were obtained from hemocytes of uninfected (depicted by Uninf, light grey) or infected (depicted by INF, dark grey) larvae. Each dot represents a biological replicate.

**S7 Fig. Scheme of 13C isotope labeling of metabolites in the non-oxidative pentose phosphate pathway**.

The nonoxidative pentose phosphate pathway (PPP) produces ribose-5-phosphate from the glycolytic products fructose-6-phosphate and glyceraldehyde-3-phosphate by using transketolase and transaldolase. Metabolism of fully labeled glucose-^13^C_6_ in non-oxidative PPP produces mostly partially labeled ribose-5-phosphates (with ^13^C_4_ /m+4 being least likely) and less fully labeled ribose-5-phosphate-^13^C_5_. Red circles represent ^13^C carbons, gray circles ^12^C carbons, orange rectangles represent labeling in ribulose-5-phosphate, grey rectangles in ribose-5-phosphate.

**S8 Fig. Analysis of hemocyte metabolism by 13C stable isotope tracing**.

Bars show the mean metabolite amounts - unlabeled form or labeled with stable 13C isotope, or both stacked, in one bar - expressed by the normalized peak area. Graphs labeled “*in vivo*” in black box - larvae were fed labeled D-glucose-^13^C . Graphs labeled “*ex vivo*” in gray box - hemocytes were incubated *ex vivo* with either labeled D-glucose-^13^C_6_ or α,α−trehalose-^13^C_6_ . Samples were obtained from hemocytes or hemolymph from uninfected (Uninf) or infected (INF) larvae. Phosphoenolpyruvate, pyruvate and UDP-Glucose graphs combine unlabeled (gray) and fully (red) or partially (pink) labeled forms of metabolites; the percentages above the columns express the fraction of the labelled from the total amount. Total Lactate shows combined lactate-12C and lactate-13C from pelleted hemocytes and supernatant representing the hemolymph. Citrate and malate show fully labeled ^13^C_3_ molecules. The sample from infected larvae was compared with that from uninfected larvae using unpaired t test or ordinary two-way ANOVA with multiple comparisons. Asterisks indicate p value (* P < 0.05, ** P < 0.01, *** P < 0.001, **** P < 0.0001) and are either above the bar in the corresponding color of the bar they compare within the stacked bars, or in black for a simple comparison. A bar without asterisks indicates a non-significant difference. Error bars represent ± SEM.

**S9 Fig. Amount of AMP, ADP and ATP and their partial labelling in vivo**.

Bars show the mean normalized peak area of AMP, ADP and ATP with different numbers of ^13^C in the molecules (m+1 with one ^13^C …m+5 with five ^13^C) and of unlabeled molecules (^12^C_5_) measured in hemocytes from larvae fed D-glucose-^13^C_6_ for six hours. Samples were obtained from hemocytes of uninfected (Uninf, light gray) or infected (INF, dark gray) larvae. Each dot represents a biological replicate. The sample from infected larvae was compared with that from uninfected larvae using ordinary two-way ANOVA with Šídák’s test for multiple comparisons and by unpaired t test; asterisks indicate p value ** P < 0.01, *** P < 0.001, **** P < 0.0001); a bar without asterisks indicates a non-significant difference. Error bars represent ± SEM.

**S10 Fig. Effects of cytoplasmic trehalase-specific mutations**.

Map of the *trehalase* gene with individual transcripts (RA-RG) and sequence from RA first exons depicting wild-type, *Treh*^*c1*^ and *Treh*^*RAΔG4*^ mutations. *Treh*^*c1*^ deletes 20 bp including the first start codon. *Treh*^*RAΔG4*^ deletes 47 bp removing both start codons, which is replaced by a cassette containing the Gal4 coding sequence. Lines show introns, boxes show exons with coding sequence in orange. (B,C) Heat map of 13C-labeled fraction of metabolites from control and *Treh* (B) and *Treh* (C) hemocytes in uninfected (Uninf) and infected (INF) conditions incubated *ex vivo* with labeled α,α−trehalose-^13^C_12_ . (D) Number of lamellocytes 22 hours after beginning of infection in control (*w*^*1118*^) and in the *Treh*^*RAΔG4*^ mutant. Each dot represents number of lamellocytes in one larva, line represents mean, samples were compared by unpaired t test.

**S11 Fig. Compensatory expression of trehalase in the Treh[RA*Δ*G4] mutant**.

Transcript-specific analysis of *trehalase* expression by RT-qPCR 18 hours after the start of infection. (A) The 25-fold increase in *Treh* expression in hemocytes (as measured by expression of the common region for all *Treh* transcripts using Treh-F1/Treh-R1 primers) in control larvae during infection is due to an increase in expression of *Treh-RA(DGE)* transcripts (Fig 1). (B) *Treh-RA(DGE)* expression is disrupted by a knocked-in Gal4 driver in the *Treh* mutant, leading to a one-third reduction in total *Treh* expression in hemocytes of uninfected larvae. (C) The lack of *Treh-RA(DGE)* expression in the *Treh*^*RAΔG4*^mutant is compensated by increased *Treh-RB/RC* expression, 7.5-fold in uninfected and 18-fold in infected larvae, whereas in wild-type larvae, *Treh-RB/RC* expression does not change after infection; analyzed using Treh-RB-F/Treh-R2 primers. (A) This compensatory infection-induced expression leads to an overall 9-fold increase in *Treh* expression in the *Treh*^*RAΔG4*^ mutant compared to a 25-fold increase in infected wild-type larvae. (A-C) Bars show mean fold change compared to control uninfected samples (expression levels were normalized by *RpL32* expression in each sample), each dot represents a biological replicate. Unpaired two-tailed t test (B) and ordinary one-way ANOVA with Tukey’s multiple comparisons test (C) were used to compare samples; asterisks indicate p value ** P < 0.01, **** P < 0.0001); ns indicate non-significant difference. Error bars represent ± SEM.

**S12 Fig. Hemocytes from control larvae unable to induce mitotic recombination in hemocytes**. emocytes from uninfected larvae that were unable to generate clones by mitotic recombination in hemocytes due to the lack of flippase (*FRT42D GFP / FRT42D Treh[cs1] RFP; Srp-Gal4 / +*) - all hemocytes express both GFP and RFP markers (yellow when merged). These larvae served as controls for larvae with mitotic recombination clones in hemocytes (Fig 6). Differential interference contrast (DIC) and fluorescence microscopy using a 40x objective.

**S1 Table. Gene expression analysis by bulk RNAseq of circulating hemocytes, lymph gland and wing disc during parasitoid wasp infection**.

MS Excel sheets with gene expression in circulating hemocytes (first sheet), lymph gland and wing disc (second sheet). RNA was extracted 72 hours after egg laying = time of infection = 0 hours, 81 hours after egg laying = 9 hours post infection/hpi and 90 hours after egg laying = 18 hpi), from hemocytes, lymph glands and wing discs of the third instar *w*^*1118*^ larvae. Barcoded 3’- end seq forward libraries were subjected to deep uni-directional sequencing of 75-base long reads using Illumina NextSeq. Trimmed reads in Fastq files were mapped to the BDGP *Drosophila melanogaster* release 6.29 genomic sequence (gene names correspond to this release) using the Mapper for RNA Seq in Geneious prime software (Biomatters). Normalized counts of reads mapped to each gene annotation were calculated as transcripts per million (TPM), expression levels were compared using the DESeq2 method in Geneious prime software.

**S2 Table. Metabolomics and stable 13C isotope tracing in circulating hemocytes during parasitoid wasp infection**.

MS Excel sheets with stable 13C isotope tracing experiments. Values are raw or normalized areas under respective chromatographic peaks. Data from the following experiments are in individual sheets: [13C in vivo] – raw and normalized data from hemocytes obtained 22 hours after start of infection from uninfected and infected *w*^*1118*^ larvae fed a diet with 50% D-Glucose-^13^C_6_ for the last 6 hours (normalization factor determination in [13C in vivo normalization] sheet). [13C ex vivo] raw and normalized data from hemocytes obtained 22 hours after start of infection from uninfected and infected *Srp>P{y[+t7*.*7]=CaryP}attP2* control larvae and incubated for 40 minutes in medium containing 5mM unlabeled trehalose and 0.5mM ^13^C_6_ labeled glucose or 5mM ^13^C_12_ labeled trehalose and 0.5mM unlabeled glucose (normalization factor determination in [13C ex vivo normalization] sheet). [13C ex vivo Trehcs1], [13C ex vivo Trehc1] and [13C ex vivo TrehRAdG4] sheets - raw data from hemocytes obtained 22 hours after start of infection from uninfected and infected *w*^*1118*^ control and *Treh*^cs1^, *Treh*^*c1*^ *or Treh*^*RAΔG4*^ mutant larvae and incubated for 40 minutes in medium containing 5mM ^13^C_12_ labeled trehalose and 0.5mM unlabeled glucose. [13C ex vivo Srp-Tret1-1-RNAi] raw and normalized data from hemocytes obtained 22 hours after start of infection from uninfected and infected *Srp>P{y[+t7*.*7]=CaryP}attP2* control larvae and larvae with hemocyte-specific Tret1-1 RNAi *Srp>P{TRiP*.*HMS02573}attP2* and incubated for 40 minutes in medium containing 5mM ^13^C_12_ labeled trehalose and 0.5mM unlabeled glucose.

**S1 File. Carbohydrate transport and metabolism gene expression analysis by bulk and single-cell transcriptomics**.

Table with bulk RNAseq gene expressions (transcripts per million - TPM, average values) of genes from SLC2 and SLC17 family of sugar transporters in *Drosophila* – the intensity of the red color corresponds to the TPM value. Bulk RNAseq expressions of selected genes in bar graphs - each dot represents a biological replicate in TPM, bars represent mean ± SEM. Single cell-RNAseq of circulating hemocytes – dot plots with average gene expressions in hemocyte clusters; color gradient of the dot represents the expression level, the size represents percentage of cells expressing the gene per cluster; downloaded from www.flyrnai.org/tools/single_cell/web/. Single cell-RNAseq of circulating hemocytes - t- Distributed Stochastic Neighbor Embedding (t-SNE) plots of Harmony-based batch correction of wasp infected 48 hours data sets downloaded from www.flyrnai.org/scRNA/blood/ (for comparison, plots with plasmatocytes marker *Hml* and lamellocyte marker *Atilla* are shown).

**S2 File. Glycolytic and pentose phosphate pathway gene expression analysis by bulk and single-cell RNAseq**.

Diagram showing metabolic pathways and tables with gene expression corresponding to Fig 2. Table with bulk RNAseq gene expressions (transcripts per million - TPM, average values) of glycolytic and PPP genes in *Drosophila* – the intensity of the red color corresponds to the TPM value. Expression of selected genes in bulk RNAseq (this work) shown as bar graphs (each dot represents a biological replicate in TPM, bars represent mean ± SEM) and single-cell RNAseq (downloaded from www.flyrnai.org/scRNA/blood/ and www.flyrnai.org/tools/single_cell/web/) shown by dot plots with average gene expressions in hemocyte clusters (color gradient of the dot represents the expression level, the size represents percentage of cells expressing the gene per cluster) and t-Distributed Stochastic Neighbor Embedding (t-SNE) plots of Harmony-based batch correction of wasp infected 48 hours data sets (for comparison, plots with plasmatocytes marker *Hml* and lamellocyte marker *Atilla* are shown).

