## Supplementary figures and images for "Metabolism of glucose and trehalose by cyclic pentose phosphate pathway is essential for effective immune response in *Drosophila*"

### S1 Figure

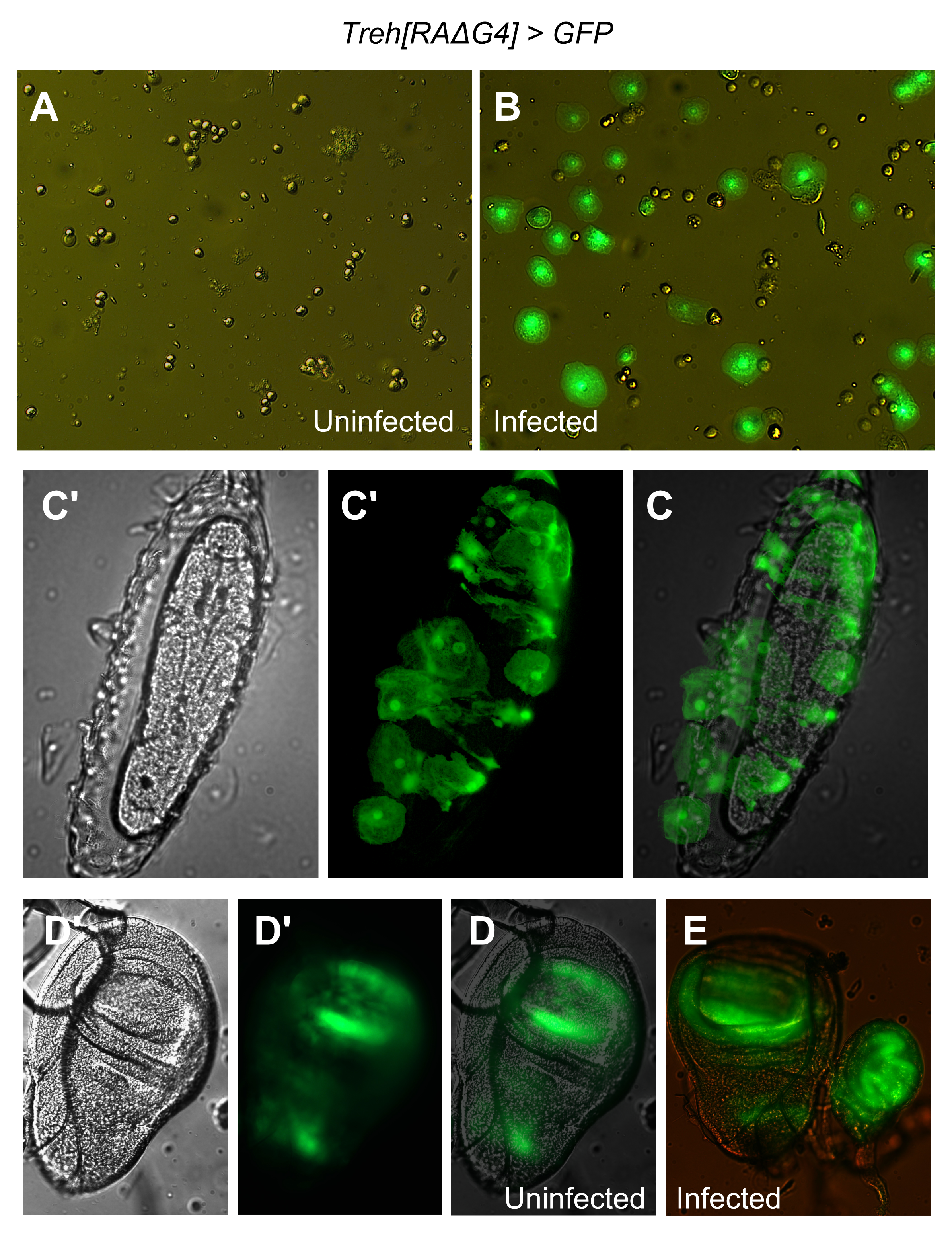

### S2 Figure

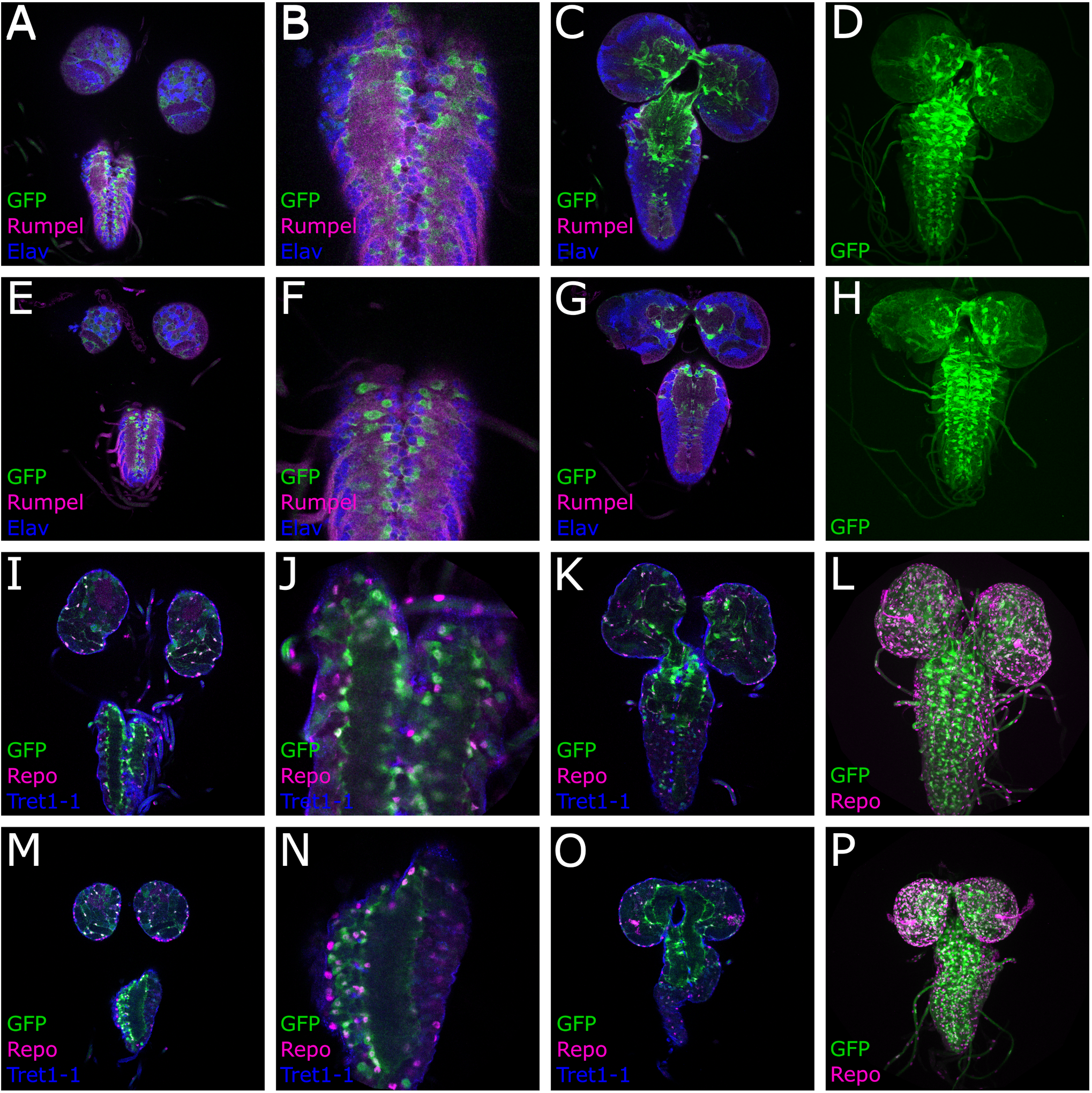

### S3 Figure

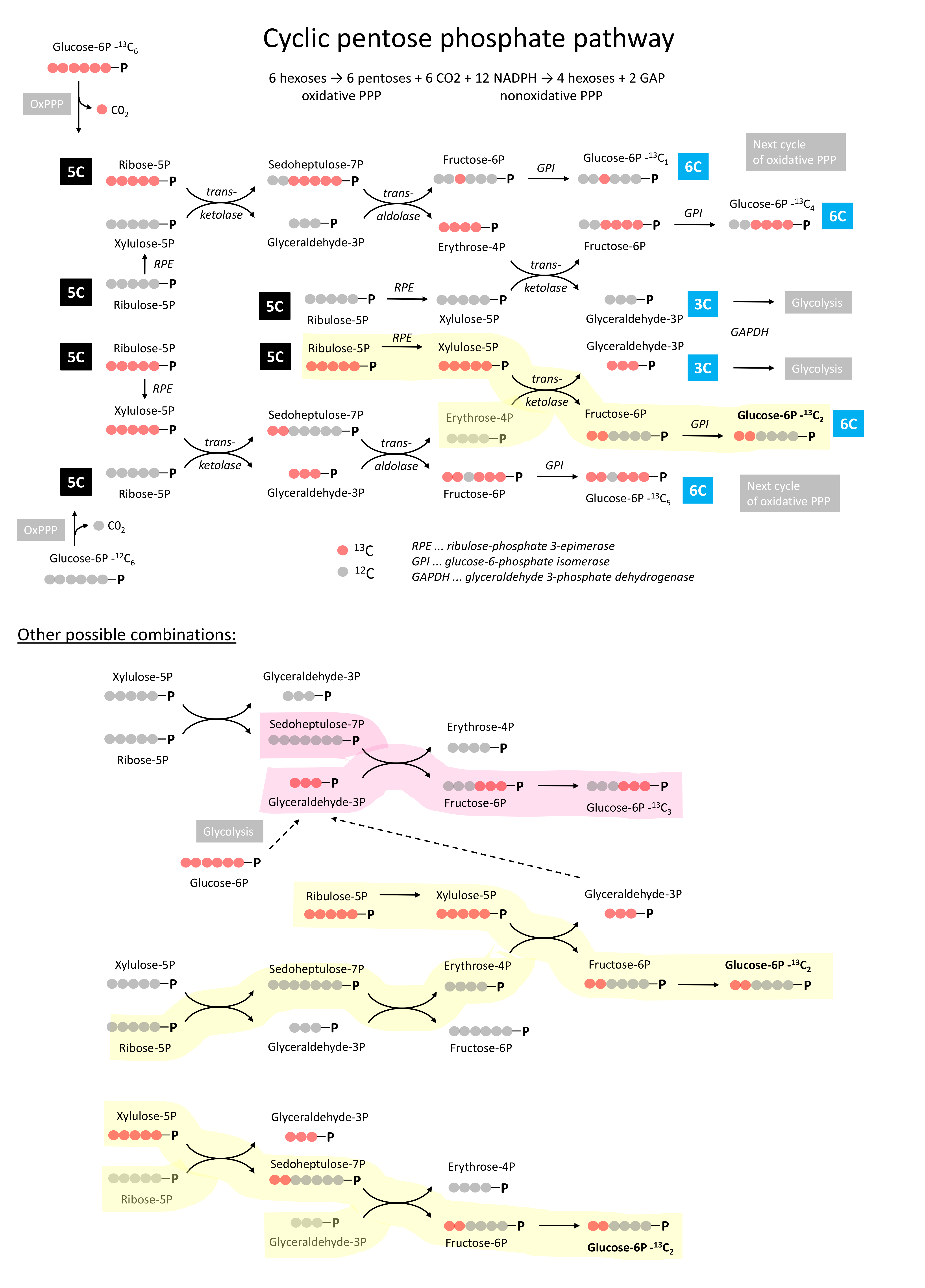

### S4 Figure

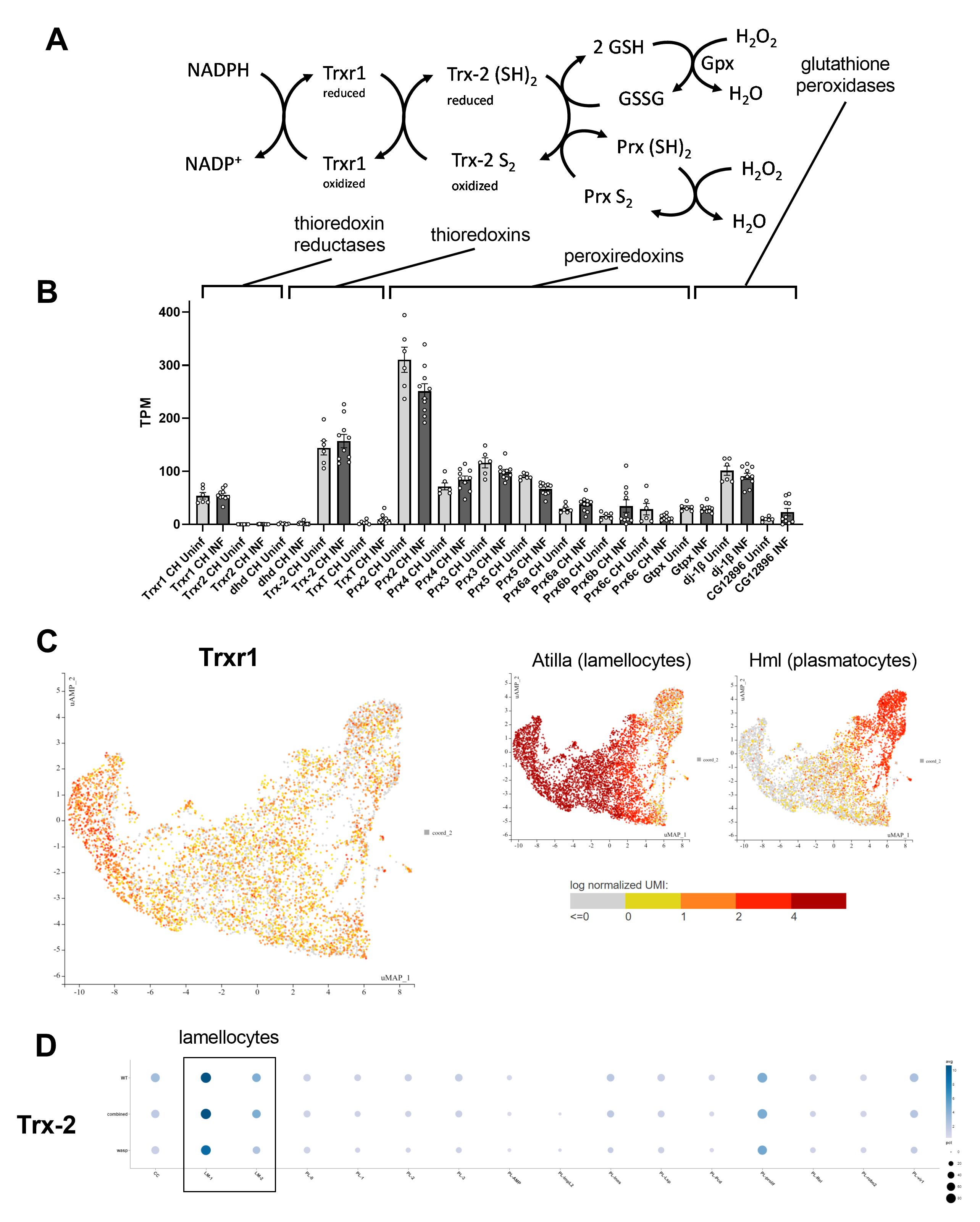

### S5 Figure

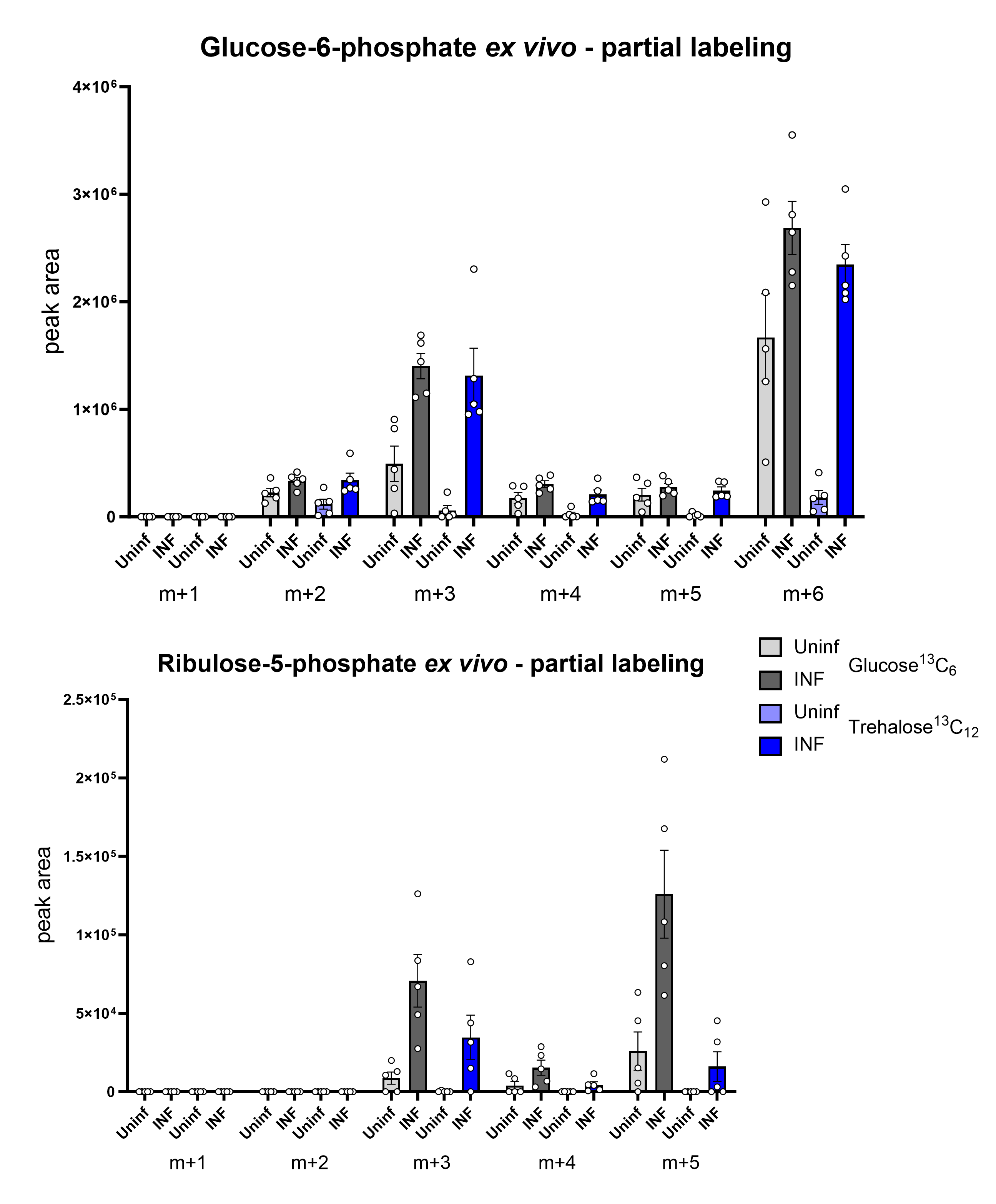

### S6 Figure

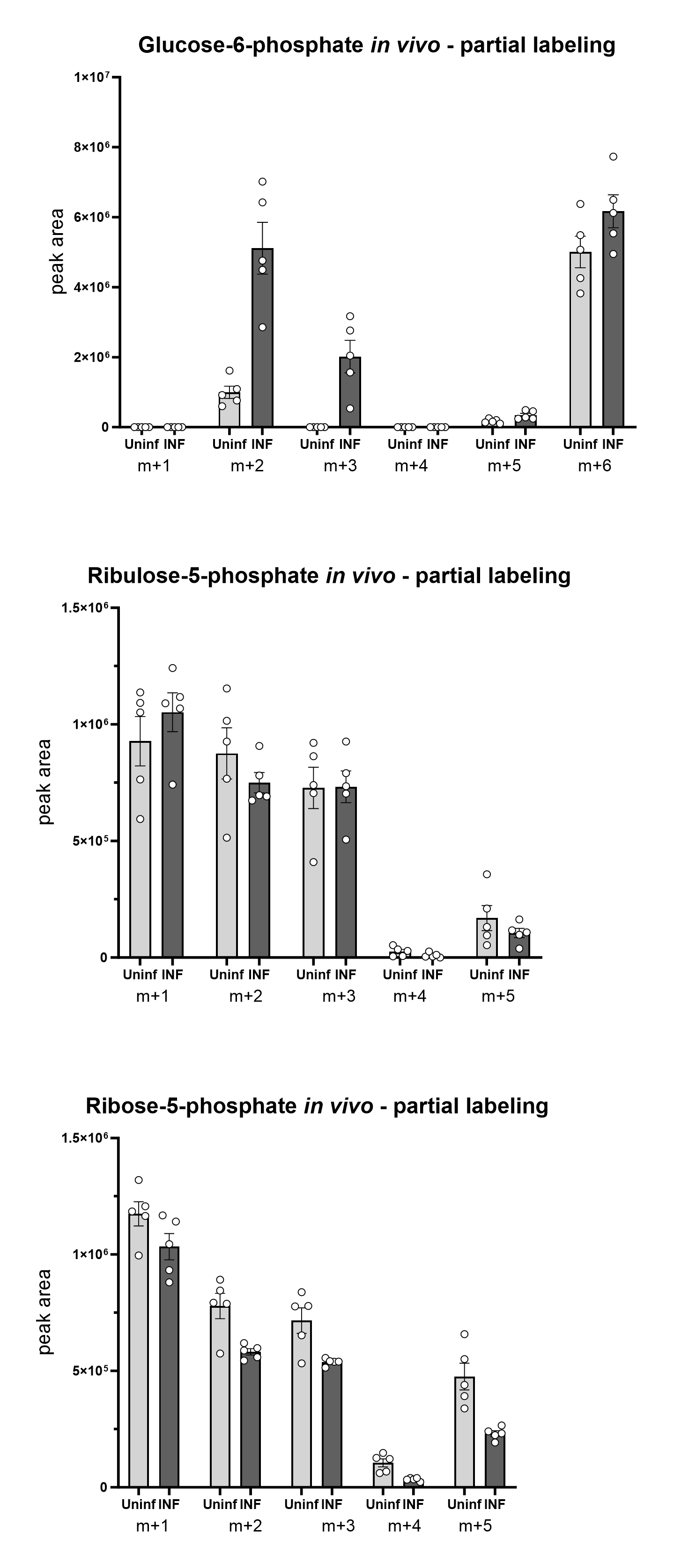

### S7 Figure

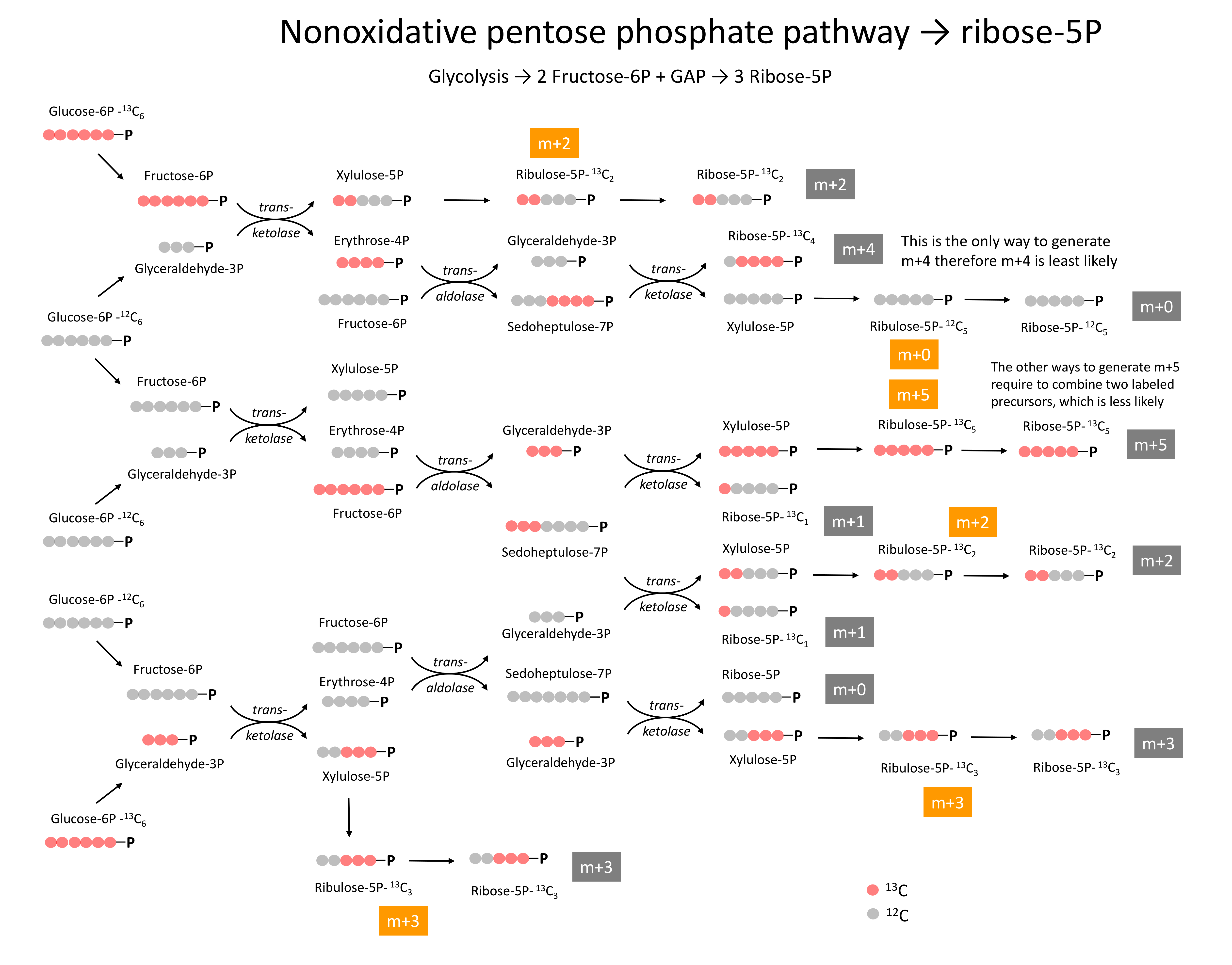

### S8 Figure

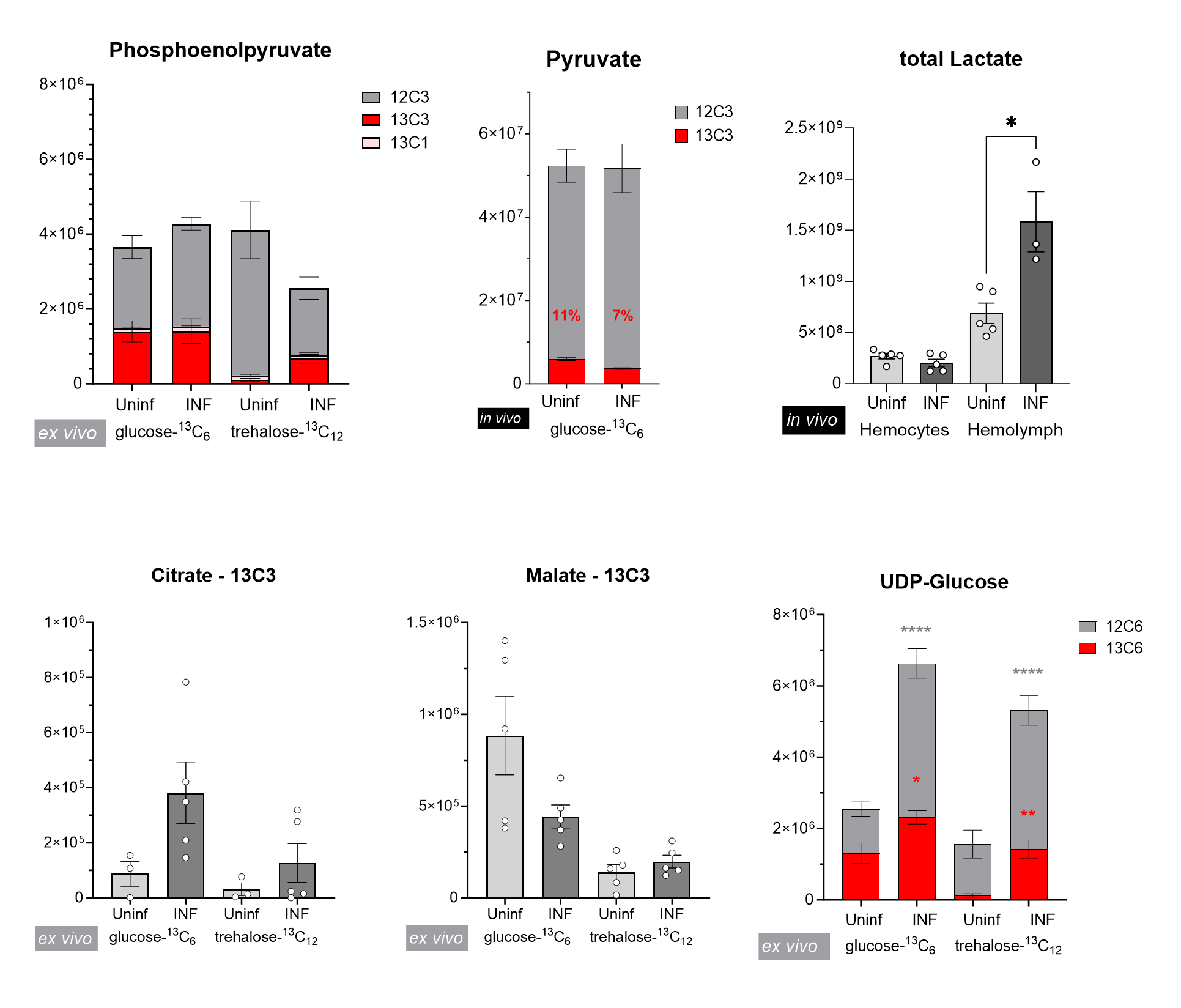

### S9 Figure

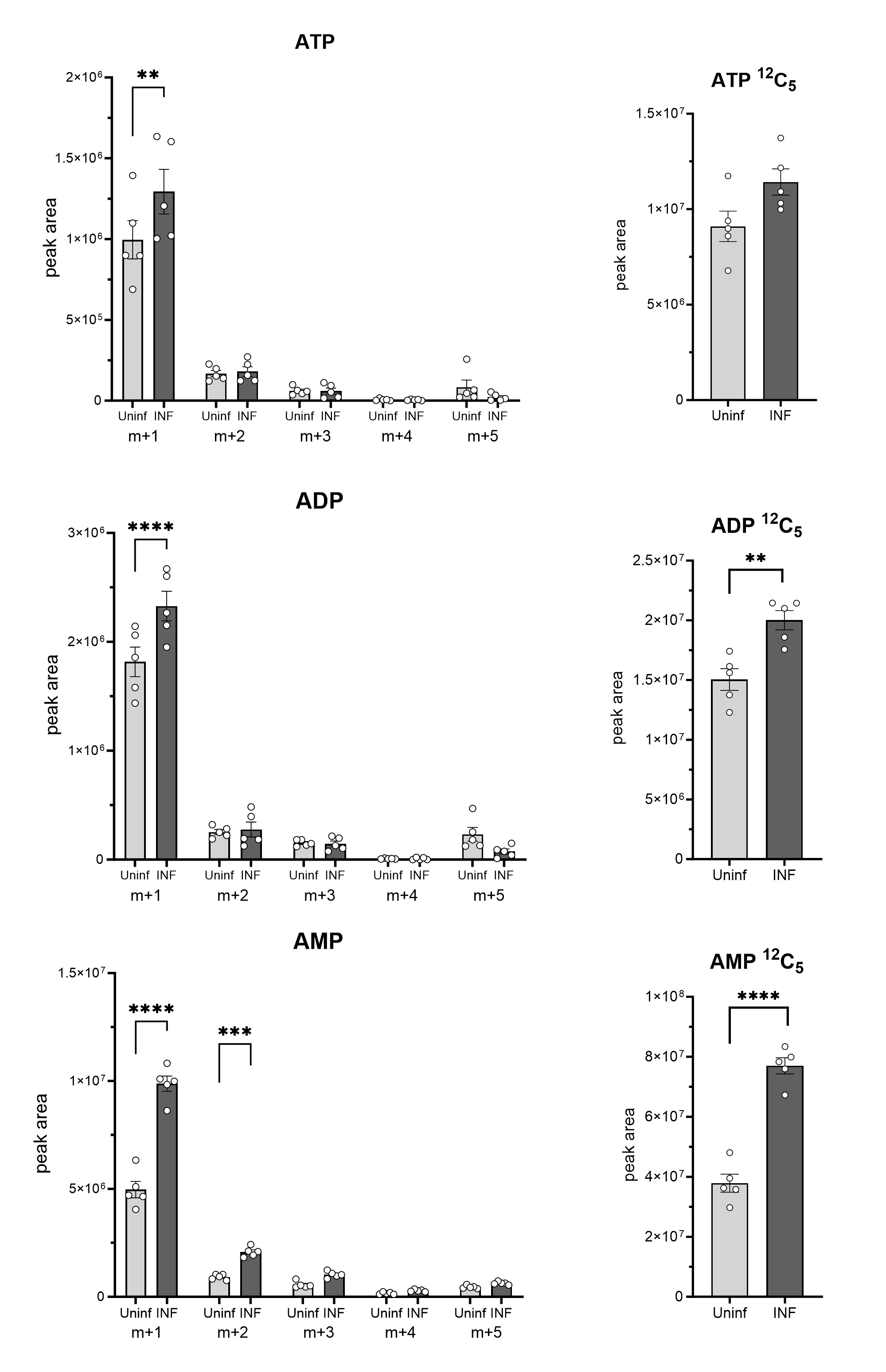

### S10 Figure

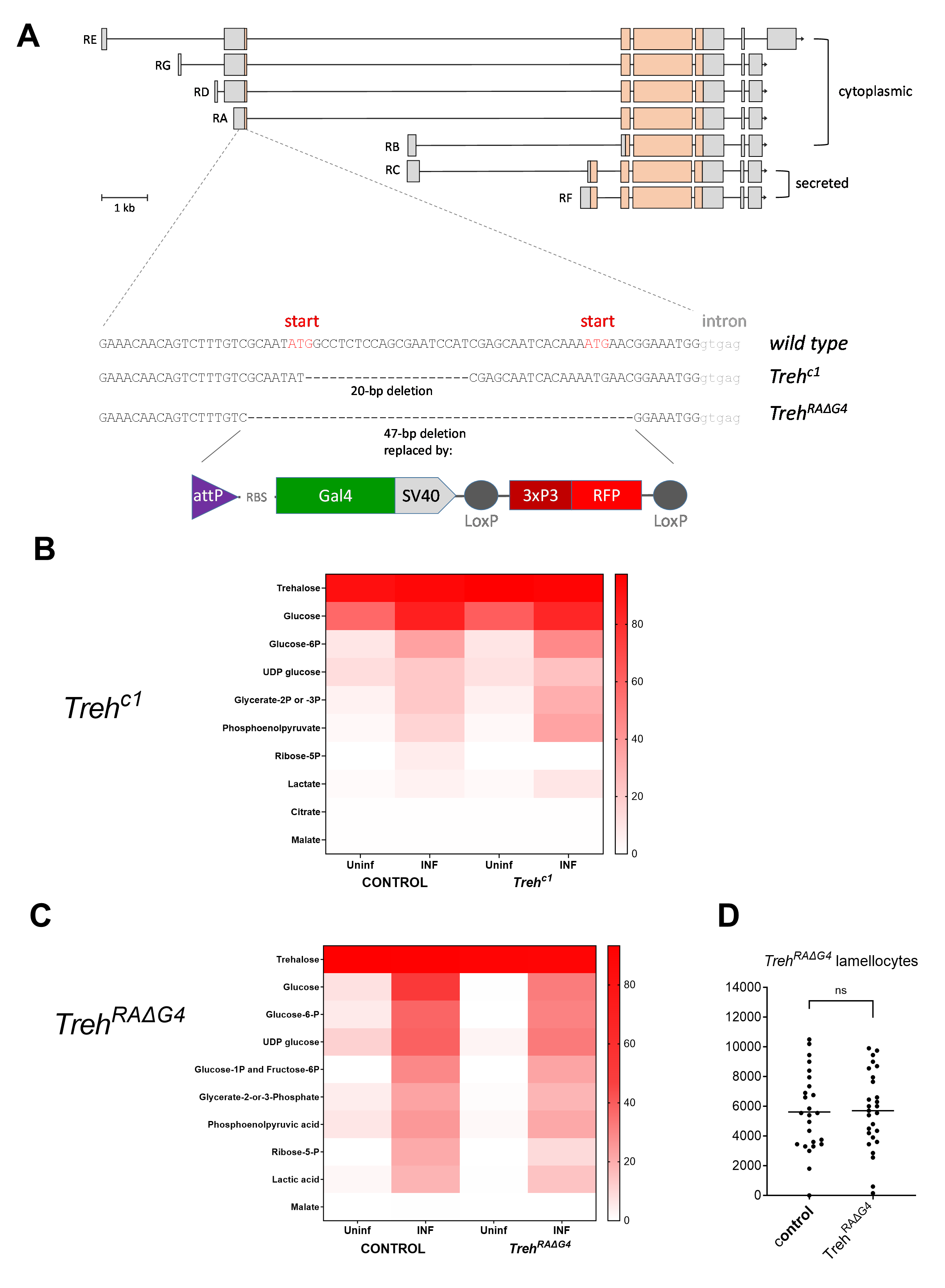

### S11 Figure

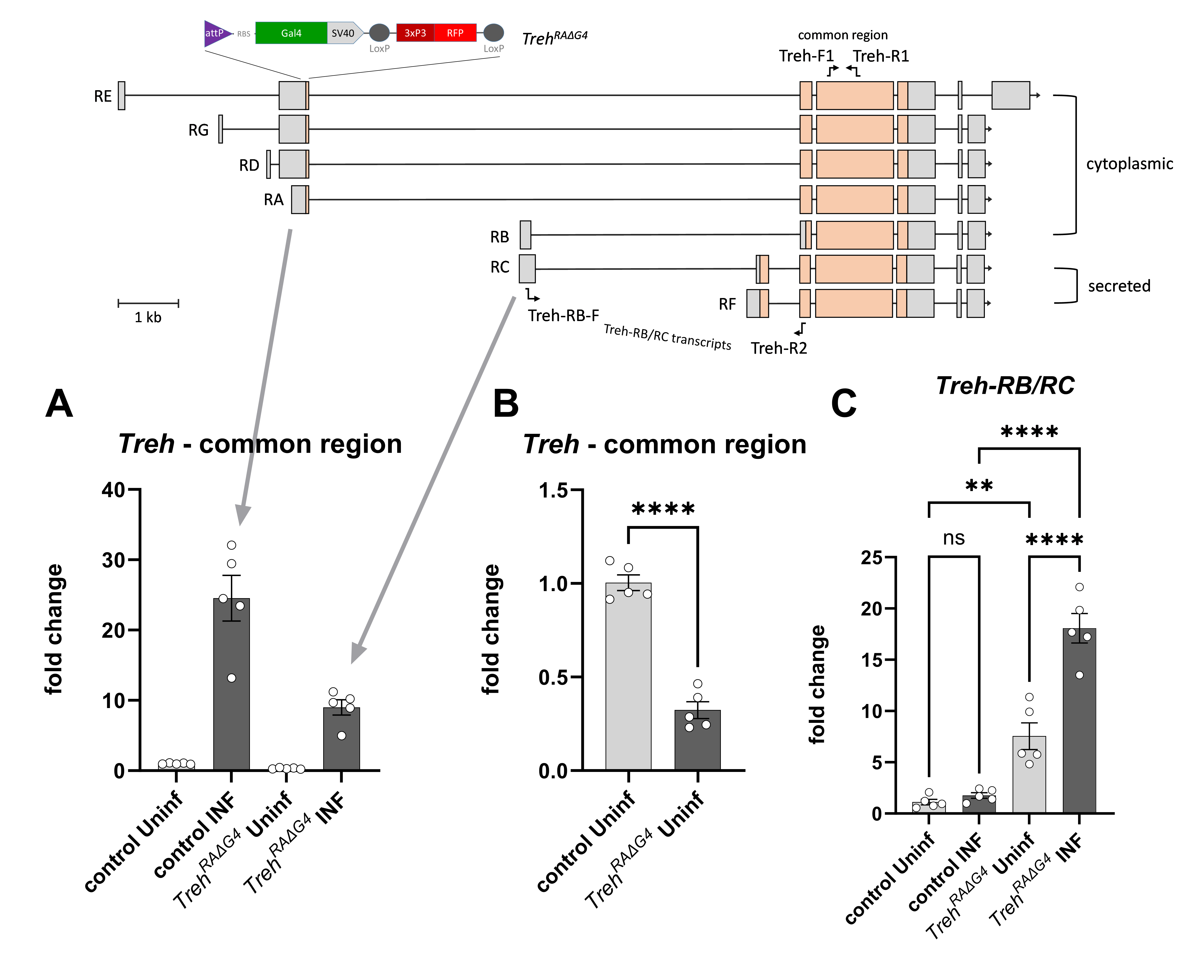

### S12 Figure

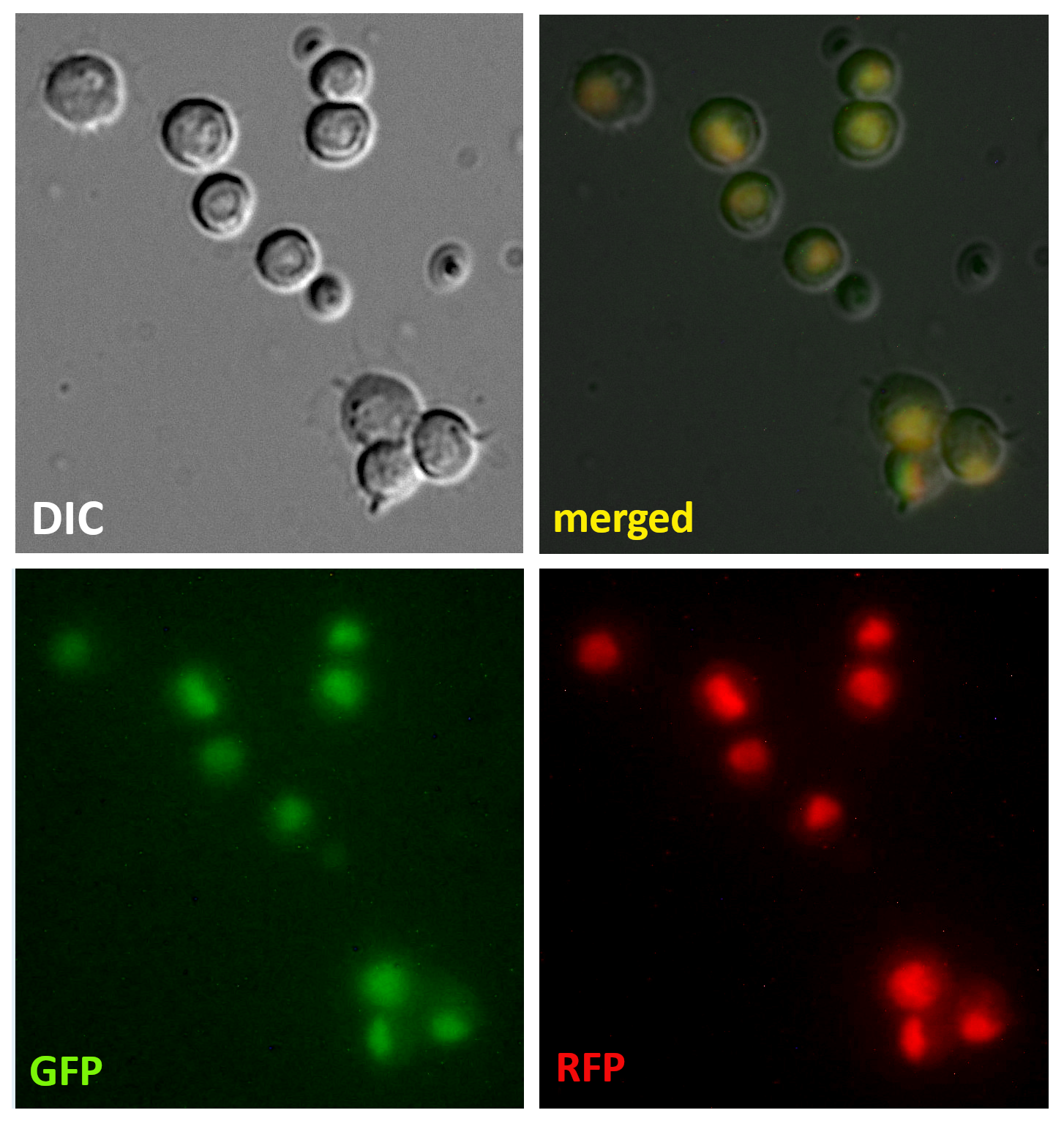
