## Supplementary material for "Metabolism of glucose and trehalose by cyclic pentose phosphate pathway is essential for effective immune response in *Drosophila*": S1 file

**S1 File** – Carbohydrate transport and metabolism - gene expression analysis by bulk and single-cell RNAseq

SLC2 and SLC17 family of hexose sugar transporters in *Drosophila*

### Bulk RNAseq [TPM]

| | CIRCULATING HEMOCYTES | | | | | LYMPH GLAND | | | | WING DISC | |
|---|---|---|---|---|---|---|---|---|---|---|---|
| GENE | 0 h | 9 h Uninf | 9 h INF | 18 h Uninf | 18 h INF | 9 h Uninf | 9 h INF | 18 h Uninf | 18 h INF | 9 h Uninf | 9 h INF |
| *MFS3* | 40.4 | 37.9 | 23.9 | 36.8 | 24.9 | 8.6 | 5.4 | 10.4 | 7.2 | 22.3 | 21.6 |
| *sut1* | 21.6 | 21.8 | 17.0 | 20.7 | 18.2 | 24.9 | 18.5 | 31.0 | 21.0 | 16.8 | 20.0 |
| *CG33281* | 14.8 | 10.8 | 1.8 | 9.2 | 4.9 | 5.1 | 1.8 | 14.8 | 10.6 | 9.2 | 7.1 |
| *CG7882* | 9.5 | 7.6 | 1.9 | 5.0 | 2.9 | 2.6 | 0.6 | 1.9 | 3.0 | 2.3 | 1.5 |
| *CG4607* | 7.8 | 4.9 | 30.0 | 10.3 | 36.6 | 1.5 | 2.4 | 2.2 | 3.3 | 1.1 | 1.5 |
| *Tret1-1* | 6.5 | 6.0 | 28.5 | 9.0 | 37.7 | 1.4 | 2.9 | 1.6 | 7.1 | 2.2 | 3.3 |
| *CG15408* | 5.9 | 3.8 | 1.5 | 3.5 | 1.2 | 1.0 | 0.3 | 5.2 | 3.7 | 3.1 | 2.6 |
| *CG6484* | 5.6 | 3.8 | 0.5 | 4.1 | 0.8 | 0.4 | 0.7 | 0.2 | 0.4 | 0.7 | 2.5 |
| *CG3285* | 4.2 | 1.9 | 0.9 | 2.5 | 0.7 | 0.3 | 0.1 | 2.4 | 2.6 | 2.4 | 1.8 |
| *CG1208* | 3.2 | 2.6 | 153.7 | 14.1 | 200.3 | 0.8 | 0.7 | 1.5 | 22.1 | 0.2 | 0.6 |
| *CG17930* | 3.0 | 1.4 | 0.3 | 4.3 | 0.6 | 0.0 | 0.2 | 0.1 | 0.1 | 1.6 | 0.4 |
| *CG6901* | 2.9 | 1.8 | 0.3 | 4.2 | 0.8 | 0.0 | 0.3 | 0.1 | 0.1 | 1.1 | 0.6 |
| *CG15406* | 2.9 | 2.8 | 1.8 | 2.8 | 1.1 | 1.2 | 0.6 | 3.5 | 3.6 | 3.2 | 2.9 |
| *CG14606* | 2.8 | 1.9 | 0.3 | 1.4 | 0.6 | 0.4 | 0.3 | 0.8 | 1.5 | 0.5 | 0.9 |
| *nebu* | 2.4 | 2.7 | 3.3 | 2.7 | 3.8 | 5.3 | 4.5 | 3.9 | 4.6 | 1.1 | 2.9 |
| *CG31100* | 2.3 | 1.7 | 1.1 | 1.6 | 1.3 | 0.2 | 0.3 | 0.2 | 0.2 | 0.5 | 0.5 |
| *pippin* | 1.7 | 2.0 | 3.0 | 2.0 | 2.0 | 1.2 | 0.4 | 2.2 | 0.5 | 2.0 | 1.3 |
| *CG14605* | 1.6 | 2.1 | 5.1 | 0.7 | 1.5 | 0.0 | 0.0 | 0.0 | 0.1 | 0.0 | 0.0 |
| *CG8837* | 1.5 | 1.3 | 0.4 | 1.1 | 0.7 | 2.0 | 2.3 | 3.4 | 7.6 | 1.9 | 1.1 |
| *CG8249* | 1.5 | 1.0 | 0.2 | 0.8 | 0.3 | 0.0 | 0.1 | 0.1 | 0.2 | 0.3 | 0.3 |
| *CG32054* | 1.4 | 1.1 | 0.2 | 1.1 | 0.2 | 0.0 | 0.1 | 0.0 | 0.1 | 0.5 | 0.4 |
| *CG32053* | 1.3 | 0.8 | 0.3 | 1.5 | 0.2 | 0.0 | 0.0 | 0.1 | 0.0 | 0.3 | 0.5 |
| *CG1213* | 1.1 | 0.8 | 1.4 | 1.2 | 1.6 | 2.4 | 3.1 | 3.0 | 4.5 | 0.1 | 0.2 |
| *CG42825* | 0.8 | 0.6 | 0.2 | 0.9 | 0.0 | 0.0 | 0.0 | 0.0 | 0.0 | 0.4 | 0.4 |
| *sut2* | 0.4 | 0.3 | 0.3 | 0.3 | 0.2 | 0.3 | 0.1 | 0.3 | 0.2 | 0.4 | 0.2 |
| *Glut1* | 0.4 | 0.3 | 0.4 | 0.3 | 0.4 | 0.5 | 0.6 | 0.7 | 0.6 | 0.5 | 0.4 |
| *sut4* | 0.4 | 0.4 | 1.1 | 0.1 | 0.4 | 0.0 | 0.0 | 0.1 | 0.3 | 0.0 | 0.0 |
| *sut3* | 0.3 | 0.5 | 0.9 | 0.2 | 0.4 | 0.0 | 0.0 | 0.0 | 0.0 | 0.0 | 0.0 |
| *CG33282* | 0.3 | 0.3 | 0.1 | 0.2 | 0.1 | 0.0 | 0.0 | 0.0 | 0.1 | 0.1 | 0.0 |
| *Glut3* | 0.2 | 0.3 | 0.6 | 0.0 | 0.1 | 0.0 | 0.0 | 0.1 | 0.0 | 0.0 | 0.0 |
| *Tret1-2* | 0.1 | 0.3 | 0.2 | 0.1 | 0.1 | 0.1 | 0.1 | 0.1 | 0.1 | 0.7 | 0.2 |
| *CG17929* | 0.1 | 0.0 | 0.1 | 0.1 | 0.0 | 0.1 | 0.2 | 0.0 | 0.4 | 0.1 | 0.1 |

### Bulk RNAseq [TPM]

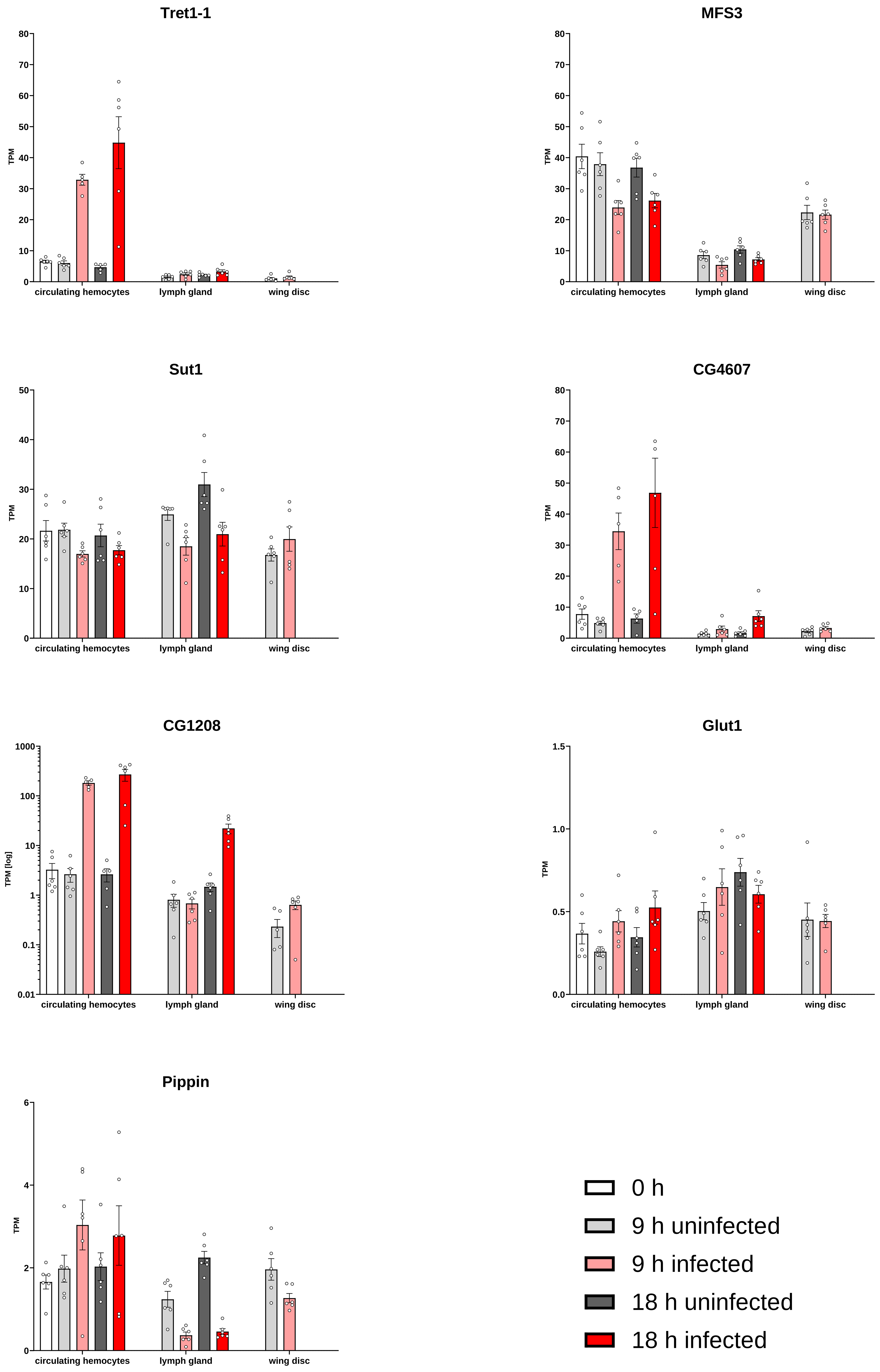

### Single cell-RNAseq of circulating hemocytes

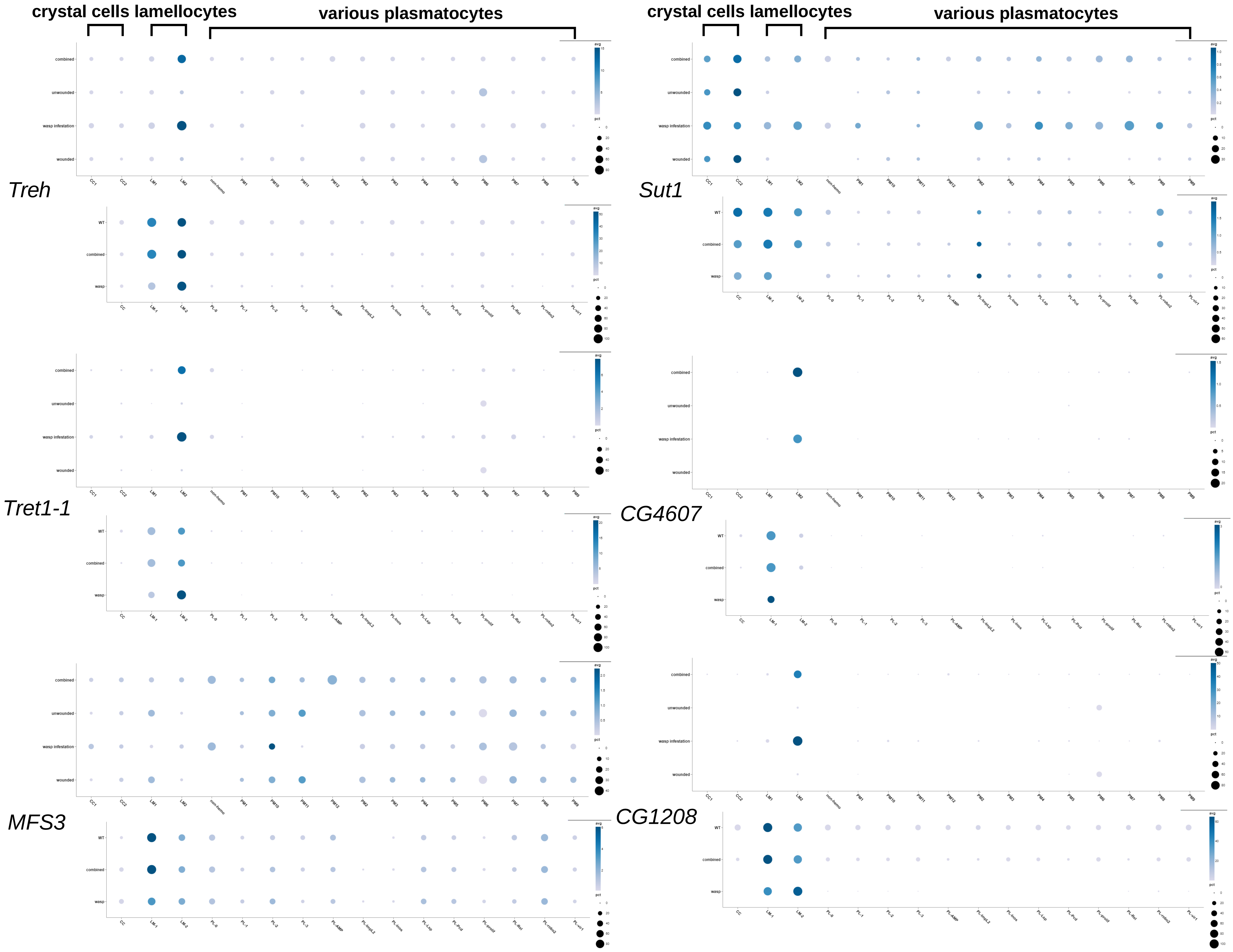


SOURCE: https://www.flyrnai.org/tools/single_cell/web/

Cattenoz PB, Sakr R, Pavlidaki A, Delaporte C, Riba A, Molina N, Hariharan N, Mukherjee T, Giangrande A. Temporal specificity and heterogeneity of Drosophila immune cells. EMBO J. 2020 Jun 17;39(12):e104486. doi: 10.15252/embj.2020104486.

Tattikota SG, Cho B, Liu Y, Hu Y, Barrera V, Steinbaugh MJ, Yoon SH, Comjean A, Li F, Dervis F, Hung RJ, Nam JW, Ho Sui S, Shim J, Perrimon N. A single-cell survey of *Drosophila* blood. Elife. 2020 May 12;9:e54818. doi: 10.7554/eLife.54818.

### Single cell-RNAseq of circulating hemocytes – 48 hours wasp infected

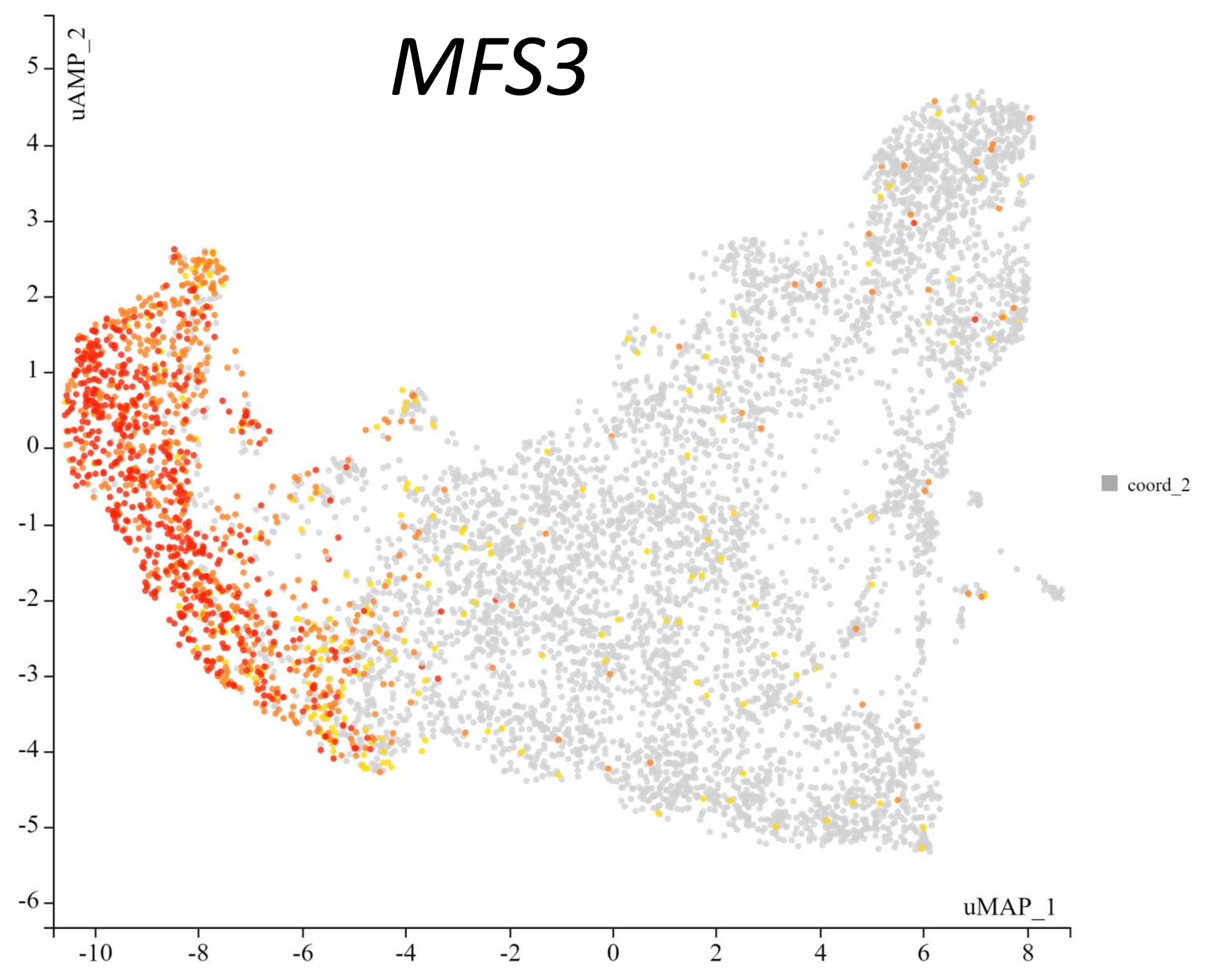


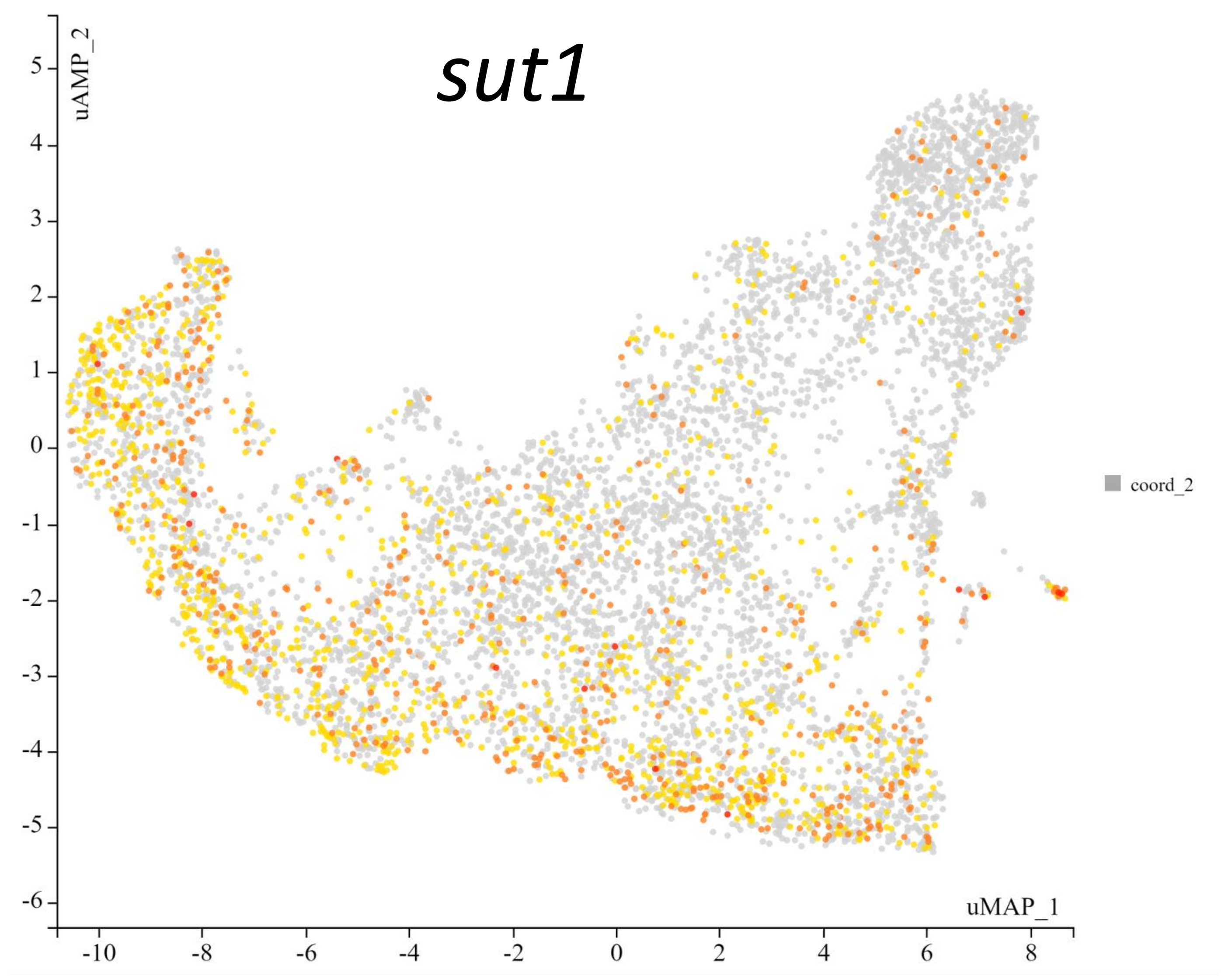


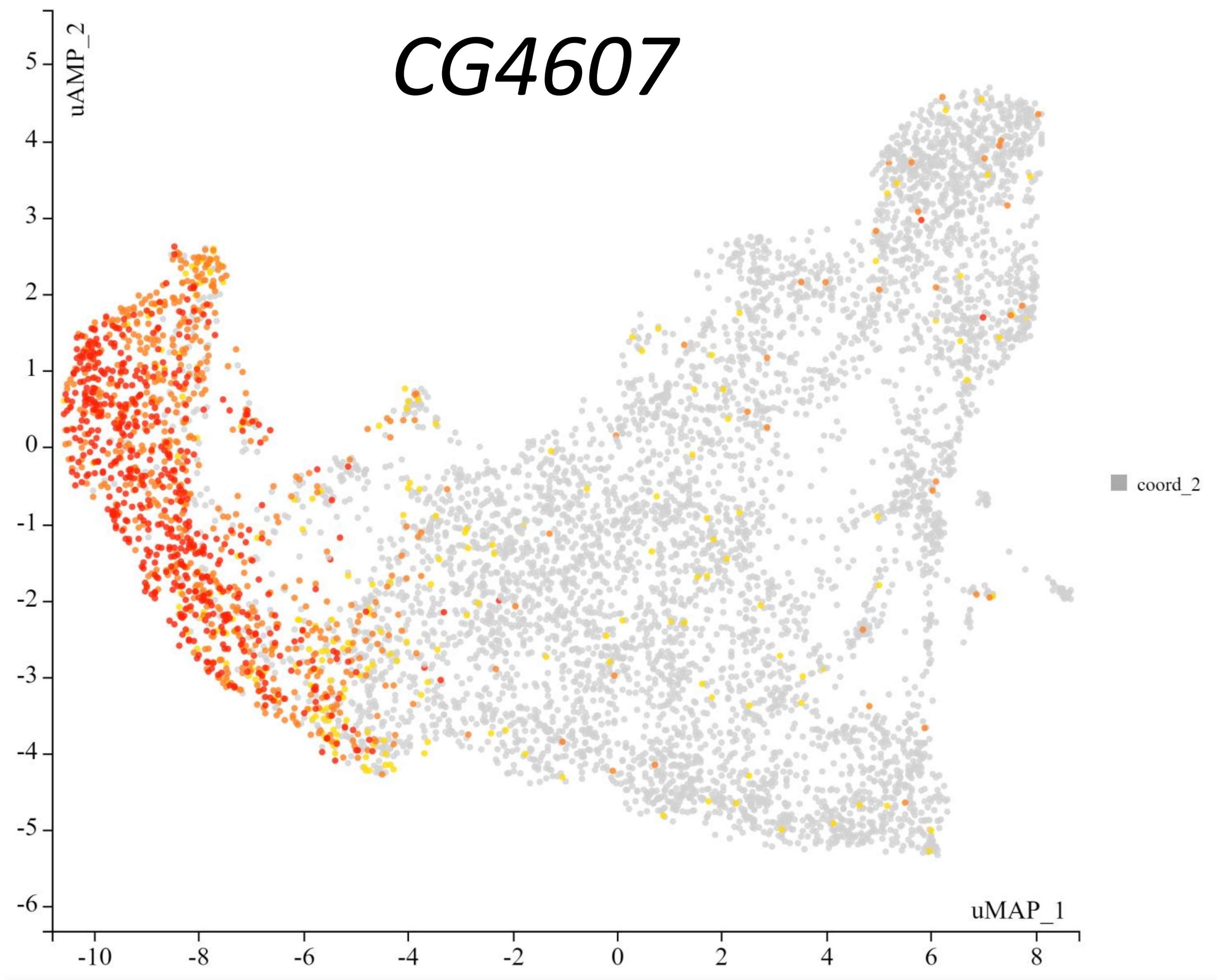


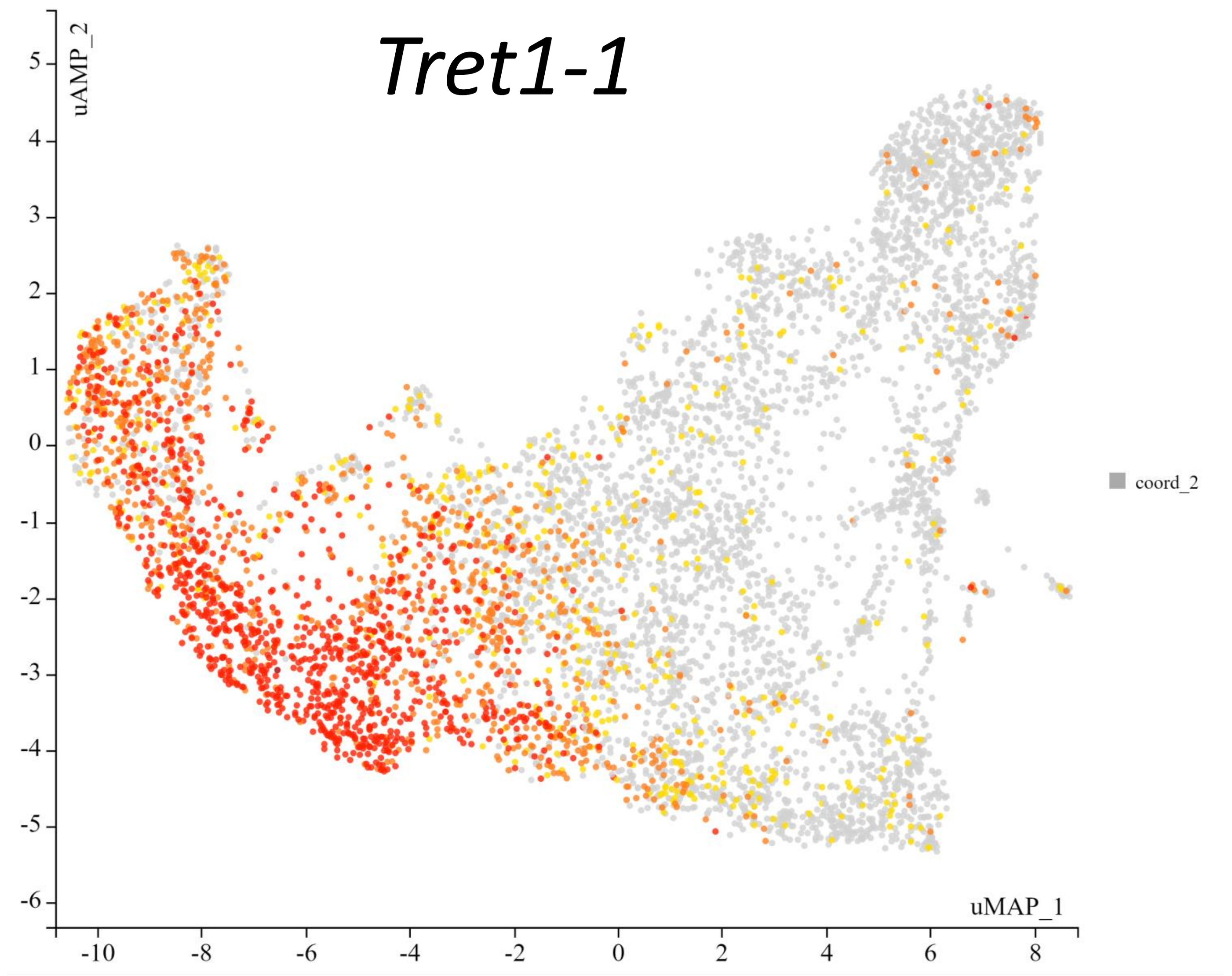


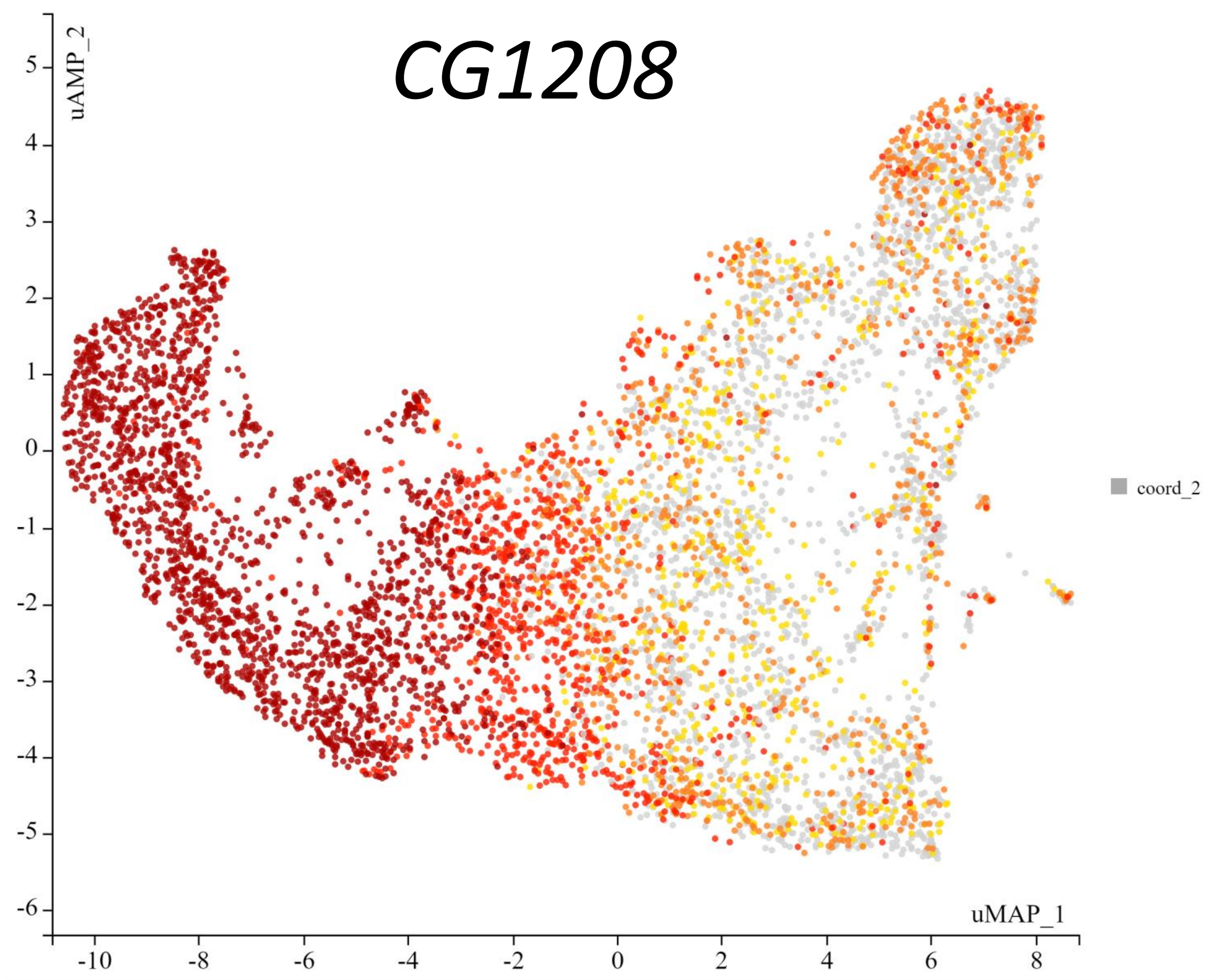


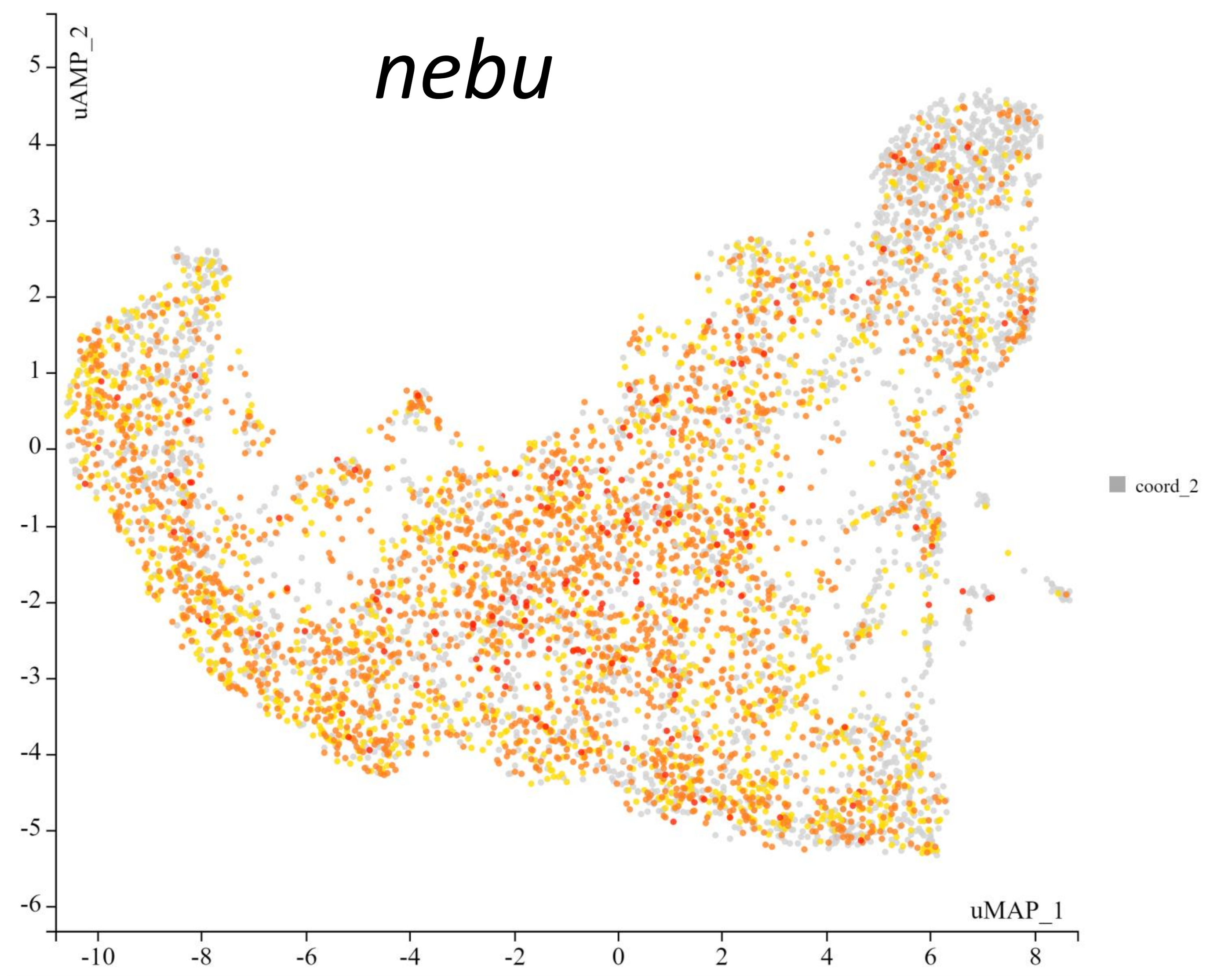


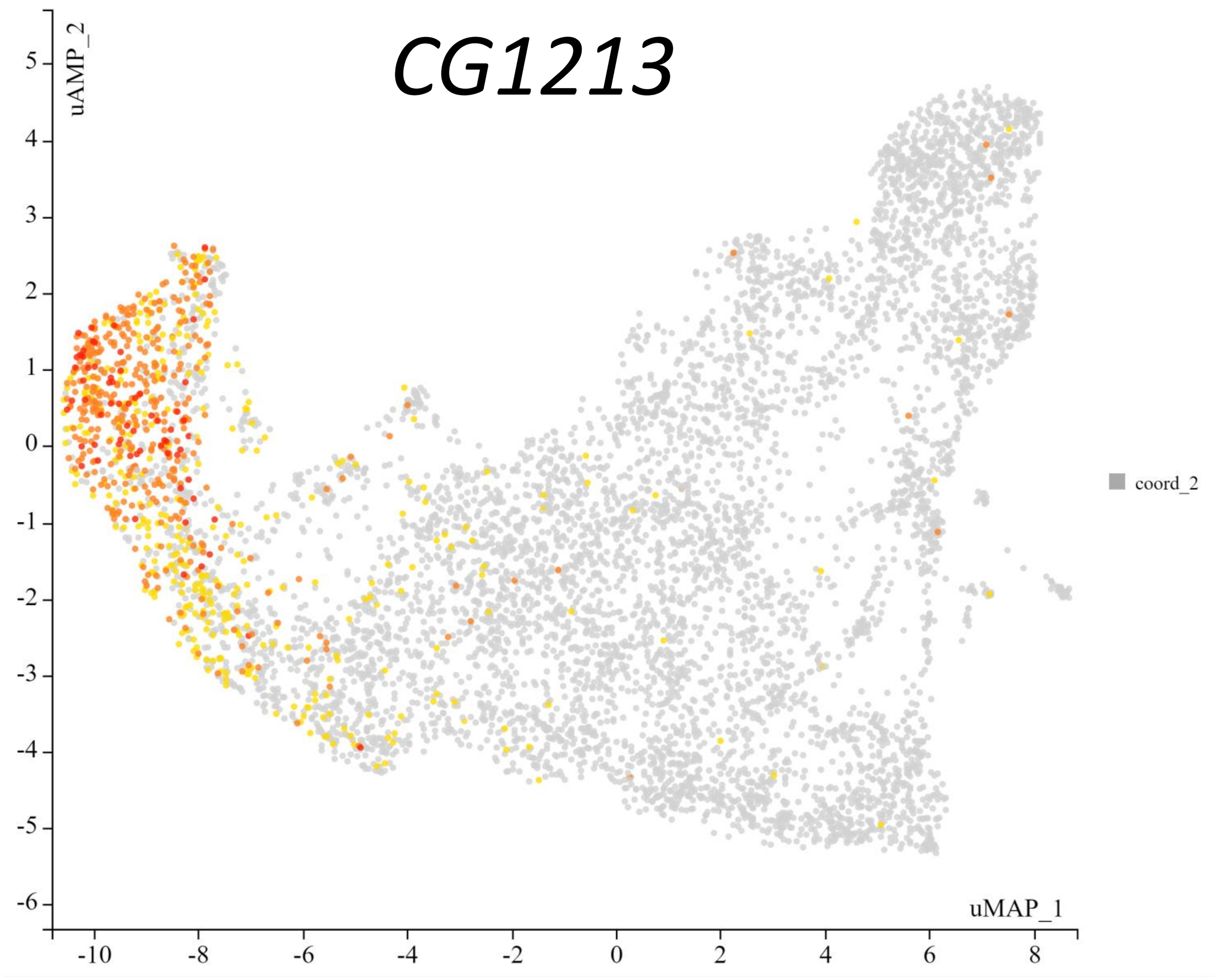


*Atilla (lamellocytes)*

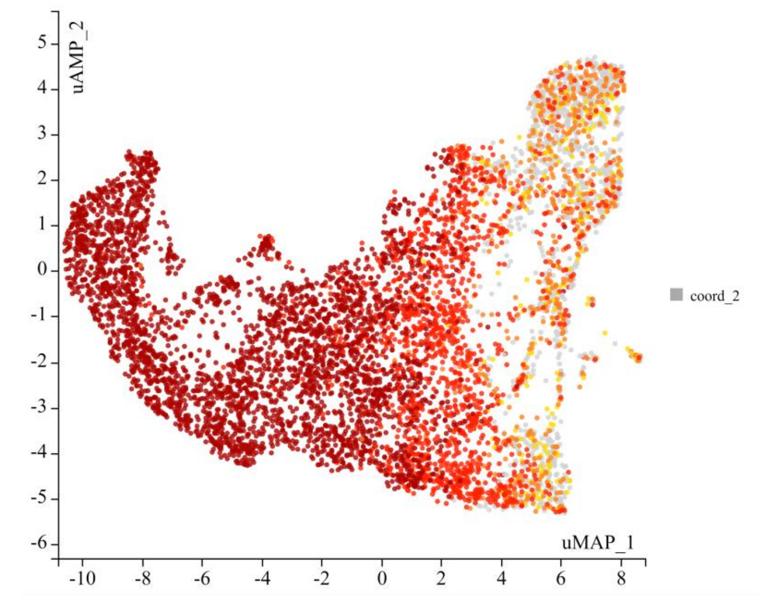

*Hml (plasmatocytes)*

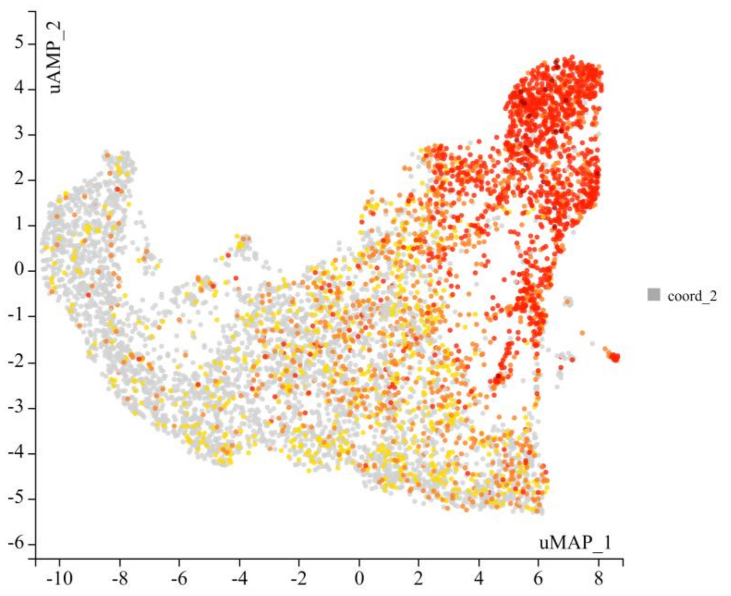

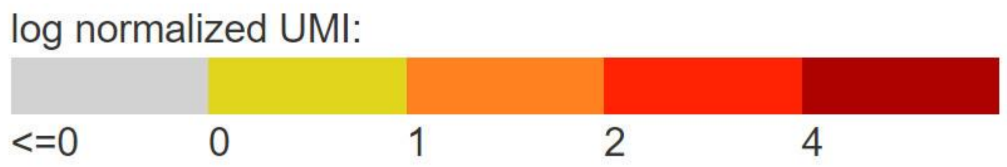


Source: https://www.flyrnai.org/scRNA/blood/

Tattikota SG, Cho B, Liu Y, Hu Y, Barrera V, Steinbaugh MJ, Yoon SH, Comjean A, Li F, Dervis F, Hung RJ, Nam JW, Ho Sui S, Shim J, Perrimon N. A single-cell survey of *Drosophila* blood. Elife. 2020 May 12;9:e54818. doi: 10.7554/eLife.54818.
