## Supplementary material for "Metabolism of glucose and trehalose by cyclic pentose phosphate pathway is essential for effective immune response in *Drosophila*": S2 File

### **S2 File** – Glycolytic and PPP gene expression analysis by bulk and single-cell RNAseq

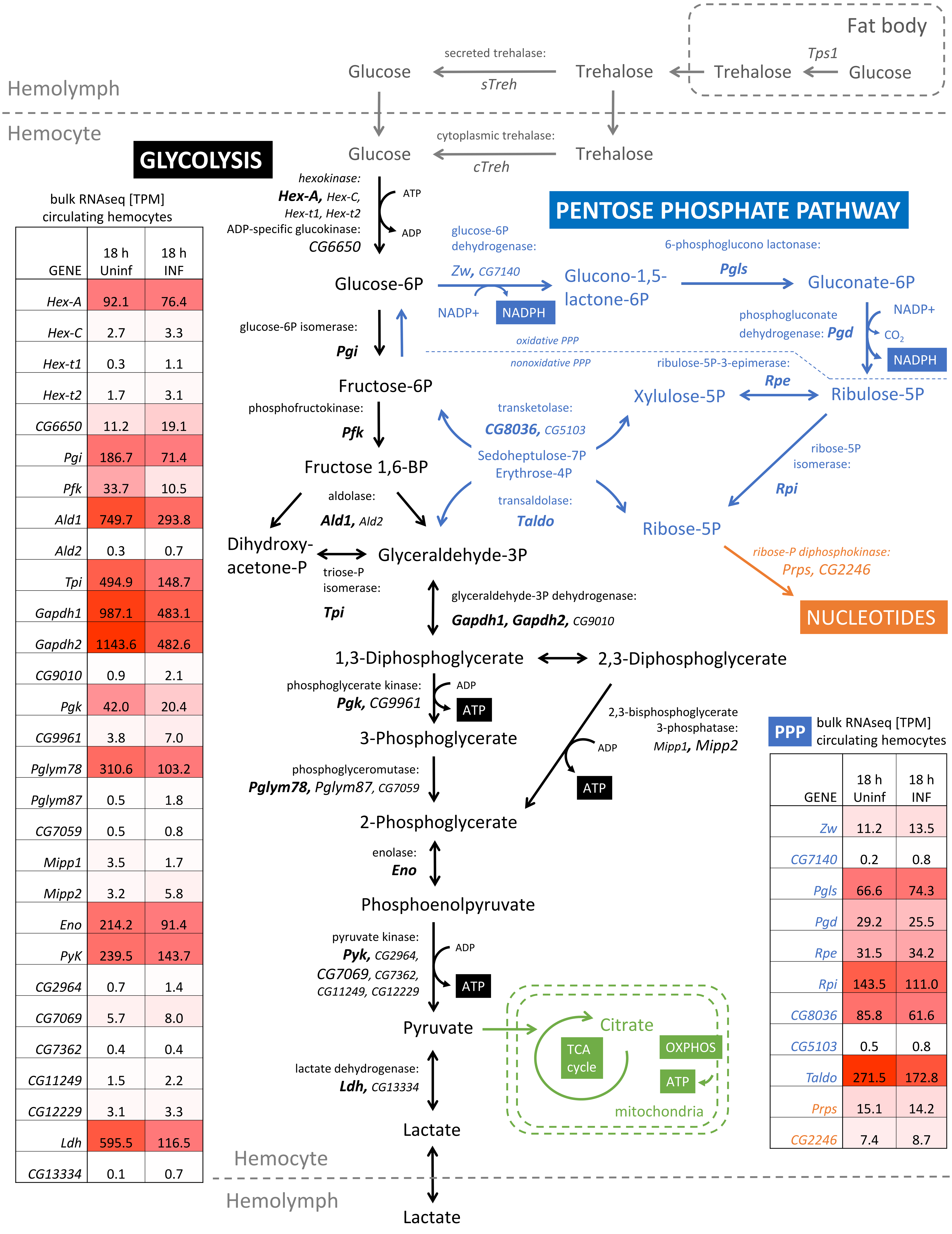


GLYCOLYSIS

bulk RNAseq [TPM]
circulating hemocytes

| GENE | 18 h Uninf | 18 h INF |
|---|---|---|
| *Hex-A* | 92.1 | 76.4 |
| *Hex-C* | 2.7 | 3.3 |
| *Hex-t1* | 0.3 | 1.1 |
| *Hex-t2* | 1.7 | 3.1 |
| *CG6650* | 11.2 | 19.1 |
| *Pgi* | 186.7 | 71.4 |
| *Pfk* | 33.7 | 10.5 |
| *Ald1* | 749.7 | 293.8 |
| *Ald2* | 0.3 | 0.7 |
| *Tpi* | 494.9 | 148.7 |
| *Gapdh1* | 987.1 | 483.1 |
| *Gapdh2* | 1143.6 | 482.6 |
| *CG9010* | 0.9 | 2.1 |
| *Pgk* | 42.0 | 20.4 |
| *CG9961* | 3.8 | 7.0 |
| *Pglym78* | 310.6 | 103.2 |
| *Pglym87* | 0.5 | 1.8 |
| *CG7059* | 0.5 | 0.8 |
| *Mipp1* | 3.5 | 1.7 |
| *Mipp2* | 3.2 | 5.8 |
| *Eno* | 214.2 | 91.4 |
| *PyK* | 239.5 | 143.7 |
| *CG2964* | 0.7 | 1.4 |
| *CG7069* | 5.7 | 8.0 |
| *CG7362* | 0.4 | 0.4 |
| *CG11249* | 1.5 | 2.2 |
| *CG12229* | 3.1 | 3.3 |
| *Ldh* | 595.5 | 116.5 |
| *CG13334* | 0.1 | 0.7 |

PPP bulk RNAseq [TPM]
circulating hemocytes

| GENE | 18 h Uninf | 18 h INF |
|---|---|---|
| *Zw* | 11.2 | 13.5 |
| *CG7140* | 0.2 | 0.8 |
| *Pgls* | 66.6 | 74.3 |
| *Pgd* | 29.2 | 25.5 |
| *Rpe* | 31.5 | 34.2 |
| *Rpi* | 143.5 | 111.0 |
| *CG8036* | 85.8 | 61.6 |
| *CG5103* | 0.5 | 0.8 |
| *Taldo* | 271.5 | 172.8 |
| *Prps* | 15.1 | 14.2 |
| *CG2246* | 7.4 | 8.7 |

### Bulk RNAseq [TPM]

#### Glycolysis

| | CIRCULATING HEMOCYTES | | | | | LYMPH GLAND | | | | WING DISC | |
|---|---|---|---|---|---|---|---|---|---|---|---|
| GENE | 0 h | 9 h Uninf | 9 h INF | 18 h Uninf | 18 h INF | 9 h Uninf | 9 h INF | 18 h Uninf | 18 h INF | 9 h Uninf | 9 h INF |
| *Hex-A* | 112.0 | 101.7 | 68.8 | 92.1 | 76.4 | 80.2 | 76.0 | 101.6 | 123.9 | 84.6 | 95.0 |
| *Hex-C* | 8.5 | 4.3 | 1.1 | 2.7 | 3.3 | 1.8 | 2.9 | 0.5 | 1.6 | 1.3 | 1.4 |
| *Hex-t1* | 0.4 | 0.7 | 1.8 | 0.3 | 1.1 | 0.0 | 0.0 | 0.0 | 0.1 | 0.0 | 0.0 |
| *Hex-t2* | 2.2 | 2.3 | 5.3 | 1.7 | 3.1 | 0.0 | 0.0 | 0.2 | 0.5 | 0.1 | 0.1 |
| *CG6650* | 12.4 | 13.8 | 23.6 | 11.2 | 19.1 | 17.8 | 13.3 | 18.4 | 15.4 | 22.7 | 24.2 |
| *Pgi* | 231.5 | 232.9 | 50.6 | 186.7 | 71.4 | 56.3 | 39.2 | 57.8 | 54.9 | 101.7 | 73.3 |
| *Pfk* | 34.7 | 36.8 | 7.2 | 33.7 | 10.5 | 5.6 | 3.6 | 7.9 | 5.8 | 14.6 | 11.2 |
| *Ald1* | 707.1 | 822.5 | 200.2 | 749.7 | 293.8 | 227.3 | 160.5 | 278.0 | 219.9 | 280.1 | 284.9 |
| *Ald2* | 0.8 | 1.2 | 1.5 | 0.3 | 0.7 | 0.0 | 0.0 | 0.0 | 0.2 | 0.0 | 0.0 |
| *Tpi* | 543.2 | 526.1 | 96.5 | 494.9 | 148.7 | 128.4 | 81.0 | 195.3 | 152.5 | 290.0 | 265.6 |
| *Gapdh1* | 979.7 | 1106.3 | 436.8 | 987.1 | 483.1 | 386.1 | 318.8 | 436.3 | 442.5 | 608.8 | 592.1 |
| *Gapdh2* | 1106.1 | 1320.0 | 296.3 | 1143.6 | 482.6 | 428.8 | 261.3 | 477.5 | 388.0 | 831.9 | 668.2 |
| *CG9010* | 1.0 | 1.2 | 2.8 | 0.9 | 2.1 | 0.8 | 0.9 | 0.9 | 1.2 | 0.4 | 0.6 |
| *Pgk* | 49.8 | 46.0 | 17.1 | 42.0 | 20.4 | 20.1 | 15.2 | 26.9 | 19.5 | 32.1 | 27.5 |
| *CG9961* | 3.4 | 4.4 | 10.8 | 3.8 | 7.0 | 2.2 | 2.7 | 3.8 | 4.2 | 4.1 | 3.5 |
| *Pglym78* | 306.4 | 325.3 | 62.4 | 310.6 | 103.2 | 90.2 | 64.0 | 92.6 | 96.6 | 164.2 | 137.0 |
| *Pglym87* | 1.9 | 2.9 | 6.6 | 0.5 | 1.8 | 0.0 | 0.0 | 0.2 | 0.4 | 0.1 | 0.1 |
| *CG7059* | 0.9 | 1.4 | 1.9 | 0.5 | 0.8 | 0.2 | 0.4 | 0.4 | 0.8 | 1.4 | 2.8 |
| *Mipp1* | 4.4 | 3.0 | 0.9 | 3.5 | 1.7 | 2.0 | 1.5 | 1.2 | 1.4 | 0.9 | 1.0 |
| *Mipp2* | 3.4 | 3.1 | 6.6 | 3.2 | 5.8 | 4.0 | 5.0 | 4.6 | 4.6 | 4.1 | 4.4 |
| *Eno* | 233.9 | 219.5 | 83.5 | 214.2 | 91.4 | 104.8 | 90.0 | 114.8 | 104.7 | 180.0 | 140.5 |
| *PyK* | 289.5 | 269.2 | 125.1 | 239.5 | 143.7 | 77.6 | 73.6 | 80.5 | 84.0 | 109.0 | 96.2 |
| *CG2964* | 1.4 | 2.3 | 3.1 | 0.7 | 1.4 | 0.0 | 0.2 | 0.1 | 0.2 | 0.1 | 0.1 |
| *CG7069* | 7.7 | 8.2 | 14.9 | 5.7 | 8.0 | 1.7 | 1.6 | 2.0 | 2.1 | 2.6 | 1.8 |
| *CG7362* | 0.6 | 0.4 | 0.8 | 0.4 | 0.4 | 0.8 | 0.3 | 0.9 | 0.6 | 0.7 | 0.8 |
| *CG11249* | 1.5 | 1.8 | 2.9 | 1.5 | 2.2 | 1.4 | 1.1 | 1.8 | 1.4 | 1.7 | 1.6 |
| *CG12229* | 4.4 | 4.4 | 5.3 | 3.1 | 3.3 | 9.5 | 9.8 | 9.6 | 9.7 | 10.9 | 8.9 |
| *Ldh* | 526.5 | 544.8 | 87.2 | 595.5 | 116.5 | 49.1 | 20.3 | 50.4 | 45.7 | 118.3 | 100.1 |
| *CG13334* | 0.4 | 0.2 | 0.7 | 0.1 | 0.7 | 1.7 | 2.0 | 1.9 | 1.3 | 0.9 | 1.2 |

#### Pentose phosphate pathway

| | CIRCULATING HEMOCYTES | | | | | LYMPH GLAND | | | | WING DISC | |
|---|---|---|---|---|---|---|---|---|---|---|---|
| GENE | 0 h | 9 h Uninf | 9 h INF | 18 h Uninf | 18 h INF | 9 h Uninf | 9 h INF | 18 h Uninf | 18 h INF | 9 h Uninf | 9 h INF |
| *Zw* | 14.2 | 13.8 | 15.1 | 11.2 | 13.5 | 22.7 | 10.0 | 31.7 | 8.4 | 7.8 | 5.5 |
| *CG7140* | 0.3 | 0.9 | 0.9 | 0.2 | 0.8 | 0.0 | 0.0 | 0.0 | 0.2 | 0.0 | 0.0 |
| *Pgls* | 87.1 | 71.8 | 74.9 | 66.6 | 74.3 | 68.5 | 73.7 | 76.8 | 83.5 | 67.5 | 49.3 |
| *Pgd* | 37.8 | 39.9 | 19.4 | 29.2 | 25.5 | 32.9 | 37.0 | 25.4 | 30.7 | 30.4 | 21.1 |
| *Rpe* | 37.9 | 35.0 | 40.3 | 31.5 | 34.2 | 51.0 | 52.6 | 53.6 | 40.2 | 60.1 | 39.9 |
| *Rpi* | 168.4 | 161.6 | 108.2 | 143.5 | 111.0 | 170.6 | 179.6 | 186.9 | 145.4 | 192.7 | 149.2 |
| *CG8036* | 126.7 | 105.5 | 44.0 | 85.8 | 61.6 | 63.0 | 44.2 | 53.7 | 41.6 | 96.8 | 74.8 |
| *CG5103* | 0.9 | 1.0 | 1.9 | 0.5 | 0.8 | 0.0 | 0.0 | 0.0 | 0.1 | 0.0 | 0.0 |
| *Taldo* | 432.2 | 345.7 | 139.6 | 271.5 | 172.8 | 196.1 | 203.7 | 189.6 | 182.4 | 284.2 | 210.2 |

### LEGEND

Crystal cells | Lamellocytes | Prohemocytes/Plasmatocytes

#### Bulk RNAseq

#### Single cell RNAseq

SOURCE: https://www.flyrnai.org/tools/single_cell/web/

Tattikota SG, Cho B, Liu Y, Hu Y, Barrera V, Steinbaugh MJ, Yoon SH, Comjean A, Li F, Dervis F, Hung RJ, Nam JW, Ho Sui S, Shim J, Perrimon N. A single-cell survey of *Drosophila* blood. Elife. 2020 May 12;9:e54818. doi: 10.7554/eLife.54818.

#### Single cell RNAseq

48 hours wasp infected

Source: https://www.flyrnai.org/scRNA/blood/

Tattikota SG, Cho B, Liu Y, Hu Y, Barrera V, Steinbaugh MJ, Yoon SH, Comjean A, Li F, Dervis F, Hung RJ, Nam JW, Ho Sui S, Shim J, Perrimon N. A single-cell survey of *Drosophila* blood. Elife. 2020 May 12;9:e54818. doi: 10.7554/eLife.54818.

log normalized UMI: <=0 | 0 | 1 | 2 | 4

#### Single cell RNAseq

SOURCE: https://www.flyrnai.org/tools/single_cell/web/

Cattenoz PB, Sakr R, Pavlidaki A, Delaporte C, Riba A, Molina N, Hariharan N, Mukherjee T, Giangrande A. Temporal specificity and heterogeneity of Drosophila immune cells. EMBO J. 2020 Jun 17;39(12):e104486. doi: 10.15252/embj.2020104486.

#### Hml

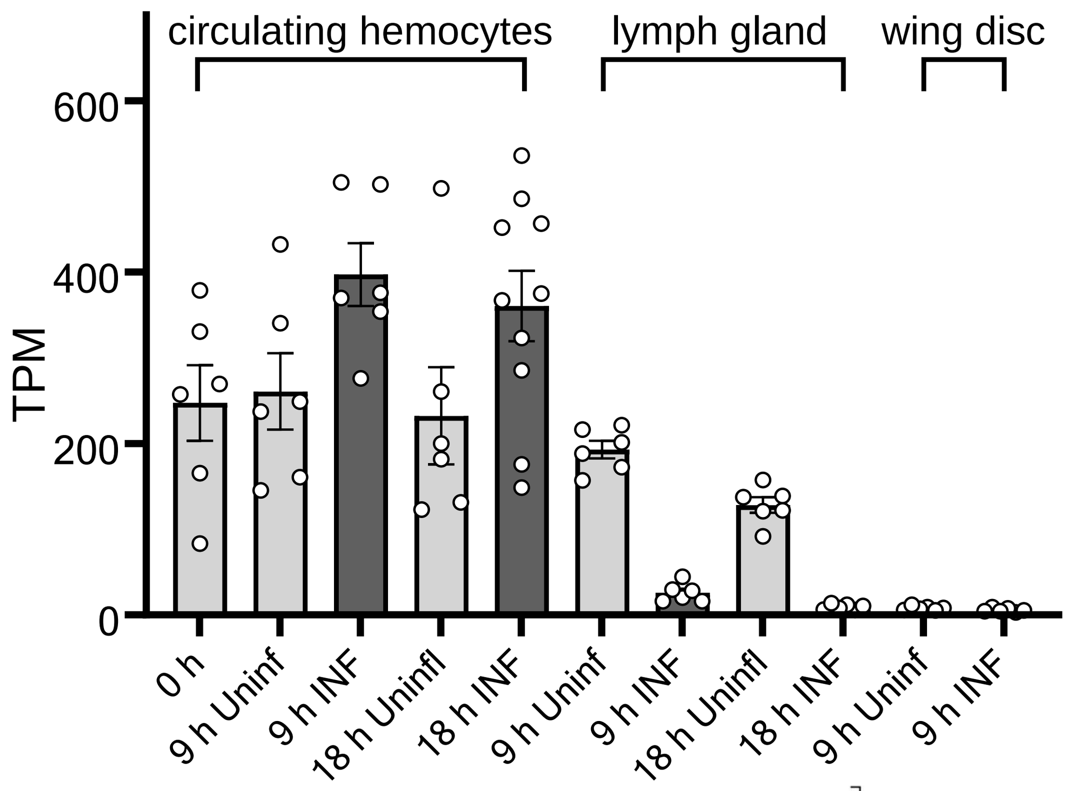


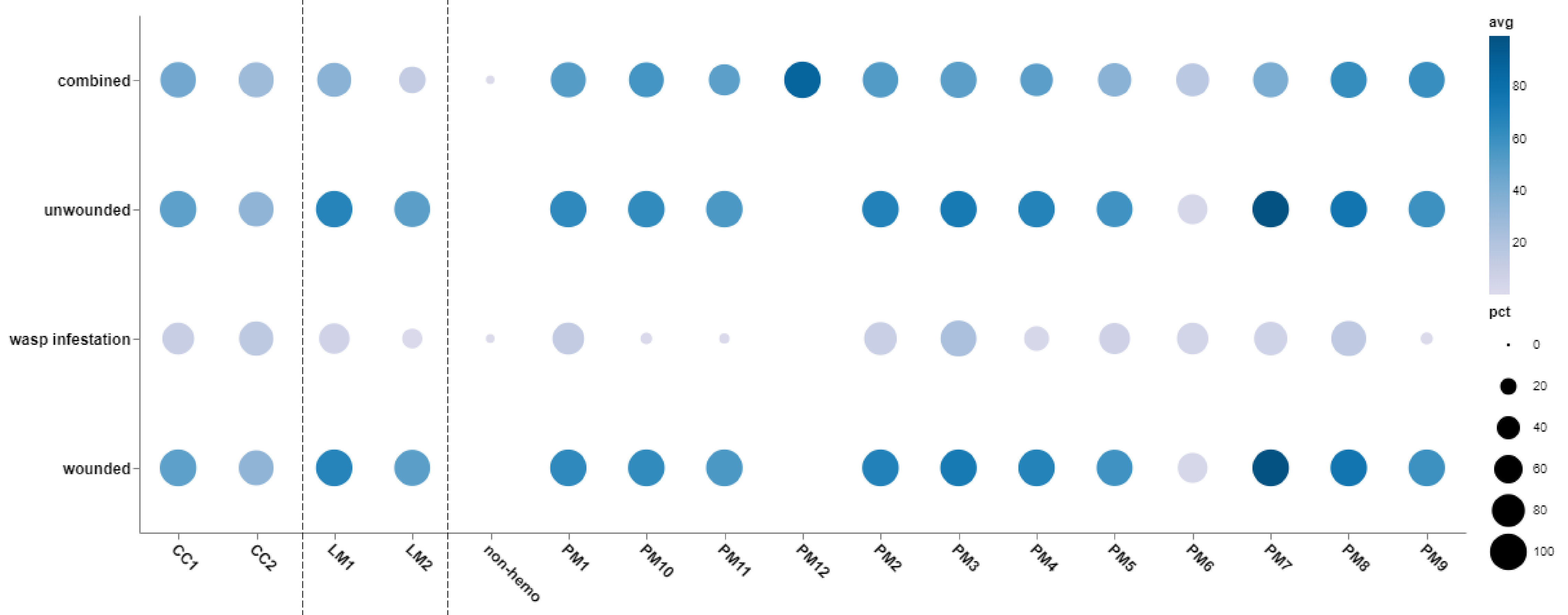


Plasmatocyte marker

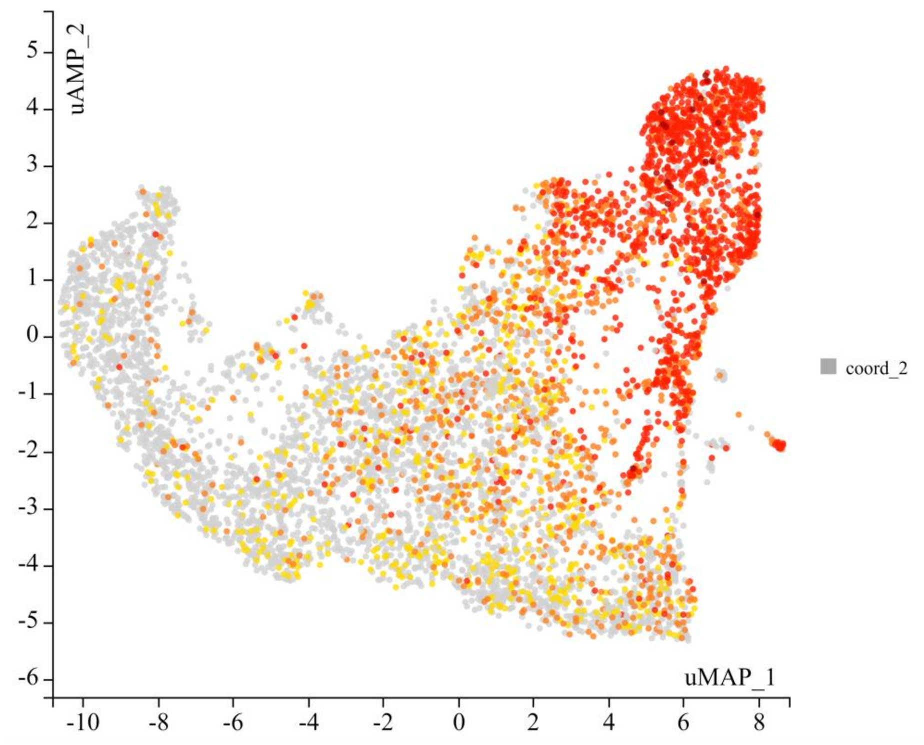


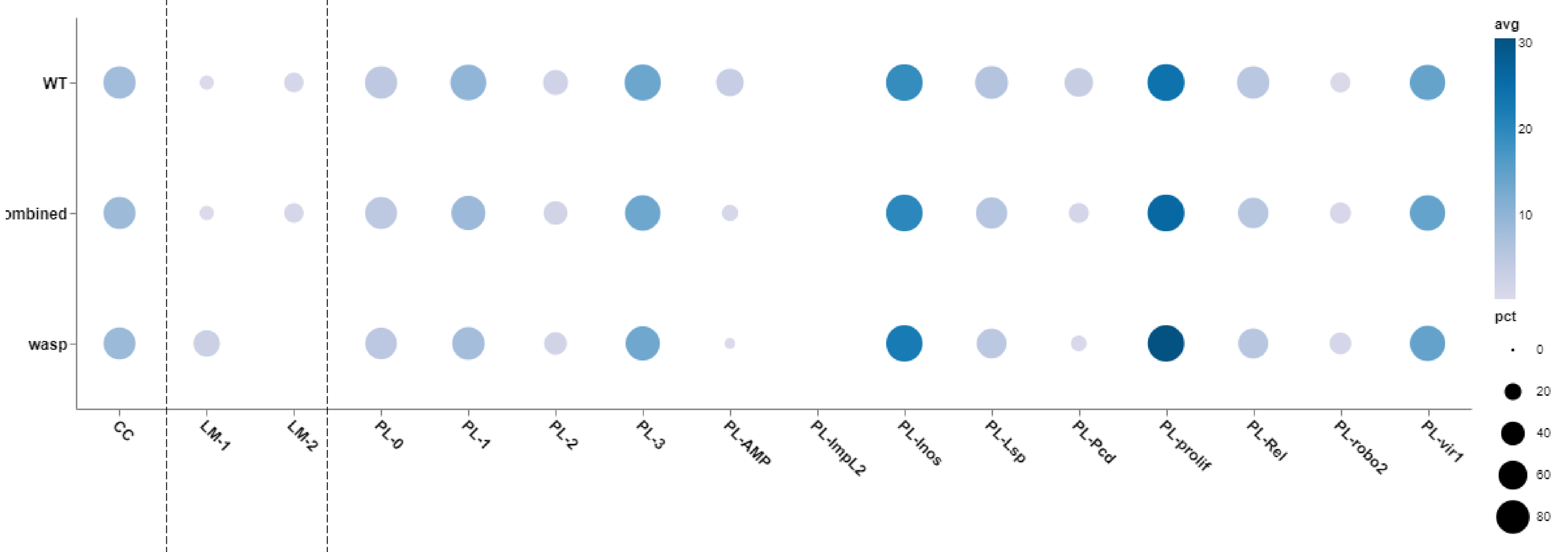


#### Atilla

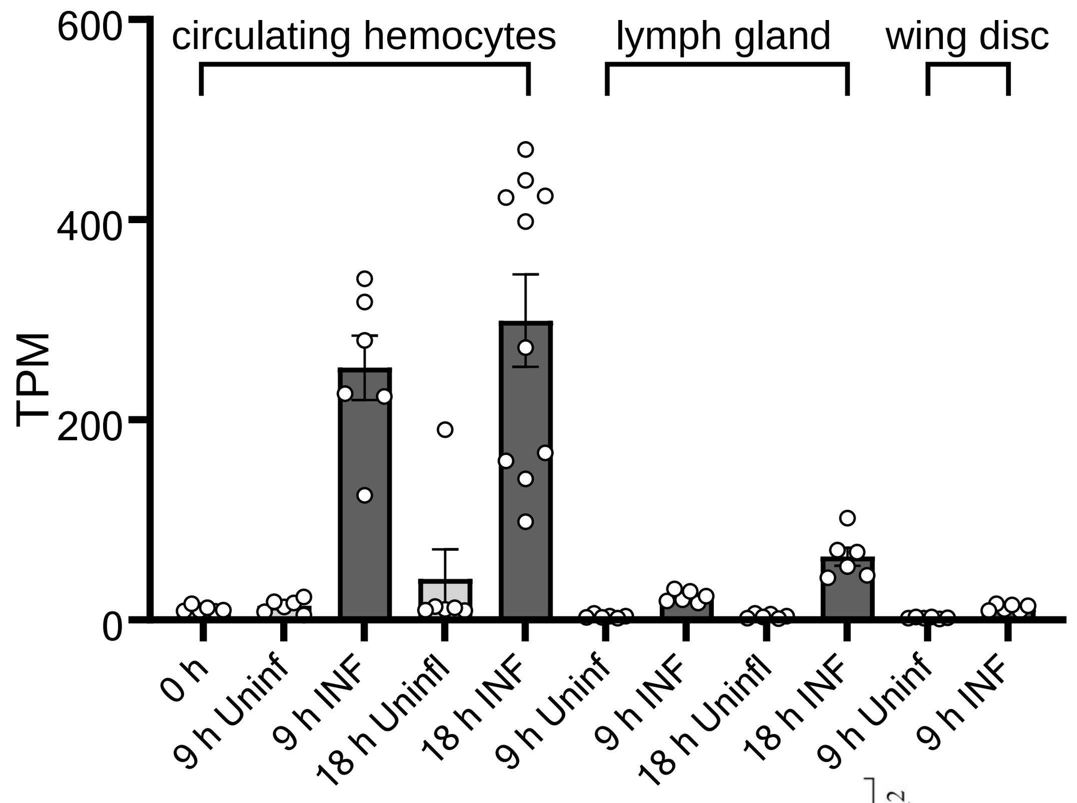


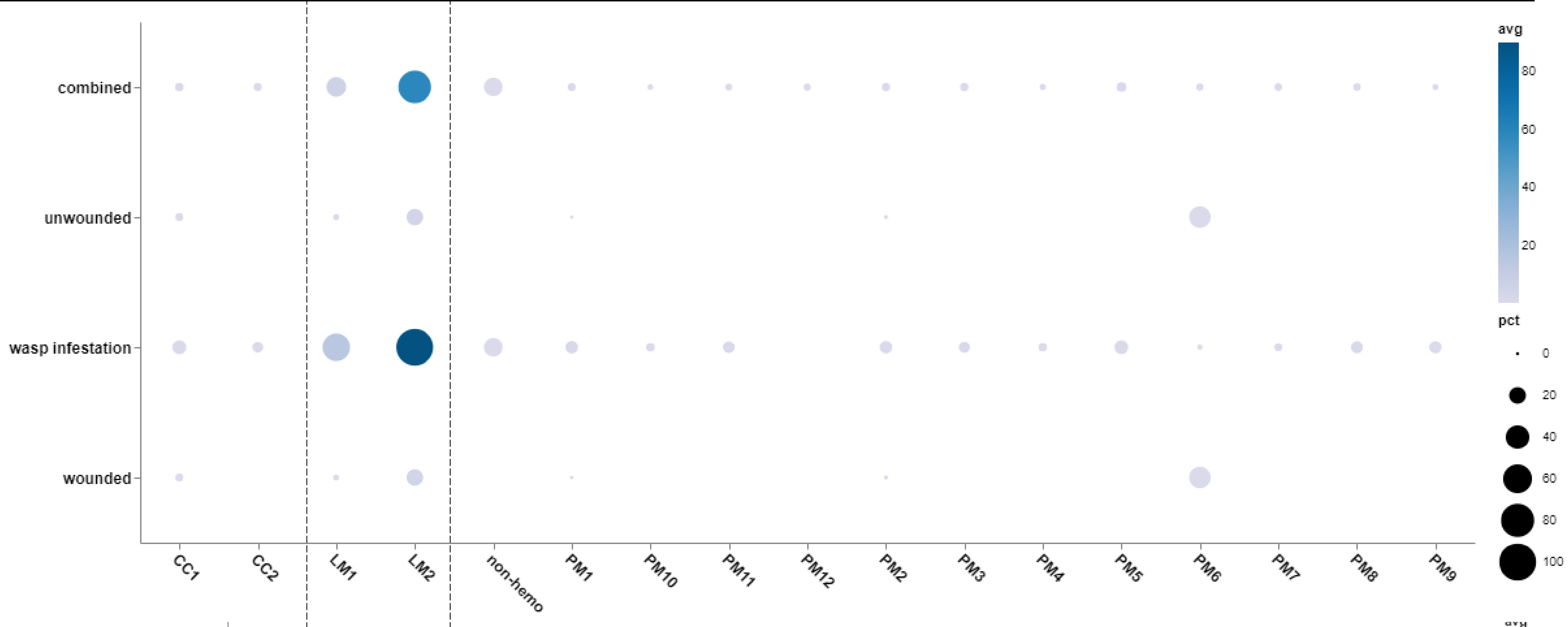


Lamellocyte marker

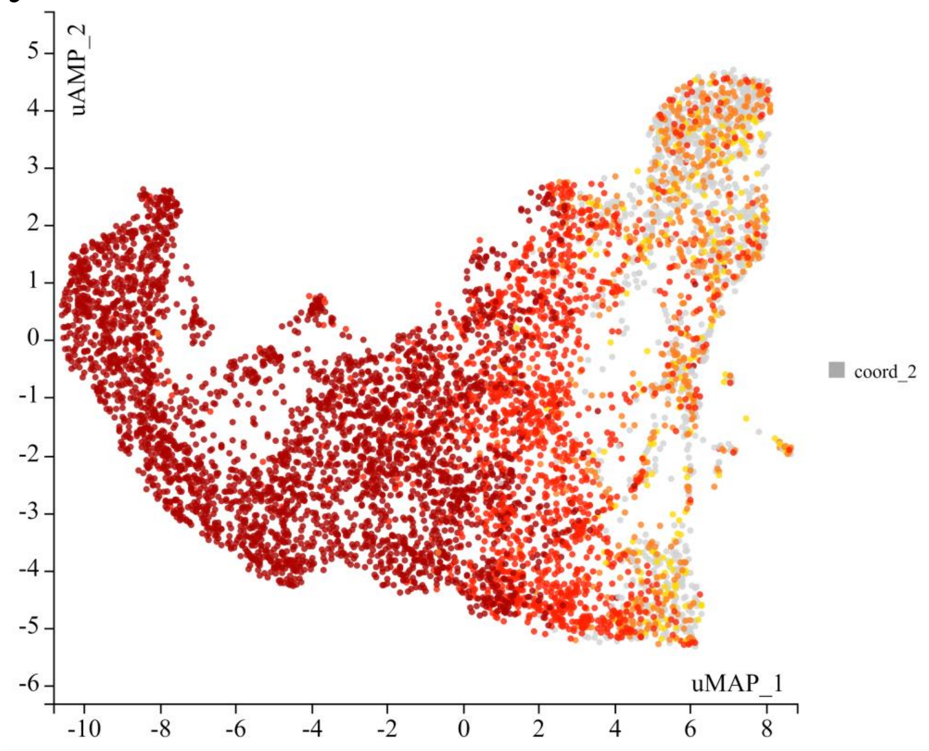


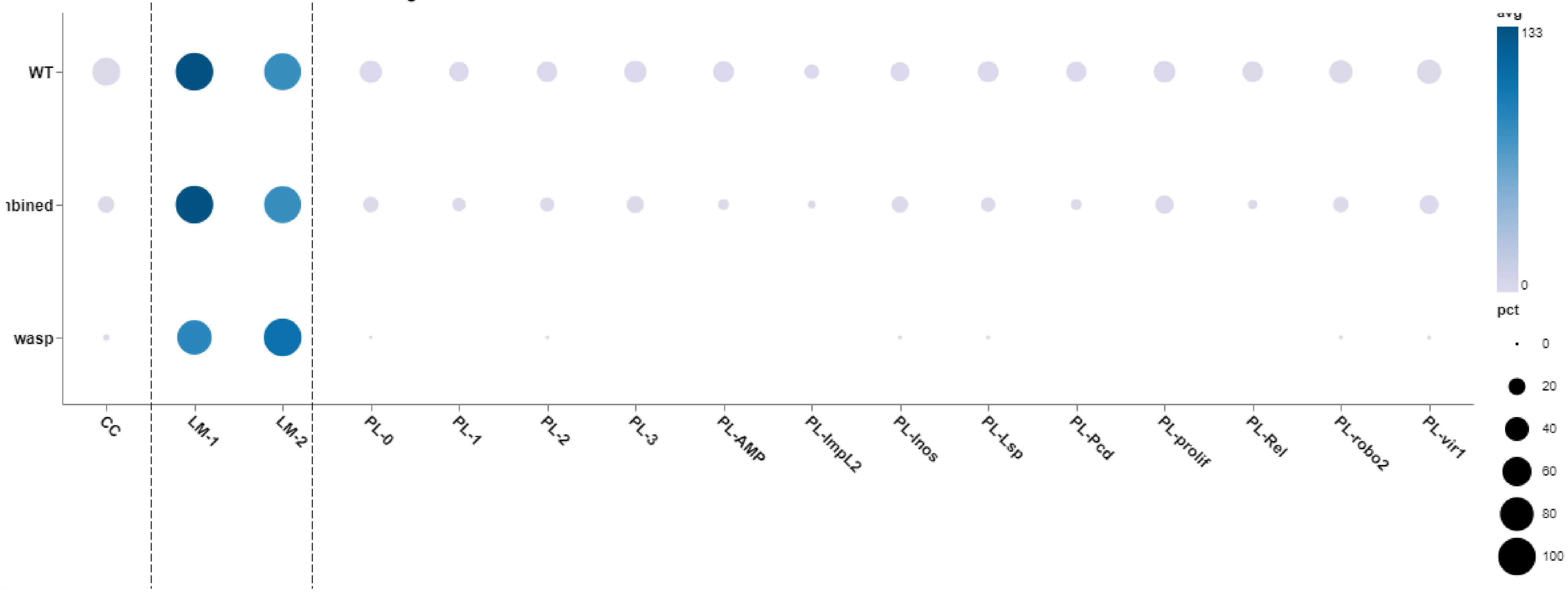

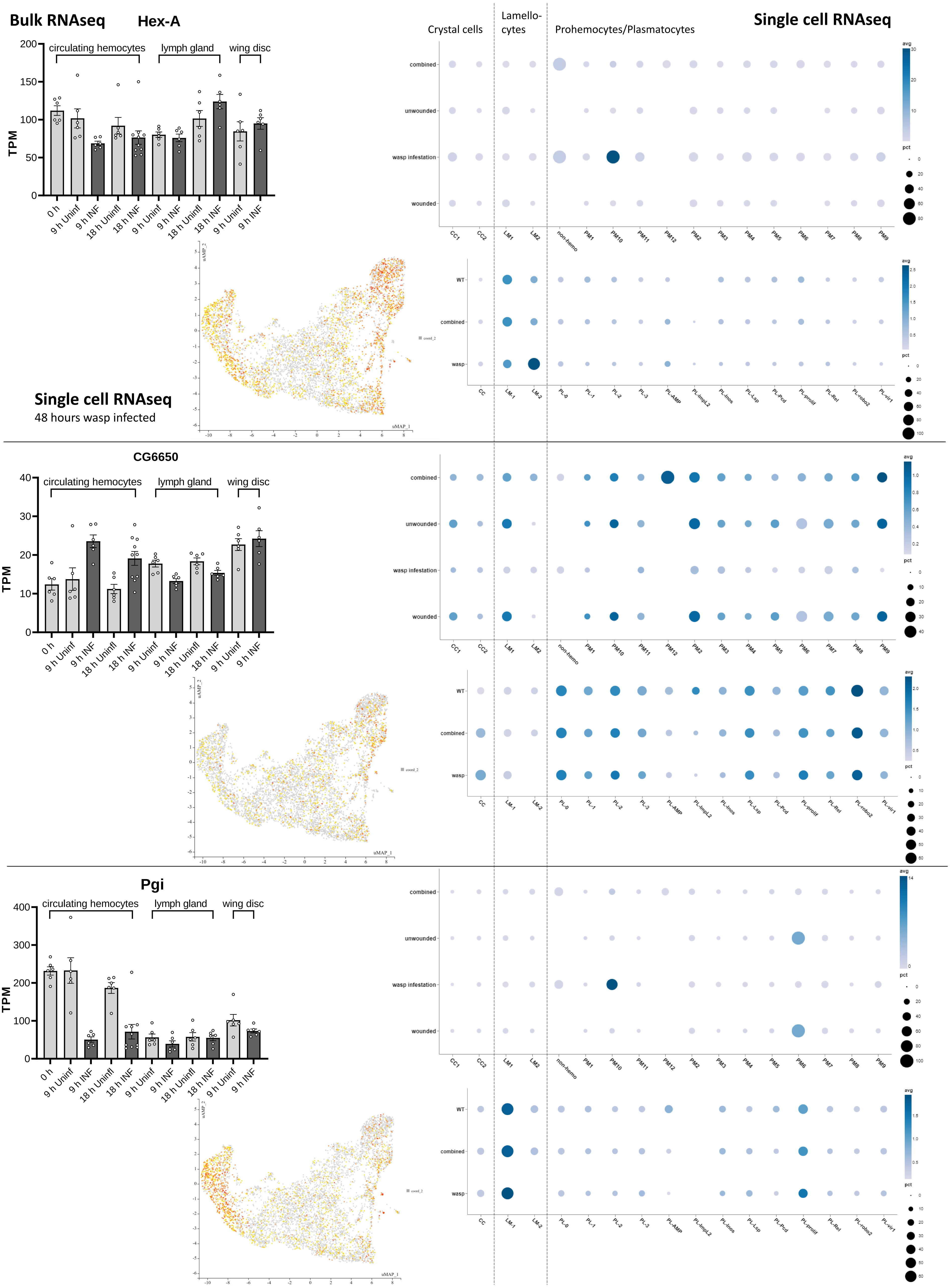

Bulk RNAseq
Hex-A
Single cell RNAseq
Crystal cells
Lamello-cytes
Prohemocytes/Plasmatocytes
circulating hemocytes
lymph gland
wing disc
TPM
0 h
9 h Uninf
9 h INF
18 h Uninfl
18 h INF
combined
unwounded
wasp infestation
wounded
CC1
CC2
LM1
LM2
non-hemo
PM1
PM10
PM11
PM12
PM2
PM3
PM4
PM5
PM6
PM7
PM8
PM9
avg
pct
WT
wasp
CC
LM-1
LM-2
PL-0
PL-1
PL-2
PL-3
PL-AMP
PL-ImpL2
PL-Inos
PL-Lsp
PL-Pcd
PL-prolif
PL-Rel
PL-robo2
PL-vir1
uMAP_1
uMAP_2
Single cell RNAseq
48 hours wasp infected
CG6650
Pgi

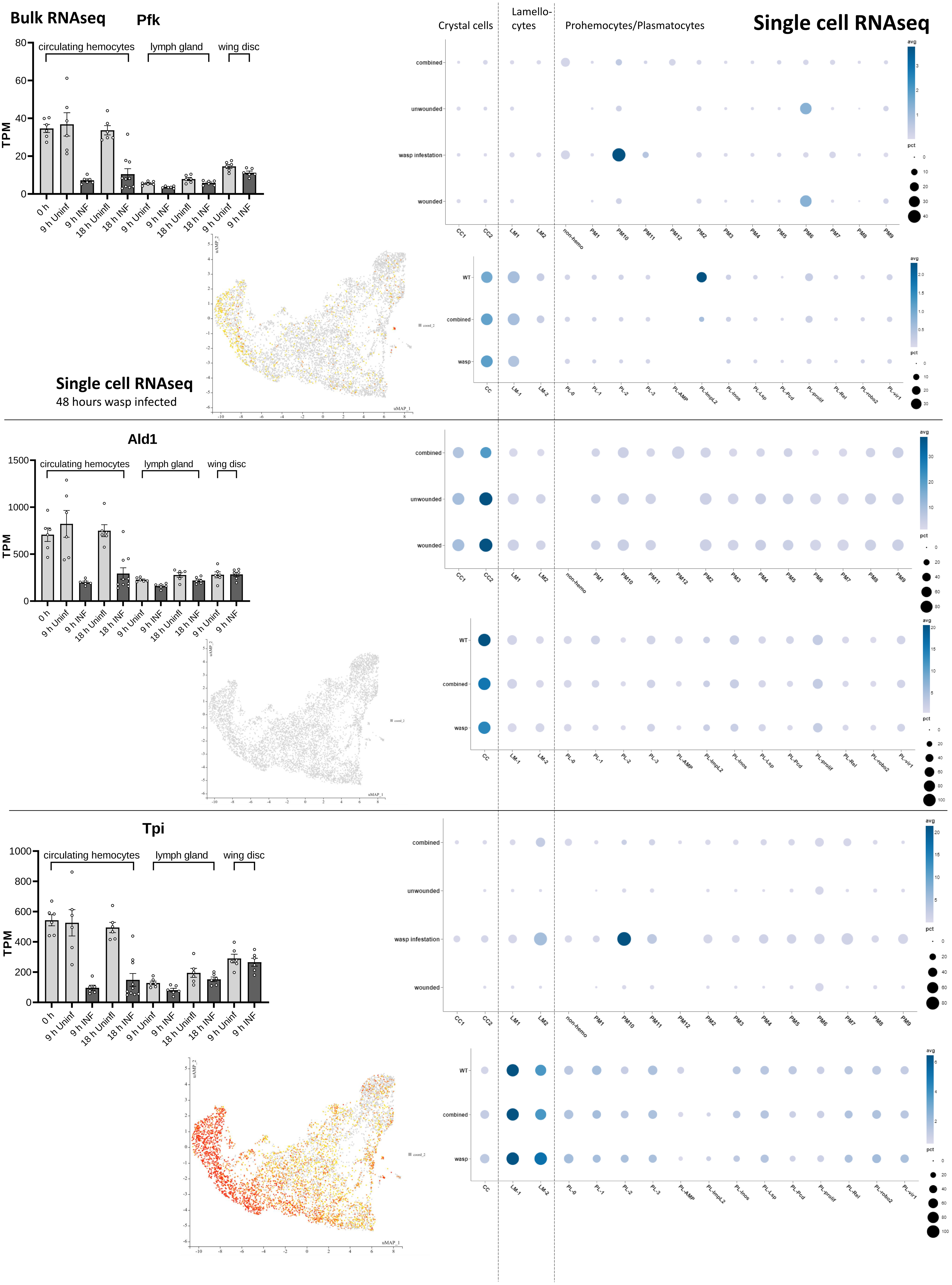
Bulk RNAseq
Pfk
Crystal cells
Lamello-
cytes
Prohemocytes/Plasmatocytes
Single cell RNAseq
circulating hemocytes
lymph gland
wing disc
TPM
0 h
9 h Uninf
9 h INF
18 h Uninfl
18 h INF
9 h Uninf
9 h INF
18 h Uninfl
18 h INF
9 h Uninf
9 h INF
combined
unwounded
wasp infestation
wounded
CC1
CC2
LM1
LM2
non-hemo
PM1
PM10
PM11
PM12
PM2
PM3
PM4
PM5
PM6
PM7
PM8
PM9
avg
pct
WT
combined
wasp
CC
LM-1
LM-2
PL-0
PL-1
PL-2
PL-3
PL-AMP
PL-ImpL2
PL-Inos
PL-Lsp
PL-Pcd
PL-prolif
PL-Rel
PL-robo2
PL-vir1
Single cell RNAseq
48 hours wasp infected
uMAP_1
uMAP_2
coord_2
Ald1
Tpi

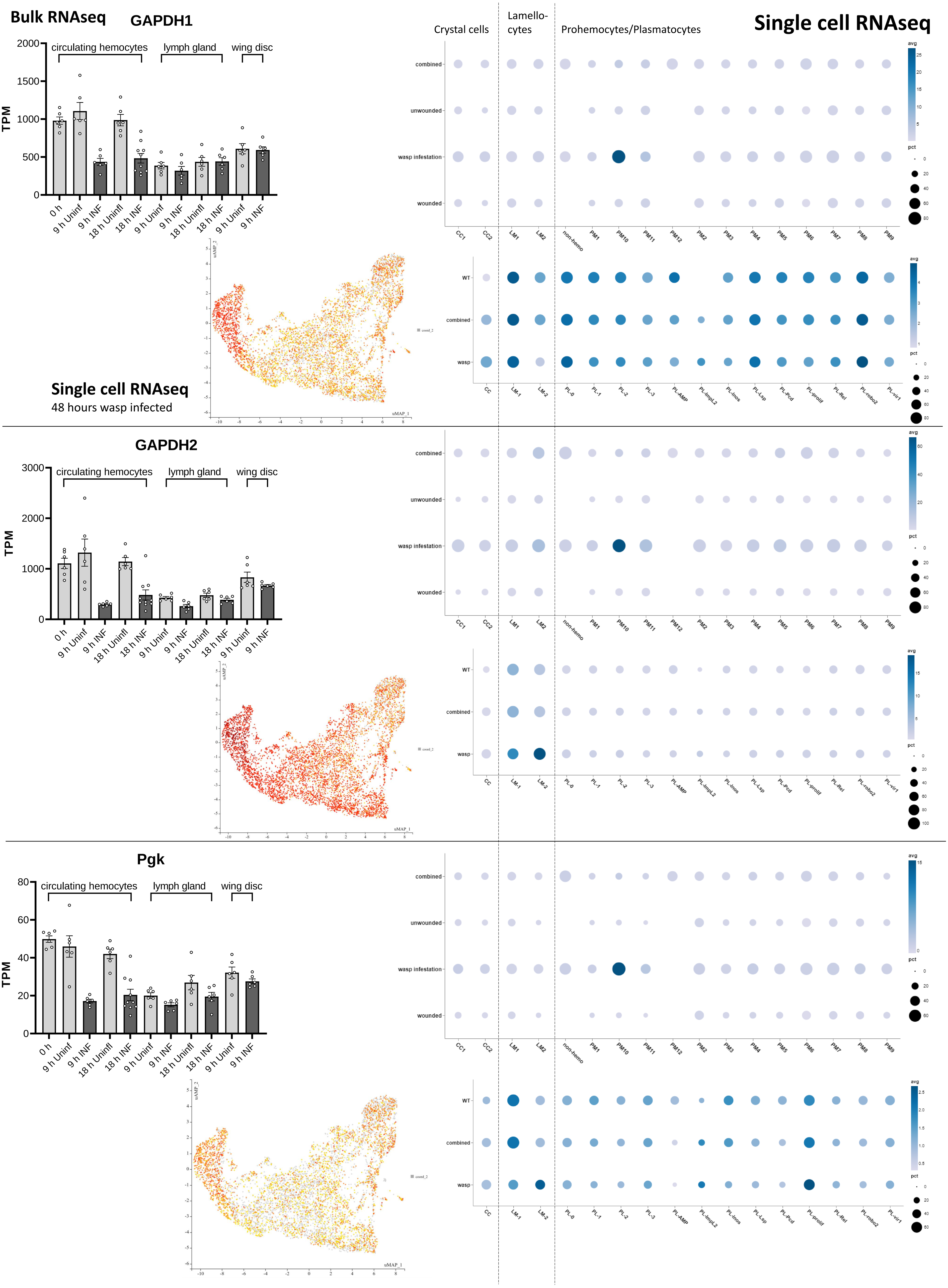

Bulk RNAseq
GAPDH1
Crystal cells
Lamello-cytes
Prohemocytes/Plasmatocytes
Single cell RNAseq
circulating hemocytes
lymph gland
wing disc
TPM
0 h
9 h Uninf
9 h INF
18 h Uninf
18 h INF
combined
unwounded
wasp infestation
wounded
CC1
CC2
LM1
LM2
non-hemo
PM1
PM10
PM11
PM12
PM2
PM3
PM4
PM5
PM6
PM7
PM8
PM9
avg
pct
WT
wasp
CC
LM-1
LM-2
PL-0
PL-1
PL-2
PL-3
PL-AMP
PL-ImpL2
PL-Inos
PL-Lsp
PL-Pcd
PL-prolif
PL-Rel
PL-robo2
PL-vir1
uMAP_1
uMAP_2
Single cell RNAseq
48 hours wasp infected
GAPDH2
Pgk

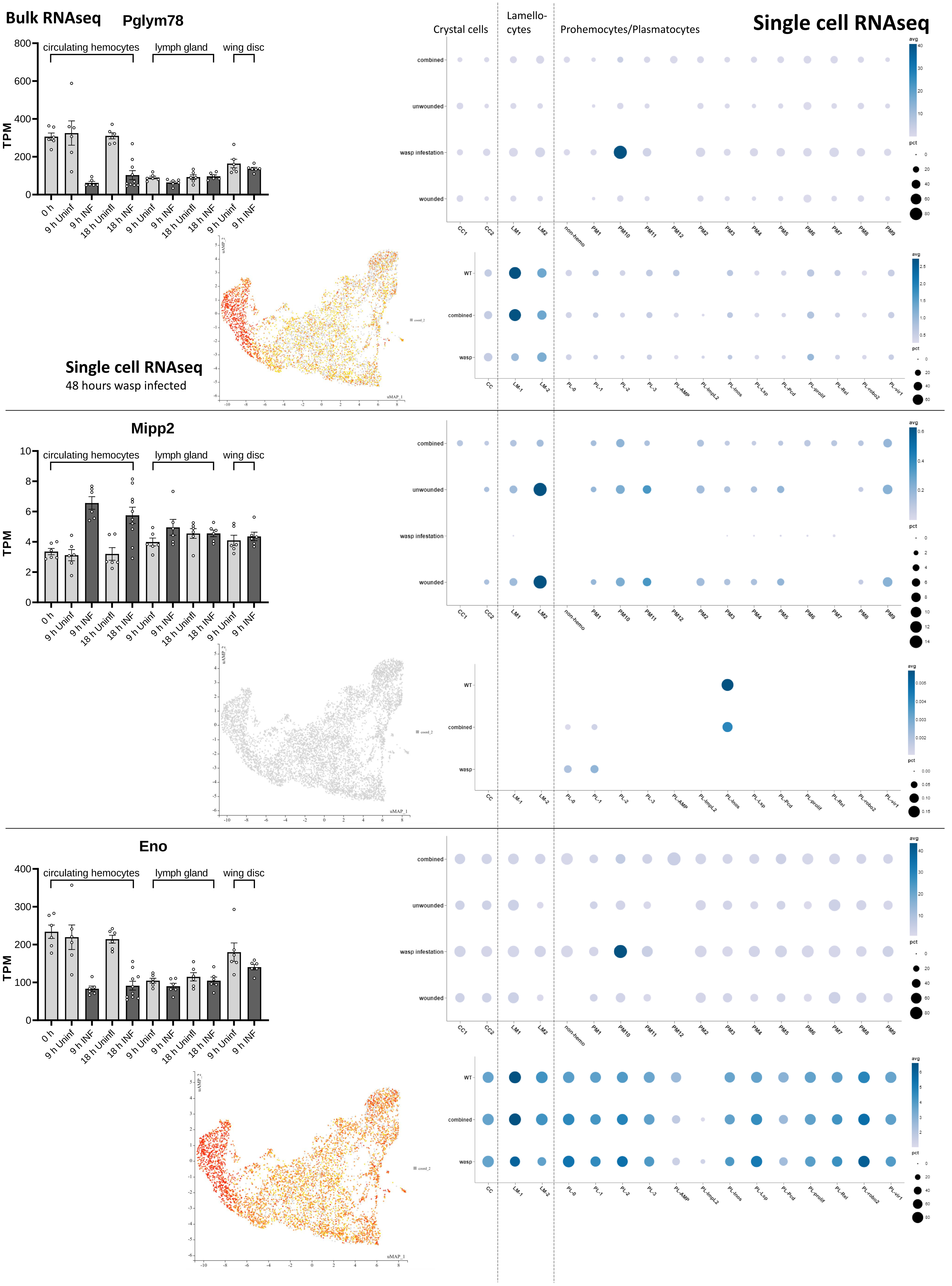

Bulk RNAseq
Pglym78
circulating hemocytes
lymph gland
wing disc
TPM
0 h
9 h Uninf
9 h INF
18 h Uninfl
18 h INF
Crystal cells
Lamello-cytes
Prohemocytes/Plasmatocytes
Single cell RNAseq
combined
unwounded
wasp infestation
wounded
WT
wasp
avg
pct
CC1
CC2
LM1
LM2
non-hemo
PM1
PM10
PM11
PM12
PM2
PM3
PM4
PM5
PM6
PM7
PM8
PM9
CC
LM-1
LM-2
PL-0
PL-1
PL-2
PL-3
PL-AMP
PL-ImpL2
PL-Inos
PL-Lsp
PL-Pcd
PL-prolif
PL-Rel
PL-robo2
PL-vir1
uMAP_1
uMAP_2
coord_2
Single cell RNAseq
48 hours wasp infected
Mipp2
Eno

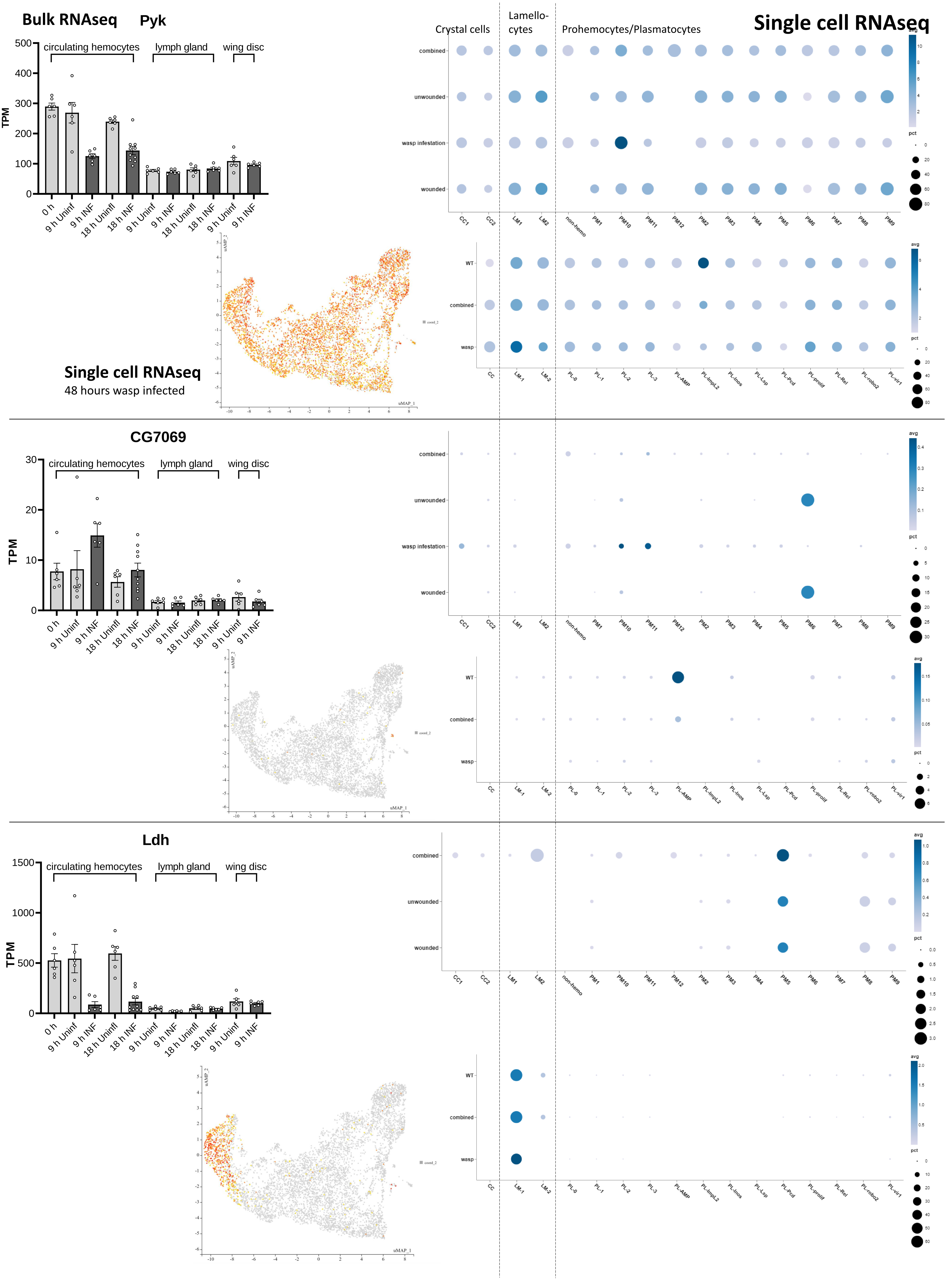

Bulk RNAseq
Pyk
circulating hemocytes
lymph gland
wing disc
TPM
0 h
9 h Uninf
9 h INF
18 h Uninfl
18 h INF
Single cell RNAseq
Crystal cells
Lamello-
cytes
Prohemocytes/Plasmatocytes
combined
unwounded
wasp infestation
wounded
CC1
CC2
LM1
LM2
non-hemo
PM1
PM10
PM11
PM12
PM2
PM3
PM4
PM5
PM6
PM7
PM8
PM9
avg
pct
WT
wasp
CC
LM-1
LM-2
PL-0
PL-1
PL-2
PL-3
PL-AMP
PL-ImpL2
PL-Inos
PL-Lsp
PL-Pcd
PL-prolif
PL-Rel
PL-robo2
PL-vir1
Single cell RNAseq
48 hours wasp infected
uMAP_1
uMAP_2
CG7069
Ldh

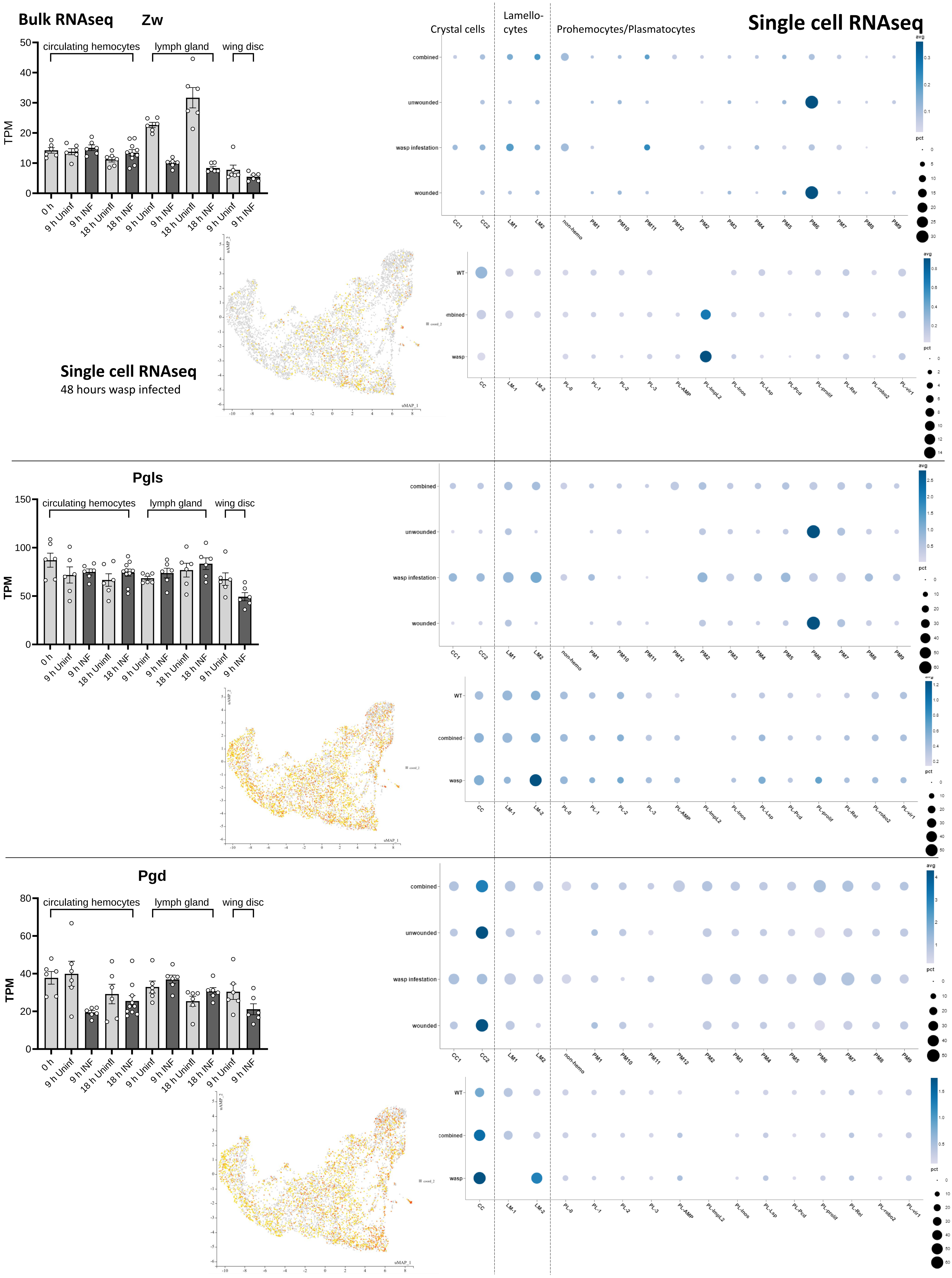

Bulk RNAseq
Zw
Single cell RNAseq
Crystal cells
Lamello-cytes
Prohemocytes/Plasmatocytes
circulating hemocytes
lymph gland
wing disc
TPM
0 h
9 h Uninf
9 h INF
18 h Uninf
18 h INF
combined
unwounded
wasp infestation
wounded
CC1
CC2
LM1
LM2
non-hemo
PM1
PM10
PM11
PM12
PM2
PM3
PM4
PM5
PM6
PM7
PM8
PM9
avg
pct
WT
wasp
CC
LM-1
LM-2
PL-0
PL-1
PL-2
PL-3
PL-AMP
PL-ImpL2
PL-Inos
PL-Lsp
PL-Pcd
PL-prolif
PL-Rel
PL-robo2
PL-vir1
Single cell RNAseq
48 hours wasp infected
uMAP_1
uMAP_2
Pgls
Pgd

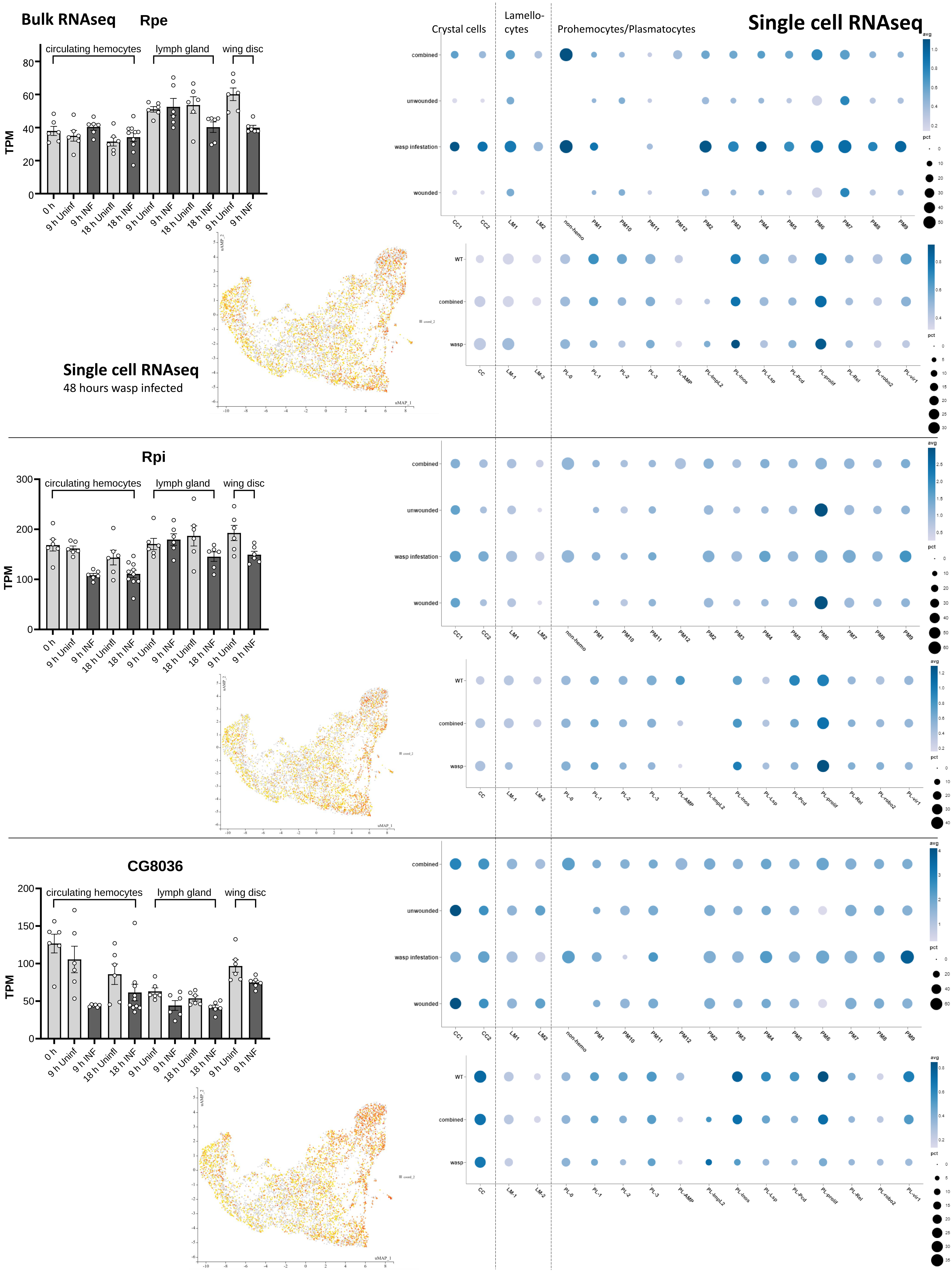
Bulk RNAseq
Rpe
circulating hemocytes
lymph gland
wing disc
TPM
0 h
9 h Uninf
9 h INF
18 h Uninfl
18 h INF
Single cell RNAseq
48 hours wasp infected
Crystal cells
Lamello-cytes
Prohemocytes/Plasmatocytes
combined
unwounded
wasp infestation
wounded
CC1
CC2
LM1
LM2
non-hemo
PM1
PM10
PM11
PM12
PM2
PM3
PM4
PM5
PM6
PM7
PM8
PM9
WT
wasp
CC
LM-1
LM-2
PL-0
PL-1
PL-2
PL-3
PL-AMP
PL-ImpL2
PL-Inos
PL-Lsp
PL-Pcd
PL-prolif
PL-Rel
PL-robo2
PL-vir1
avg
pct
uMAP_1
uMAP_2
Rpi
CG8036

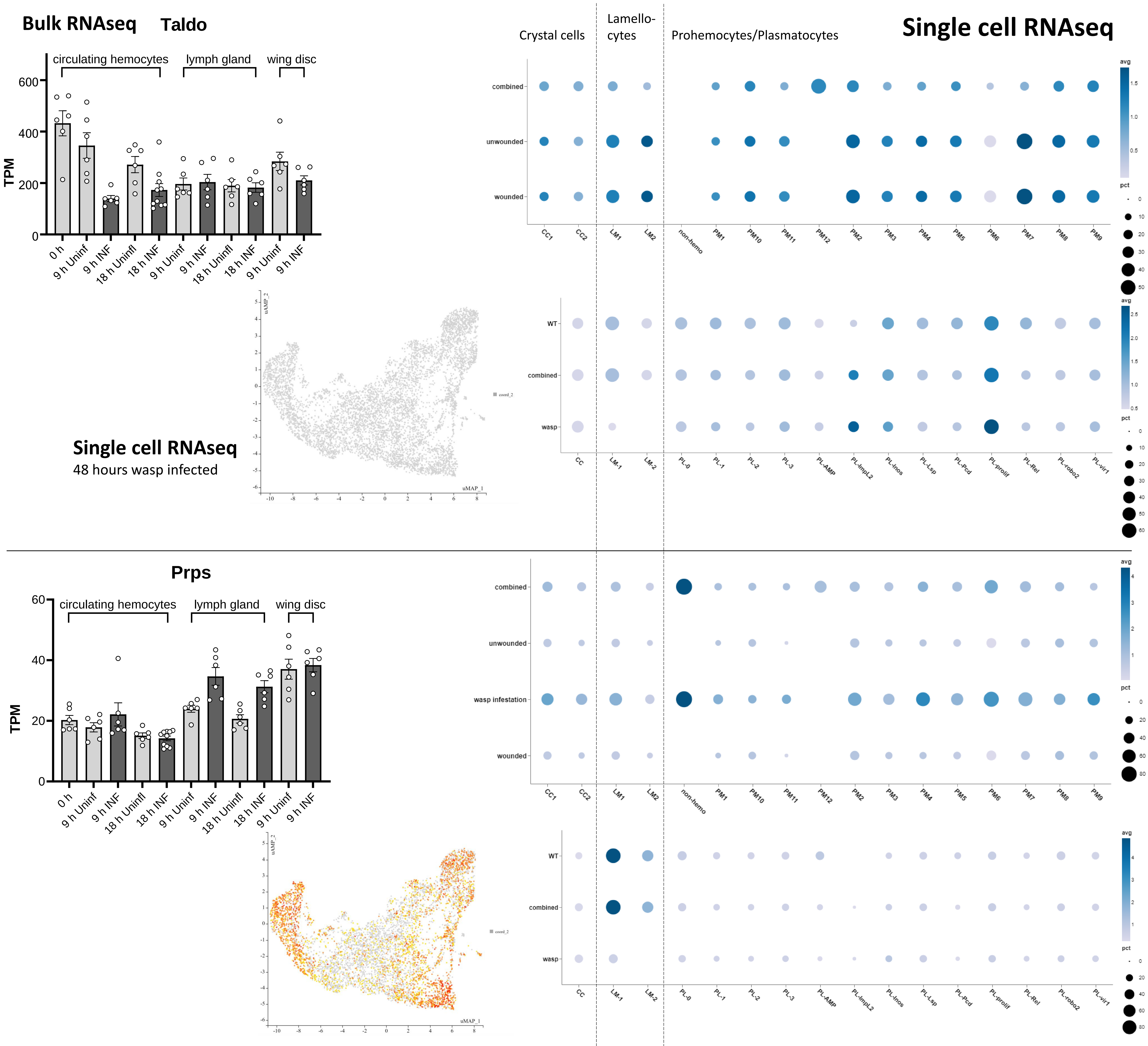

Bulk RNAseq
Taldo
Crystal cells
Lamello-cytes
Prohemocytes/Plasmatocytes
Single cell RNAseq
circulating hemocytes
lymph gland
wing disc
TPM
0 h
9 h Uninf
9 h INF
18 h Uninfl
18 h INF
9 h Uninf
9 h INF
18 h Uninfl
18 h INF
9 h Uninf
9 h INF
combined
unwounded
wounded
CC1
CC2
LM1
LM2
non-hemo
PM1
PM10
PM11
PM12
PM2
PM3
PM4
PM5
PM6
PM7
PM8
PM9
avg
pct
Single cell RNAseq
48 hours wasp infected
uMAP_1
uMAP_2
WT
combined
wasp
CC
LM-1
LM-2
PL-0
PL-1
PL-2
PL-3
PL-AMP
PL-ImpL2
PL-Inos
PL-Lsp
PL-Pcd
PL-prolif
PL-Rel
PL-robo2
PL-vir1
Prps
circulating hemocytes
lymph gland
wing disc
TPM
combined
unwounded
wasp infestation
wounded
